# Stereochemical Insights into Sarpagan and Akuammiline Alkaloid Biosynthesis

**DOI:** 10.1101/2024.12.19.629490

**Authors:** Scott Galeung Alexander Mann, Melina Paz-Galeano, Mohammadamin Shahsavarani, Jacob Owen Perley, Jun Guo, Jorge Jonathan Oswaldo Garza-Garcia, Yang Qu

## Abstract

The Apocynaceae family produces a diverse array of monoterpenoid indole alkaloids (MIAs) with significant pharmaceutical value. Among these, sarpagan and akuammiline alkaloids stand out for their complex stereochemistry, derived from the enzymatic cyclization and rearrangement of geissoschizine. This study investigates the stereochemical outcomes of sarpagan bridge enzymes (SBEs) and rhazimal synthases (RHS), key players in geissoschizine cyclization and MIA diversification. Using two known and five newly identified enzymes from six plant species, we show that RHS enzymes from *Alstonia scholaris*, *Vinca minor*, and *Amsonia tabernaemontana* exclusively produce the 16*R* rhazimal stereoisomer. Meanwhile, SBEs from *Catharanthus roseus*, *Tabernaemontana elegans*, *Vinca minor*, and *Rauvolfia serpentina* likely generate 16*R* polyneuridine aldehyde; however, downstream aldehyde reductase, deformylase, and esterase activities further epimerize and alter the C16 stereochemistry, yielding naturally occurring alkaloids with distinct C16 stereochemistry across species. These findings, supported by in vitro assays and in planta silencing of *C. roseus* CrSBE after we reroute biosynthetic flux toward mutated sarpagan MIAs, further reveal enzymatic control over C16 stereochemistry in sarpagan MIA biosynthesis. By elucidating the transformation of diastereomeric intermediates, this work provides key insights into the stereochemical and enzymatic diversification of MIAs in nature.

## Introduction

The Apocynaceae family is renowned for its exceptional diversity of monoterpenoid indole alkaloids (MIAs), a class of structurally complex and pharmaceutically significant natural products. Clinically important drugs, such as the anticancer drugs vinblastine and vincristine, exemplify the therapeutic value of MIAs (Luca *et al*., 2012). These compounds are categorized based on their carbon skeleton arrangements, which arise through distinct biosynthetic pathways. Among the diverse MIA subclasses, sarpagan and akuammiline alkaloids stand out due to their intricate stereochemical frameworks and pharmaceutical relevance. Notable examples include the antiarrhythmic drug ajmaline (sarpagan) and the natural analgesic akuammine (akuammiline) (Menzies *et al*., 1998; Guo *et al*., 2024). Both alkaloid classes originate from the central precursor strictosidine (Fig. 1), undergoing enzymatically regulated cyclization, rearrangement, and modification steps to establish their defining chiral centers. Understanding these stereochemical pathways is vital, as the biological activities of these compounds often hinge on their precise three-dimensional configurations.

**Figure 1.**
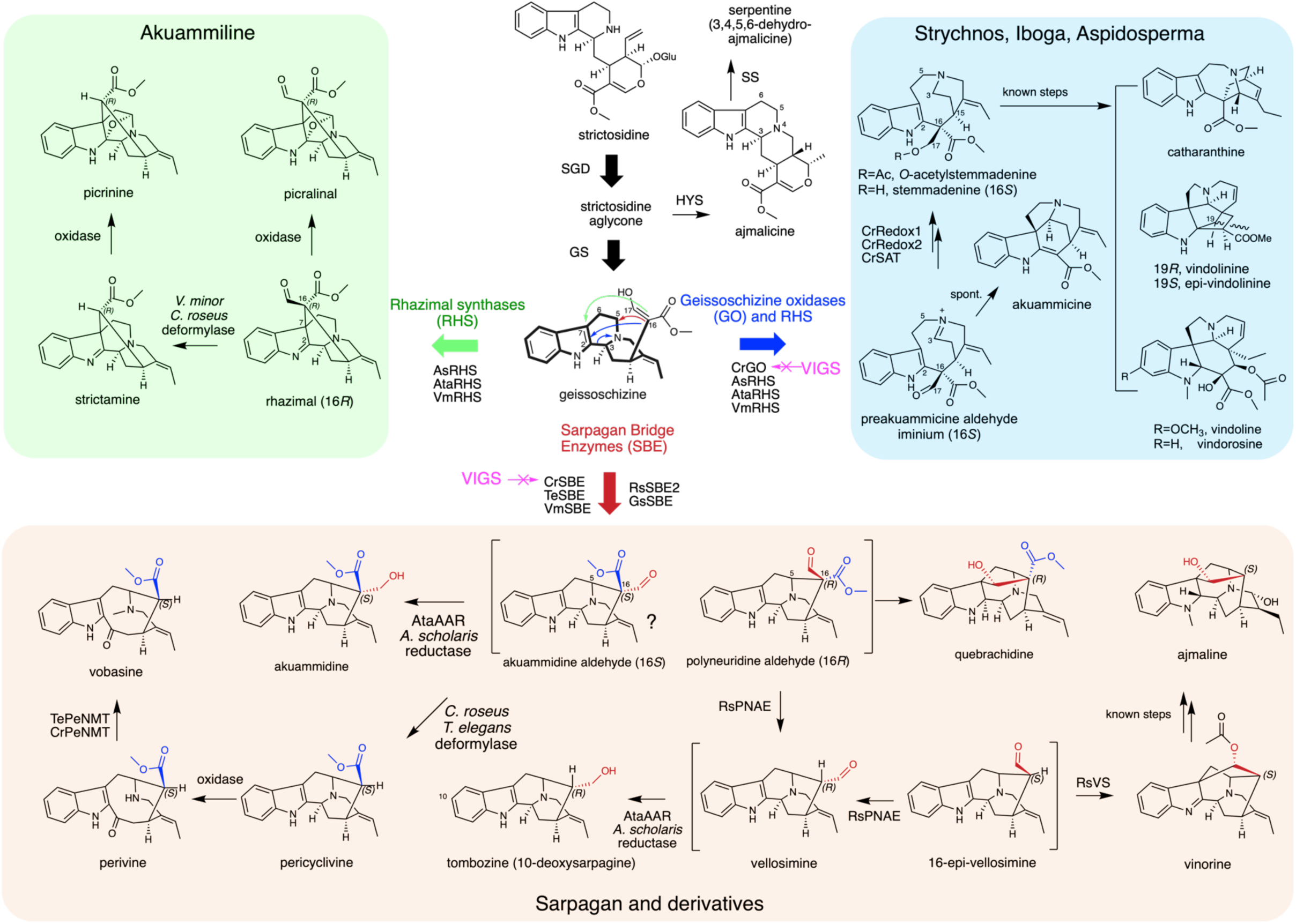
The biosynthetic framework of monoterpenoid indole alkaloids (MIAs) for akuammiline, sarpagan, strychnos, iboga and aspidosperma types. 19*E*-geissoschizine cyclizations by homologous cytochrome P450 monooxygenases (CYPs) play a pivotal role in producing these major MIA skeletons.

Upon deglycosylation, strictosidine generates a series of reactive, labile intermediates collectively known as strictosidine aglycone, which exist in equilibrium. Various cinnamyl alcohol dehydrogenase (CAD)-like reductases selectively reduce the iminium intermediates, leading to the formation of three major carbon skeletons: corynanthe, yohimbine, and heteroyohimbine with multiple stereochemical variations (Stavrinides *et al*., 2016; Qu *et al*., 2017, 2018a; Kim *et al*., 2023; Stander *et al*., 2023). Among these, 19*E*-geissoschizine (Fig. 1) serves as a pivotal intermediate for the biosynthesis of three major MIA skeletons: strychnos, akuammiline, and sarpagan (Scott, 1974; Qu *et al*., 2017). These skeletons differ in the site of C– C bond formation: C2–C16 for strychnos, C7–C16 for akuammiline, and C5–C16 for sarpagan, initiated by *N*4-oxidation and cyclization via the reactive 17-enol (Fig. 1). These reactions are catalyzed by homologous cytochrome P450 monooxygenases (CYPs), namely geissoschizine oxidase (GO) from *Catharanthus roseus*, rhazimal synthase (RHS) from *Alstonia scholaris*, and sarpagan bridge enzyme (SBE) from *Rauvolfia serpentina*, all of which have been cloned and functionally characterized in recent years (Dang *et al*., 2018; Qu *et al*., 2018a; Wang *et al*., 2022; Guo *et al*., 2024). Despite these advances, a significant gap remains regarding the stereochemical fate of C16 during the formation of these key bridges, particularly for the sarpagan and akuammiline skeletons.

The strychnos alkaloids, including akuammicine and norfluorocurarine, represent a smaller subset of MIAs where the original C16 stereochemistry from GO activity is eliminated during downstream transformations. For instance, spontaneous deformylation of GO product preakuammicine aldehyde iminium yields akuammicine (Fig. 1), while esterase-mediated decarboxylation generates norfluorocurarine, both intermediates for other MIA subclasses (Qu *et al*., 2018a; Hong *et al*., 2022). In contrast, tandem iminium and aldehyde reductions by CrRedox1 and CrRedox2 preserve the C16 stereochemistry in critical intermediates stemmadenine and *O*-acetylstemmadenine (Fig. 1) (Qu *et al*., 2018a). However, reductase-mediated deacetylation induces C15–C16 bond cleavage, forming a C16–C17 alkene that again eliminates the original C16 stereochemistry (Benayad *et al*., 2016; Qu *et al*., 2018a,b; Caputi *et al*., 2018; Jarret *et al*., 2019; Eng *et al*., 2022). The resulting dehydrosecodine serves as an intermediate to all iboga, aspidosperma, and pseudoaspidosperma alkaloids (Qu *et al*., 2015, 2018a,b; Caputi *et al*., 2018; Williams *et al*., 2019; Eng *et al*., 2022; Kamileen *et al*., 2022; Guo *et al*., 2024), including the key monomers catharanthine and vindoline that undergo coupling to form the clinically important anticancer drug vinblastine in *C. roseus* (Fig. 1).

Geissoschizine’s C7–C16 bridging can instead yield rhazinaline (16*S*) in *Rhazya stricta* (Chatterjee *et al*., 2006) and rhazimal (16*R*), along with derivatives like akuammilline, picralinal, and strictamine in *C. roseus*, *A. scholaris*, and *Vinca minor* (Fig. 1) (Kohl *et al*., 1981; Yamauchi *et al*., 1990; Levac *et al*., 2016; Vrabec *et al*., 2022). Similarly, the C5–C16 bridging, known as the sarpagan bridge, can theoretically generate both akuammidine aldehyde (16*S*) and polyneuridine aldehyde (16*R*). Downstream alkaloids may retain the akuammidine aldehyde stereochemistry, as seen in vobasine from *Tabernaemontana elegans* (Farzana *et al*., 2024), pericyclivine in *C. roseus* (Qu *et al*., 2018a), and sarpagine in *R. serpentina* (Bartlett *et al*., 1962). while polyneuridine aldehyde stereochemistry is observed in alkaloids such as ajmaline and quebrachidine (Fig. 1) (Ahamada *et al*., 2016; Turpin *et al*., 2020; Guo *et al*., 2024).

Our previous studies revealed that *C. roseus* GO produces exclusively 16*S* product, confirmed via 1D/2D Nuclear Magnetic Resonance (NMR) studies, including Nuclear Overhauser Effect Spectroscopy (NOESY) (Qu *et al*., 2018a). By contrast, *A. scholaris* AsRHS generates 16*R* stereochemistry, as evidenced by its alcohol derivative rhazimol, verified against synthetic standards (Zhang *et al*., 2019; Wang *et al*., 2022). However, the stereochemistry of SBE products remains experimentally uncharacterized, with conclusions thus far inferred only from downstream products.

In this study, we systematically investigate the products and stereochemical outcomes of SBE and RHS enzymatic reactions. By analyzing two previously characterized enzymes alongside five newly identified enzymes from six Apocynaceae species, we demonstrate that RHS enzymes from *A. scholaris*, *V. minor*, and *Amsonia tabernaemontana* exclusively produce the 16*R* rhazimal stereoisomer. We also show that SBEs products from *C. roseus*, *T. elegans*, *V. minor*, and *R. serpentina* can be epimerized by downstream reductase, deformylases, and esterase into alkaloids with distinct C16-stereochemistry, such as ajmaline (polyneuridine stereochemistry) and pericyclivine/vobasine (akuammidine stereochemistry). Our findings are further supported by *in planta* silencing of a new enzyme *C. roseus* CrSBE, coupled with metabolically redirecting the biosynthetic flux to the otherwise muted sarpagan pathway in this plant. Our results update and advance the understanding of sarpagan MIA C16-stereoisomer formation, addressing longstanding questions about the stereochemical dynamics that drive MIA biosynthesis and diversification.

## Results

### CrSBE is necessary for 16*S*-sarpagan alkaloids biosynthesis in *C. roseus*

When we previously discovered and characterized CrGO, we examined the effects of silencing the corresponding gene in *C. roseus* leaves. CrGO silencing caused significant reduction of vindoline and catharanthine, downstream of the GO-product preakuammicine aldehyde iminium (Fig. 1) (Qu *et al*., 2018a). We also observed a dramatic increase of pericyclivine and perivine, which typically accumulate at low levels (Qu *et al*., 2018a). These findings indicated a suppressed flux toward 16*S*-sarpagan MIAs due to strong CrGO activity in wild type *C. roseus*.

The *C. roseus* leaf epidermis is the most important tissue for MIA formation and diversification (Kulagina *et al*., 2022). After analyzing a *C. roseus* leaf epidermal transcriptome (Murata *et al*., 2008), we identified an epidermis-specific GO homolog, which we designated as *C. roseus* Sarpagan Bridge Enzyme (CrSBE, Supplementary table 1) based on subsequent experimental validation. The Expressed Sequence Tag (EST) counts, a proxy for gene expression, were 38 and 8 for CrGO and CrSBE in epidermis, respectively. The amino acid sequence of CrSBE was 51% identical to CrGO and 77% identical to *R. serpentina* RsSBE2 (Guo *et al*., 2024). No additional GO or SBE homologs were detected in this transcriptome generated from laser-dissected leaf epidermis. We hypothesized that CrSBE was responsible for 16*S*-sarpagan alkaloid biosynthesis in *C. roseus* and conducted virus-induced gene silencing (VIGS) experiments to investigate its function *in planta*.

Silencing CrSBE transcripts did not affect the major leaf alkaloids, such as vindoline and catharanthine; however, the low-abundance sarpagan MIA pericyclivine was notably reduced to 15.0% when CrSBE transcript levels dropped to 12.3% (Supplementary fig. 1). Given the low natural abundance of sarpagan alkaloids in *C. roseus*, we reasoned that co-silencing CrGO could enhance their accumulation and better demonstrate the physiological relevance of CrSBE. To this end, we silenced CrGO individually and also employed a single chimeric VIGS construct to achieve simultaneous silencing of CrGO and CrSBE.

Silencing CrGO transcripts by 89.6% resulted in 89.2% and 82.6% reduction of the two major alkaloids catharanthine and vindoline/vindorosine (demethoxyvindoline) in *C. roseus*, respectively (Fig.1, Fig. 2A-C), consistent with our previous findings. Similarly, vindolinine and epi-vindolinine, the third most abundant MIAs differing at C19 stereochemistry, also decreased by 82.7%. In contrast, ajmalicine and serpentine levels, which form prior to geissoschizine cyclization (Fig. 1), remained unchanged. In response to significantly reduced GO levels, 19*E*-geissoschizine accumulated to 1.45 mg/g fresh leaf, which was otherwise below detection limit in empty vector controls (Figs. 2B, C). In this background, pericyclivine and perivine levels increased significantly, by 9.8-fold and 4.1-fold, respectively, as free geissoschizine became available to CrSBE. Co-silencing GO and SBE by 64.5% and 95.9% (Fig. 2A), respectively, led to 97.5% and 92.2% reductions in pericyclivine and perivine, alongside reductions in major MIAs and further accumulation of geissoschizine (Figs. 2A, B). These results indicated that CrSBE is physiologically responsible for the sarpagan alkaloid biosynthesis in *C. roseus*.

**Figure 2.**
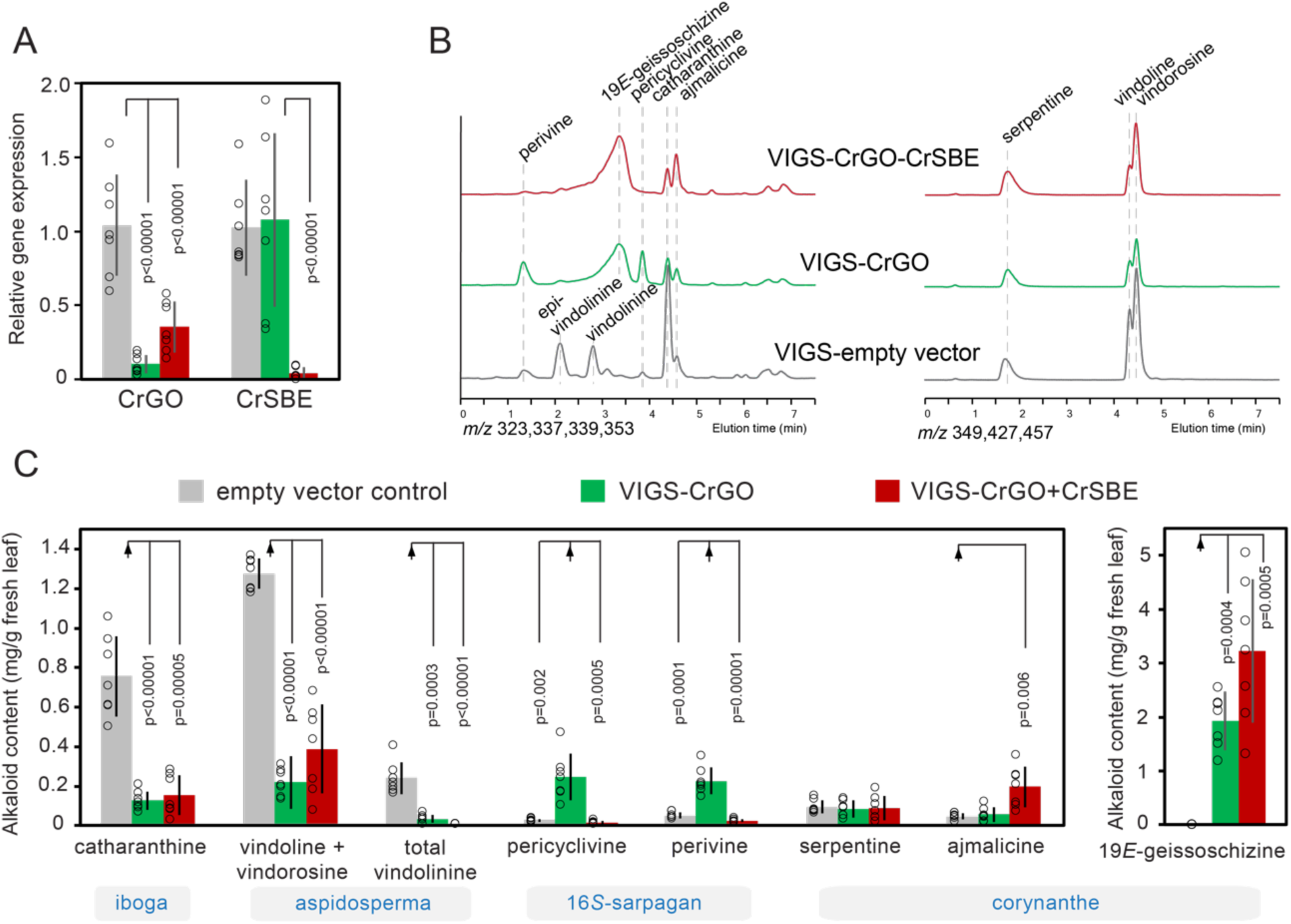
Virus-induced gene silencing (VIGS) of CrSBE significantly reduced sarpagan alkaloid accumulation in *C. roseus* leaf. Co-silencing CrGO redirected metabolic flux to the sarpagan biosynthetaic pathway, enabling proper evaluation of CrSBE’s physiological role. (A) Quantatitive Reverse Transcription Polymerase Chain Reaction (qRT-PCR) analysis of the relative expression levels of CrGO and CrSBE in the empty vector control, VIGS-CrGO, and VIGS-CrGO+CrSBE co-silencing plants. (B) Representative LC-MS chromatograms for selected ions [M+H]^+^ *m/z* 323,337,339,349,353,427,457 show the reduction and increase of alkaloids in control and VIGS plants. (C) Quantification of major alkaloids in control and VIGS plants. Data was generated from seven independent plants per treatment. Individual data points are shown as circles, and error bars represent standard deviation. Statistical analysis was performed using the Student’s *t*-test. The structures of alkaloids are found in Figure 1.

Since polyneuridine aldehyde (16*R*) has been the only hypothetical product of SBE (Pfitzner & Stöckigt, 1983; Dang *et al.,* 2018) and pericyclivine/perivine contain the opposite C16-stereochemistry, we carefully re-analysed the *C. roseus* MIA chemistry. With LC-MS/MS, NOESY and other NMR experiments (Supplementary figs. 2-8), we confirmed the C16-stereochemistry of pericyclivine purified from VIGS-GO plants (Qu *et al*., 2018a). The up-field chemical shift of pericyclivine’s carbomethoxy group (δ 3.02 ppm), caused by indole shielding, is indicative for akuammidine 16*S* stereochemistry (Table 1 and 2). In NOESY spectra, the H16, H5, and H15 cross peaks were also consistent with such stereochemistry. For perivine, we confirmed its stereochemistry with a commercial standard, which could also be enzymatically methylated to form vobasine (*N*4-methylperivine) with akuammidine 16*S* stereochemistry as we previously determined by NOESY NMR (Farzana *et al*., 2024). These results revealed the discrepancy between the hypothetical SBE product and the 16*S*-sarpagan alkaloid chemistry in *C. roseus*, suggesting the production of the akuammidine aldehyde that would give rise to the consistent stereochemistry in downstream products pericyclivine and perivine (Fig. 1).

**Table 1.** ^1^H NMR chemical shifts of alkaloids in this study.

|  | Akuammidine |  | Polyneuridine |  | Pericyclivine |  |  | Rhazimal | Strictamine |
| --- | --- | --- | --- | --- | --- | --- | --- | --- | --- |
|  | This study | Ref (Lounasmaa & Jokela, 1996) | This study | Ref (Lounasmaa & Jokela, 1996) | Ref (Qu *et al.*, 2018a) | This study | Ref (Lounasmaa & Jokela, 1996) | This study | This study |
|  | CDCl<sub>3</sub> | CDCl<sub>3</sub> | CDCl<sub>3</sub> | CDCl<sub>3</sub> | acetone-d<sub>6</sub> | CDCl<sub>3</sub> | CDCl<sub>3</sub> | CDCl<sub>3</sub> | CDCl<sub>3</sub> |
| **NH** | 7.75 br s | 7.90 br s | 7.77 br s | 7.81 br s | 9.87 br s | 7.73 br s | 7.78 br s | - | - |
| **3** | 4.23 d | 4.24 br d | 4.07 dd | 4.06 dd | 4.20 br d | 4.23 d | 4.21 br d | 4.62 d | 4.68 d |
| **5** | 3.06 d | 3.1 m | 4.27 d | 4.27 br d | 3.69 ddd | 3.68 ddd | 3.68 ddd | 2.62 ddd<br>2.67 ddd | 2.59 ddd<br>2.71 ddd |
| **6** | 2.91 dd<br>3.29 dd | 2.94 dd<br>3.30 dd | 2.94 dd<br>3.10 dd | 2.94 br d<br>3.10 dd | 2.80 m<br>3.20 dd | 2.91 dd<br>3.24 dd | 2.91 dd<br>3.24 dd | 2.12 dd<br>3.85 dd | 1.99 dd<br>3.74 ddd |
| **9** | 7.43 d | 7.42 d | 7.49 d | 7.48 d | 7.40 d | 7.41 d | 7.41 d | 7.56 d | 7.42 d |
| **10** | 7.05 dt | 7.05 dt | 7.10 dt | 7.10 dt | 6.98 dt | 7.04 dt | 7.04 dt | 7.22 dt | 7.16 dt |
| **11** | 7.11 dt | 7.11 dt | 7.15 dt | 7.15 dt | 7.05 dt | 7.11 dt | 7.10 dt | 7.37 dt | 7.33 dt |
| **12** | 7.28 d | 7.28 d | 7.31 d | 7.31 dd | 7.33 d | 7.29 d | 7.28 d | 7.63 d | 7.62 d |
| **14** | 1.86 dd<br>2.66 dd | 1.85 dd<br>2.67 dd | 1.88 m<br>1.90 ddd | 1.85 ddd<br>1.91 ddd | 1.74 dddd<br>2.63 ddd | 1.78 dddd<br>2.59 ddd | 1.76 ddd<br>2.58 ddd | 2.19 dd<br>2.49 ddd | 1.74 dd<br>2.68 ddd |
| **15** | 3.06 d | 3.1 m | 3.21 t | 3.21 dd | 2.96 br s | 2.99 br t | 2.98 m | 3.83 br s | 3.50 br s |
| **16** | - | - | - | - | 2.83 m | 2.82 dd | 2.82 dd | - | 2.08 d |
| **17** | 3.68 d<br>3.83 d | 3.67 d<br>3.83 d | 3.62 d<br>3.71 d | 3.61 d<br>3.71 d | - | - | - | 8.51 s | - |
| **18** | 1.65 dt | 1.65 ddd | 1.60 dt | 1.60 br d | 1.65 ddd | 1.62 dt | 1.62 ddd | 1.60 dd | 1.55 dd |
| **19** | 5.41 q | 5.39 br q | 5.28 qt | 5.28 br q | 5.28 q | 5.27 tq | 5.27 br q | 5.51 q | 5.50 q |
| **21** | 3.60 dd<br>3.61 dt | 3.58 def<br>3.58 def | 3.60 m<br>3.65 m | 3.6 m<br>3.6 m | 3.60 m<br>3.60 m | 3.64 t<br>3.64 t | 3.6 m<br>3.6 m | 3.15 d<br>4.12 d | 3.11 d<br>4.05 dt |
| **OCH<sub>3</sub>** | 2.94 s | 2.94 s | 3.73 s | 3.73 s | 3.10 s | 3.06 s | 3.75 s | 3.81 s | 3.72 s |

### NaBH_4_ reduces SBE products to 16*R* polyneuridine and 16*S* akuammidine epimers

*C. roseus* exclusively accumulates 16*S* sarpagan MIAs without any known 16*R* sarpagan MIAs or derivatives. Similarly, in *T. elegans*, the primary MIA is the 16*S*-sarpagan vobasine, with no evidence of 16*R* sarpagan MIAs (Mohammed *et al*., 2021; Farzana *et al*., 2024). In comparison, *Rauvolfia* and *Vinca* species produce both 16*S* derivatives, such as akuammidine and sarpagine, and 16*R* derivatives, such as quebrachidine, vincamajine (*N*1-methylquebrachidine), 11-hydroxypolyneuridinne, and ajmaline (Bartlett *et al*., 1962; Il’yashenko *et al*., 1977; Boğa *et al*., 2011; Bahadori *et al*., 2012; Adizov & Tashkhodjaev, 2019). This stereochemical variation further prompted us to investigate the stereochemistry of SBE products.

We expressed CrSBE, two new *T. elegans* and *V. minor* orthologs TeSBE and VmSBE, and a previously studied *R. serpentina* RsSBE2, sharing 72-82% amino acids identity among each other, in yeast to evaluate their in vitro activities. A phylogenetic analysis demonstrated close ancestry among SBEs, GOs, and RHSs and other MIA oxidizing CYPs (Fig. 3). As eukaryotic CYPs are membrane-bound, microsomes (total yeast membranes) were isolated for enzymatic assays. All four enzymes could convert 19*E*-geissoschizine with [M+H]^+^ *m/z* 353 into a broadly eluting compound of *m/z* 351, co-eluting with authentic polyneuridine aldehyde standard produced by the Evanno group (Fig. 4A). The broad peak was likely a result of conformer exchange and enol-keto resonance, similar to the substrate 19*E*-geissoschizing itself in our LC system. Tandem mass spectrometry (MS/MS) confirmed sarpagan bridge formation with characteristic daughter ion *m/z* 166, identical with the polyneuridine aldehyde standard (Supplementary fig. 2) (Dang *et al*., 2018; Guo *et al*., 2024). However, the broad-eluting behavior was unable to distinguish between the polyneuridine aldehyde and akuammidine aldehyde epimers. In addition to the cyclized aldehyde, a further oxidized but not cyclized byproduct with *m/z* 349 (Fig. 4B) was tentatively identified as 3,4,5,6-dehydrogeissoschizine, based on its nearly identical UV absorption profile with that of serpentine (3,4,5,6-dehydroajmalicine, Supplementary fig. 2). Such UV profile is indicative of fully aromatized, beta-carboline structure, which is different from the profile of indole at 280 nm absorption maxima.

**Figure 3.**
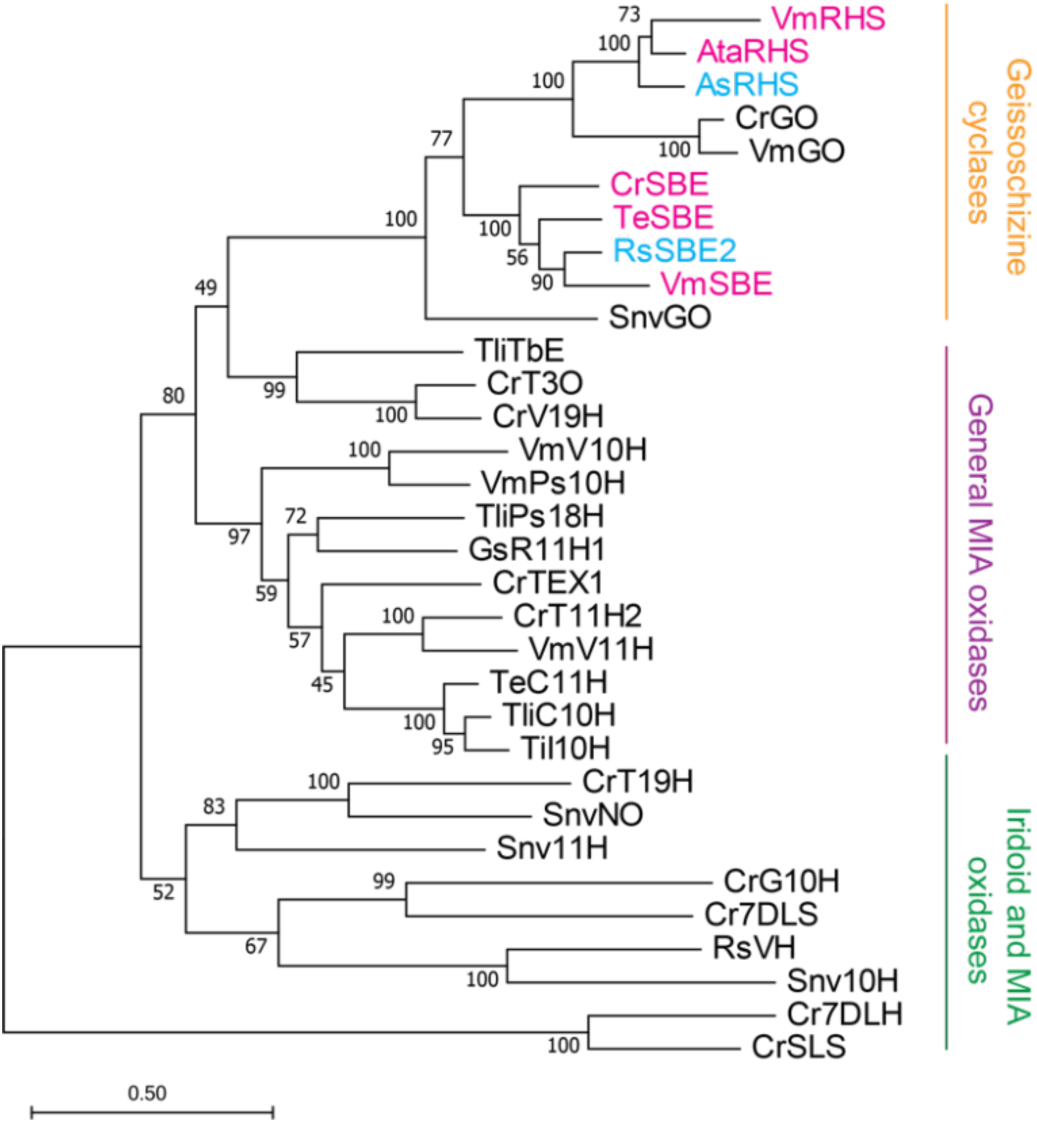
Phylogenetic analysis of geissoschizine oxidases (GOs), sarpagan bridge enzymes (SBEs), rhazimal synthases (RHSs), and other characterized cytochrome P450 monooxygenases (CYPs) involved in MIA biosynthesis. Newly identified and characterized enzymes from this study are highlighted in red, while previously characterized RHS and SBE enzymes are marked in blue. Evolutionary analyses were conducted in MEGA11. The evolutionary history was inferred by using the Maximum Likelihood method and JTT matrix-based model. The tree with the highest log likelihood is shown. The percentage of trees in which the associated taxa clustered together is shown next to the branches (100 bootstrap replicates). Initial tree(s) for the heuristic search were obtained automatically by applying Neighbor-Join and BioNJ algorithms to a matrix of pairwise distances estimated using the JTT model, and then selecting the topology with superior log likelihood value. The tree is drawn to scale, with branch lengths measured in the number of substitutions per site (scale bar). As: *Alstonia scholaris*; Ata: *Amsonia tabernaemontana*; Cr: *Catharanthus roseus*; Gs: *Gelsemium sempervirens*; Snv: *Strychnos nux-vomica*; Rs: *Rauvolfia serpentina*; Te: *Tabernaemontana elegans*; Tli: *Tabernaemontana litoralis*; Ti: *Tabernanthe iboga*; Vm: *Vinca minor*; R11H1: rankinidine 11-hydroxylase 1; Ps18H: pseudovincadifformine 18-hydroxylase; I10H: ibogamine 10-hydroxylase; T16H2, tabersonine 16-hydroxylase 2; TEX1: tabersonine 14,15-α-epoxidase; TbE: tabersonine 14,15-β-epoxidase; T3O: tabersonine 3-oxidase; V19H: (+)-vincadifformine 19-hydroxylase; Snv10H: strychnine 10-hydroxylase; Snv11H: β-colubrine 11-hydroxylase; NO: norfluorocurarine 18-hydroxylase; T19H: tabersonine 19-hydroxylase; VH: vinorine hydroxylase; 7DLH: 7-deoxyloganic acid hydroxylase; SLS: secologanine synthase.

**Figure 4.**
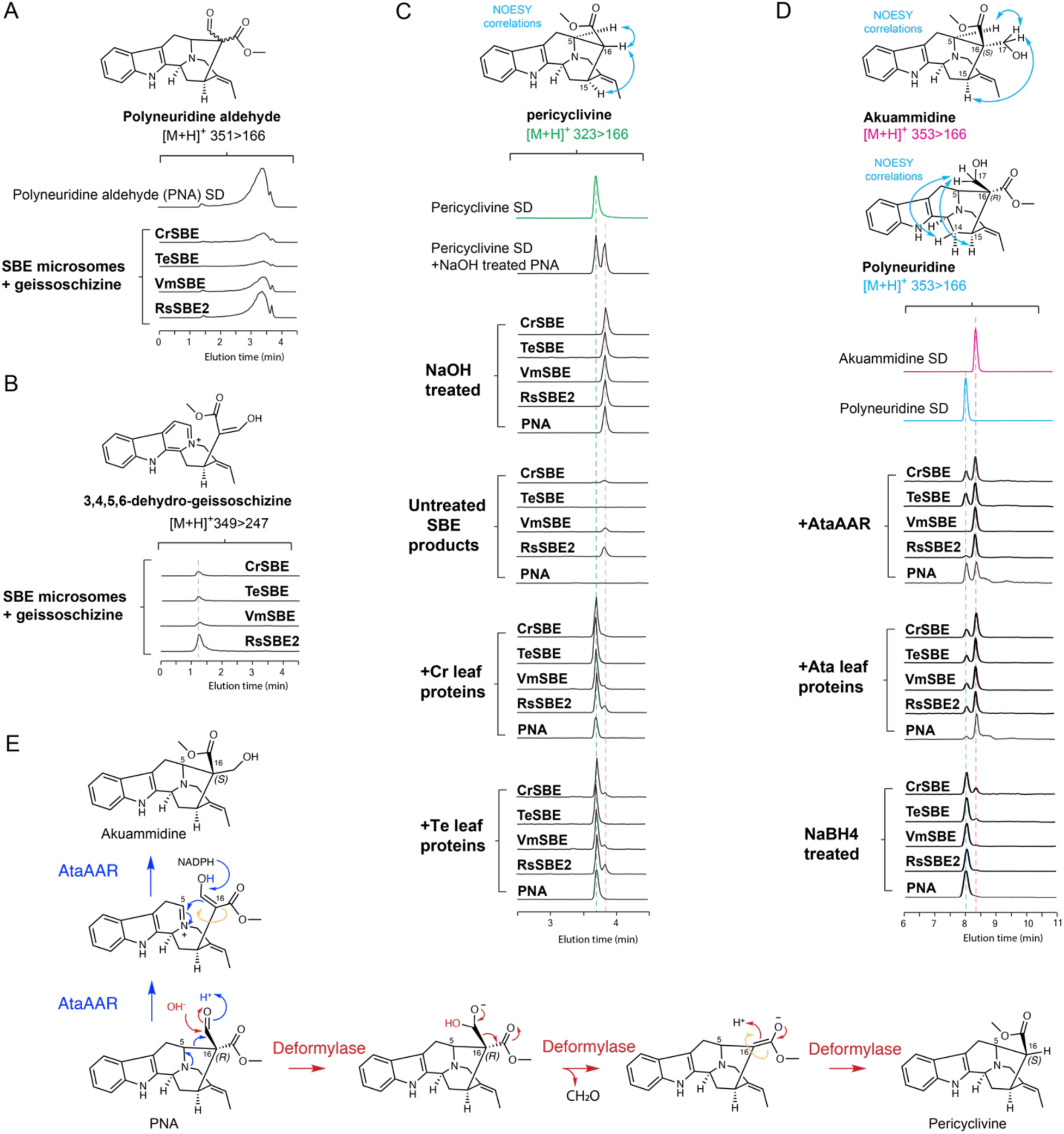
Sarpagan bridge enzymes (SBEs) from *C. roseus (Cr)*, *T. elegans (Te)*, *V. minor (Vm)*, and *R. serpentina (Rs)* catalyze geissoschizine sarpagan bridge formation, with downstream species-specific enzymes further converting the products into akuammidine and pericyclivine. (**A**) The products of SBEs from four species co-eluted with polyneuridine aldehyde (PNA) standard. (**B**) SBEs also produced 3,4,5,6-dehydrogeissoschizine byproduct. (**C**) Treating SBE products with NaOH or plant leaf proteins led to distinct deformylated products. *C. roseus* and *T. elegans* deformylated SBE products and PNA standard to a single product pericyclivine, while NaOH treatment formed 16-epi-pericyclivine instead. (**D**) Reducing SBE products with NaBH_4_ or plant leaf proteins led to opposite C16 stereochemistry. *A. tabernaemontana (Ata)* leaf proteins and recombinant akuammidine aldehyde reductase (AtaAAR) both reduced SBE products and PNA to akuammidine with minor amounts of polyneuridine. NaBH_4_ reduction instead produced mostly 16-epi-pericyclivine. (**E**) Proposed reduction and deformylation reaction mechanism, involving putative C16-epimerization steps. The LC-MS/MS chromatograms show multiple reaction monitoring (MRM) transitions using parameters [M+H]^+^ *m/z* 351>166 for the aldehydes, *m/z* 349>247 for 3,4,5,6-dehydrogeissoschizine, *m/z* 323>166 for pericyclivine, and *m/z* 353>166 for akuammidine and polyneuridine. The MS/MS spectra for these alkaloids are shown in Supplementary figure 2.

Solid polyneuridine aldehyde is stable yet it readily degrades in solution (Ahamada *et al*., 2016; Turpin *et al*., 2020). While it has been frequently described in literature, its NMR spectroscopic study was achieved only once by the Evanno group, by oxidizing quebrachidine to polyneuridine aldehyde and quickly analysing it in deuterated DMSO (Ahamada *et al*., 2016). Akuammidine aldehyde, however, has not been studied by NMR. Due to the instability issue, enzymatically produced sarpagan bridge products have never been properly examined. Instead, their identity has been indirectly inferred by their conversions into downstream MIAs, such as vinorine via polyneuridine aldehyde esterase (PNAE) and vinorine synthase (VS) (Pfitzner & Stöckigt, 1983; Dogru *et al*., 2000; Bayer *et al*., 2004; Gerasimenko *et al*., 2004; Dang *et al*., 2018; Guo *et al*., 2024), or vellosimine and tombozine (both with inverted 16*S* stereochemistry) by total *R. serpentina* total proteins (Fig. 1) (Pfitzner *et al*., 1984; Schmidt & Stöckigt, 1995).

Due to the instability of the aldehyde and the scale of in vitro assays, we studied the aldehyde stereochemistry by NaBH_4_ reduction and compared the reduced alcohols to authentic akuammidine and polyneuridine standards. The akuammidine standard was purified from *A. tabernaemontana* (Supplementary fig. 9) in this study and we confirmed the structure and C16 stereochemistry by NMR (Supplementary fig. 10-15). The H17, H5, and H15 correlations in NOESY spectra indicated its 16*S* stereochemistry. NMR experiments of polyneuridine acquired from the Evanno group, showed H17 correlation to H6, H14, and H15 in NOESY spectra, confirming the 16*R* stereochemistry (Supplementary fig. 16-21) (Ahamada *et al*., 2016; Turpin *et al*., 2020). Additionally, the two alcohols could be easily distinguished through ^1^H NMR in CDCl_3_ by comparing their chemical shifts of carbomethoxy group (δ 2.94 ppm for akuammidine caused by strong indole shielding and δ 3.73 ppm for polyneuridine) and H5 (δ 3.06 for akuammidine and 4.27 for polyneuridine) (Table 2).

**Table 2.** ^13^C NMR chemical shifts of alkaloids in this study.

|  | Akuammidine |  | Polyneuridine |  | Pericyclivine |  |  | Rhazimal |  | Strictamine |  |
| --- | --- | --- | --- | --- | --- | --- | --- | --- | --- | --- | --- |
|  | This study | Ref (Lounasmaa & Jokela, 1996) | This study | Ref (Lounasmaa & Jokela, 1996) | Ref | This study | Ref (Lounasmaa & Jokela, 1996) | This study | Ref (Atta-ur-Rahman & Habib-ur-Rehman, 1986) | This study | Ref (Atta-ur-Rahman & Habib-ur-Rehman, 1986) |
|  | CDCl<sub>3</sub> | CDCl<sub>3</sub> | CDCl<sub>3</sub> | CDCl<sub>3</sub> | acetone-d<sub>6</sub> | CDCl<sub>3</sub> | CDCl<sub>3</sub> | CDCl<sub>3</sub> | CDCl<sub>3</sub> | CDCl<sub>3</sub> | CDCl<sub>3</sub> |
| **2** | 137.4 | 136.6* | 137.09 | 136.2* | 138.8 | 137.5 | 136.6* | 189.4 | 190.50 weak | 191.4 | 190.98 |
| **3** | 50.7 | 51.4 | 49.14 | 49.0 | 50.0 | 50.6 | 50.4 | 54.8 | 54.70 | 55.2 | 55.15 |
| **5** | 58.1 | 58.0 | 53.72 | 53.6 | 52.7 | 53.2 | 53.0 | 51.9 | 51.80 | 52.1 | 51.98 |
| **6** | 25.0 | 24.7 | 22.51 | 22.3 | 23.9 | 24.4 | 24.2 | 37.7 | 37.70 | 33.9 | 36.13* |
| **7** | 106.5 | 106.2 | 106.39 | 106.2 | 104.4 | 105.9 | 105.7 | 57.7 | - | 56.3 | 58.40 weak |
| **8** | 127.1 | 126.9 | 126.70 | 126.5 | 127.3 | 127.2 | 127.1 | 143.6 | 138.60* | 146.3 | 138.15* |
| **9** | 118.2 | 118.0 | 118.51 | 118.3 | 117.2 | 118.0 | 117.8 | 124.7 | 124.56 | 123.6 | 123.47 |
| **10** | 119.6 | 119.4 | 119.65 | 119.5 | 118.4 | 119.4 | 119.2 | 126.4 | 126.23 | 125.7 | 125.61 |
| **11** | 121.6 | 121.5 | 121.78 | 121.6 | 120.5 | 121.5 | 121.3 | 129.0 | 128.81 | 128.3 | 128.19 |
| **12** | 111.0 | 110.9 | 110.97 | 110.8 | 110.9 | 111.0 | 110.8 | 121.6 | 120.2 | 121.1 | 119.93 |
| **13** | 136.7 | 137.0* | 136.33 | 136.5* | 136.9 | 136.7 | 137.4* | 155.8 | 155.80 weak | 155.7 | 156.10 weak |
| **14** | 29.4 | 29.2 | 29.03 | 28.9 | 27.0 | 27.1 | 26.9 | 32.2 | 32.1 | 36.2 | 33.62* |
| **15** | 29.6 | 29.4 | 30.74 | 30.6 | 27.7 | 27.4 | 27.2 | 34.2 | 34.2 | 32.6 | 32.50 |
| **16** | 55.6 | 50.3 | 53.58 | 53.4 | 43.9 | 44.0 | 43.8 | 70.2 | 67.80 weak | 55.5 | 55.64 |
| **17** | 69.0 | 68.8 | 63.37 | 63.2 | - | - | - | 194.8 | 194.6 | - | - |
| **18** | 13.1 | 13.0 | 12.90 | 12.7 | 12.3 | 13.0 | 12.9 | 13.3 | 13.3 | 13.1 | 12.97 |
| **19** | 116.7 | 116.8 | 116.12 | 116.0 | 113.2 | 114.6 | 114.4 | 120.4 | 121.5 | 119.9 | 121.02 |
| **20** | 137.7 | 137.1* | 136.74 | 136.9* | 140.9 | 139.6 | 139.5* | 138.5 | 143.30* weak | 138.4 | 146.35* |
| **21** | 55.6 | 55.5 | 55.93 | 55.8 | 55.8 | 56.2 | 56.0 | 53.8 | 53.6 | 53.8 | 53.8 |
| **CH<sub>3</sub>-O-CO** | 174.1 | 173.8 | 176.58 | 176.4 | 172.5 | 173.0 | 172.9 | 168.2 | 171.00 weak | 171.8 | 171.8 |
| **CH<sub>3</sub>-O-CO** | 51.5 | 50.6 | 52.35 | 52.2 | 49.8 | 50.8 | 50.9 | 52.8 | 52.6 | 51.7 | 51.7 |

\* The assignments may be reversed.

When compared to the alcohol standards, NaBH₄ reduction of SBE products yielded both alcohol epimers, with polyneuridine being the dominant form (Fig. 4C). Relative to VmSBE and RsSBE2, the reduced products from CrSBE and TeSBE reactions contained a higher proportion of akuammidine. In contrast, NaBH₄ reduction of authentic polyneuridine under identical conditions produced only a single product, polyneuridine (Fig. 4C). This apparent discrepancy among the four SBEs prompted us to further investigate the mixed stereochemistry by deformylation.

### Enzymatic deformylation of SBE products yielded a single product pericyclivine, whereas base-catalyzed deformylation produced a pericyclivine epimer

Meteignier et al. (2025) reported high pH-induced deformylation of the MIA rhazimal to produce strictamine (16-deformyl-rhazimal). Inspired by this discovery, we treated the SBE products with NaOH. Treating either the SBE products or the polyneuridine aldehyde standard at pH 12 led exclusively to a single deformylated product, eluting slightly behind the pericyclivine standard (Fig. 4D), but sharing an identical MS/MS fragmentation pattern with a dominant *m/z* 166 daughter ion, indicating a preserved sarpagan bridge structure. This product was also detected as a minor by-product in our SBE in vitro reactions at pH 7.5. We tentatively identified this MIA as 16-epi-pericyclivine. The deformylation experiments yet suggested that all four SBEs produce only the 16*S* polyneuridine aldehyde.

In contrast to base-catalyzed deformylation, treating the SBE products with total desalted leaf proteins from *C. roseus* and *T. elegans* resulted exclusively in pericyclivine as the single deformylated product (Fig. 4D). Unexpectedly, *C. roseus* and *T. elegans* leaf proteins also transformed pure polyneuridine aldehyde into pericyclivine, despite its opposite C16 stereochemistry (Fig. 4D). We propose that this unidentified deformylation reaction involves enolate formation facilitated by the neighboring carbomethoxy group. Reformation of the carbomethoxy ketone leads to 16-epimerization at enzyme active site, yielding pericyclivine with the akuammidine stereochemistry (Fig. 4E). Pericyclivine then is subsequently transformed into naturally occurring perivine in *C. roseus* (Fig. 2B, C) and the widely distributed MIA vobasine in *T. elegans* (Farzana *et al*., 2024). The apparent epimerization during enzymatic deformylation also provides a mechanistic explanation for the absence of 16*R*-sarpagan MIAs in these two species.

### Identification of an akuammidine aldehyde reductase that preferentially yields akuammidine

Since akuammidine has been purified from *A. tabernaemontana*, we hypothesized that reductases responsible for akuammidine aldehyde reduction may exist in this species and other MIA-producing plants. Desalted leaf proteins from *A. tabernaemontana* efficiently reduced not only the SBE products but also the polyneuridine aldehyde standard, yielding predominantly akuammidine along with minor amounts of polyneuridine (Fig. 4C). This intriguing epimerization from 16*R* to 16*S* suggests that the sarpagan bridge likely opens prior to aldehyde reduction. Following epimerization and reduction, the sarpagan bridge reforms at enzyme active site to complete the reaction (Fig. 4E).

To date, two aldehyde reductases have been characterized for MIA biosynthesis. CrRedox2 reduces the stemmadenine aldehyde to its alcohol form stemmadenine (Qu *et al*., 2018a). Its homolog in *A. scholaris*, namely the rhazimal reductase AsRHR1/2 sharing 82% and 77% amino acid identity respectively with CrRedox2, reduce rhazimal to rhazimol (Wang *et al*., 2022). We suspect that the putative akuammidine aldehyde reductase (AAR) activities were contributed by homologous aldehyde reductases. We searched the *A. tabernaemontana* leaf transcriptome and identified two AAR candidates. One of the two purified, his-tagged proteins (Supplementary fig. 22) exhibited reductase activity, successfully converting SBE products predominantly into akuammidine with small amounts of polyneuridine (Fig. 4C). Pure polyneuridine aldehyde was reduced to rather equal amounts of alcohol epimers (Fig. 4C) We therefore named this enzyme AtaAAR, which shared 85% and 91% amino acid identity with CrRedox2 and AsRHR1, respectively. The identification of AtaAAR thus explains the formation of akuammidine in *A. tabernaemontana*.

### Polyneuridine aldehyde esterase and akuammidine aldehyde reductase produce vellosimine and tombozine C16-epimers

Despite which aldehyde epimer may be produced by SBEs, the enzymatic deformylation reaction exclusively produces a single product pericyclivine sharing the akuammidine aldehyde stereochemistry. Interestingly, the Stöckigt group previously also reported that the *R. serpentina* polyneuridine aldehyde esterase (RsPNAE) catalyzes the decarbomethoxylation of both polyneuridine and akuammidine aldehyde (Fig. 1) (Corey & Kim, 1972; Pfitzner *et al*., 1984). While the fate of akuammidine aldehyde in this reaction was not further explored, the group indicated that short incubation of polyneuridine aldehyde with PNAE predominantly yielded 16-epi-vellosimine, which retains the same stereochemistry with the polyneuridine aldehyde precursor. Prolonged incubation with PNAE resulted in the gradual epimerization of 16-epi-vellosimine to vellosimine. These epimers were distinguished by comparing the NMR chemical shifts of their aldehyde groups.

To elucidate the products of SBE reaction and further investigate the fate of sarpagan C16 stereochemistry, we repeated these experiments using purified RsPNAE (Supplementary fig. 22) on both pure polyneuridine aldehyde and SBE products. Within just 1 minute of incubation with RsPNAE, polyneuridine aldehyde was converted to vellosimine epimers (*m/z* 293, Fig. 5A, B, Supplementary fig. 2), with the earlier-eluting epimer accumulating in greater quantity. Prolonged incubation (80 minutes) with PNAE resulted in the gradual epimerization of the earlier-eluting epimer into the later-eluting epimer, consistent with the Stöckigt group’s findings. We identified the earlier-eluting epimer as 16-epi-vellosimine and the later-eluting epimer as vellosimine based on Stöckigt’s assignment.

**Figure 5.**
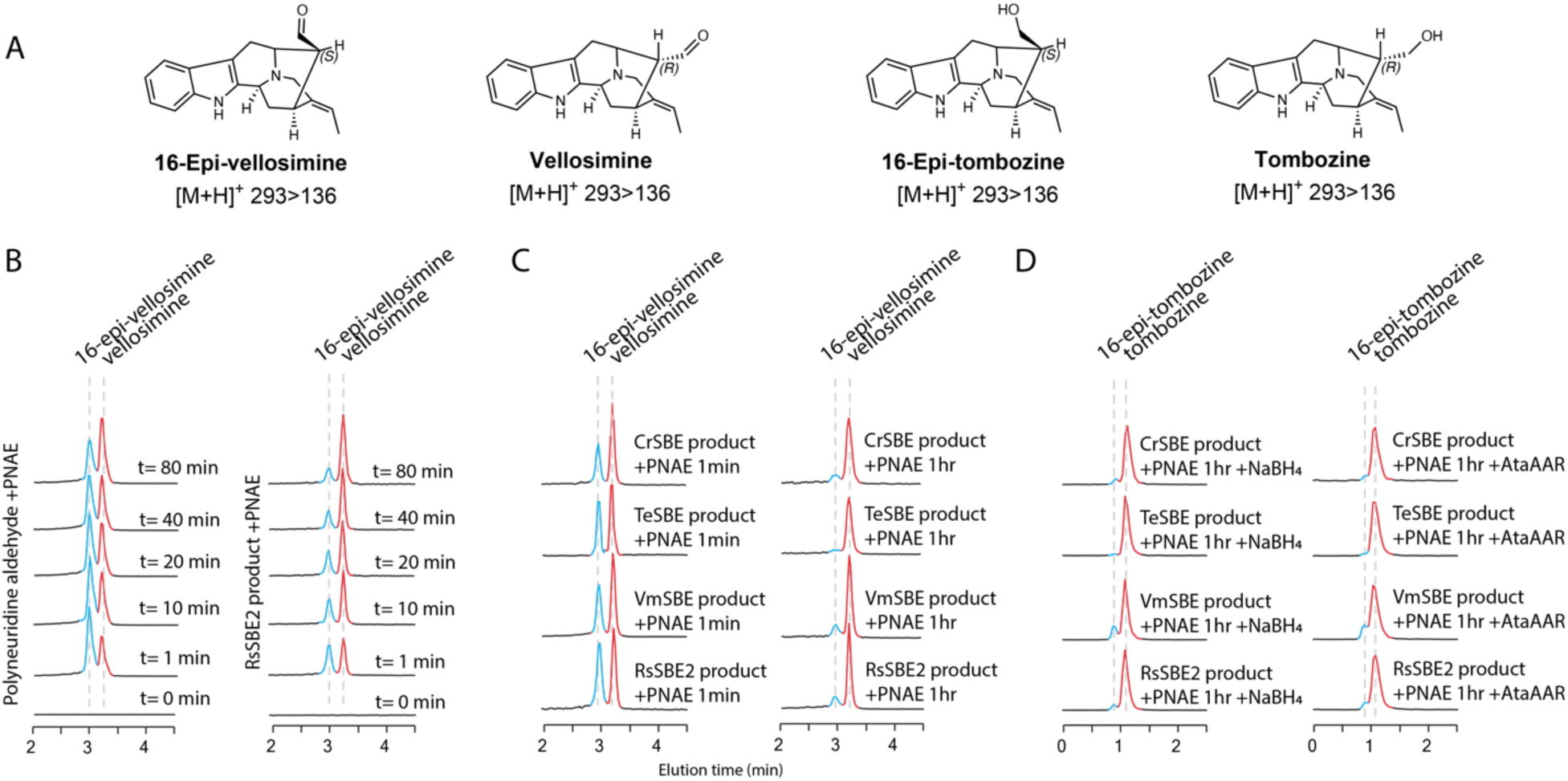
Polyneuridine aldehyde esterase (PNAE) catalyzes the decarbomethoxylation of SBE products and polyneuridine aldehyde, producing a mixture of vellosimine C16-epimers, which are subsequently reduced to their corresponding alcohols by akuammidine aldehyde reductase (AtaAAR) (**A**) Structures of MIAs in these reactions. (**B**) PNAE produced vellosimine epimers from polyneuridine aldehyde and RsSBE2 products. 16-epi-vellosimine gradually epimerized to vellosmine overtime in the presence of PNAE. (**C**) PNAE reaction with products from four SBEs at 1 min and 1 hr. The PNAE mediated epimerization of 16-epi-vellosimine to vellosimine is evident in these reactions. (**D**) AtaAAR reduces both vellosimine epimers to their corresponding alcohols, 16-epi-tombozine and tombozine. The LC-MS/MS chromatograms show multiple reaction monitoring (MRM) transitions using parameters [M+H]^+^ *m/z* 293>136 for vellosimine epiemrs and *m/z* 295>138 for tombozine epimers. The MS/MS spectra for these alkaloids are shown in Supplementary figure 2.

When the products of four SBE were subjected to PNAE assays, both vellosimine epimers were again rapidly formed in just a minute (Fig. 5B, C). Over the course of 80 minutes, PNAE-mediated epimerization led to increased accumulation of vellosimine sharing the akuammidine C16 stereochmistry (Fig. 5B, C). Both vellosimine epimers could be further reduced by the newly identified AtaAAR, yielding their corresponding alcohols (Fig. 5D). Identical results were obtained when reducing the epimers with NaBH_4_ (Fig. 5D).

### Rhazimal synthases from three species produce strictly 16*R* product

Oxidative C7–C16 cyclization of 19*E*-geissoschizine theoretically produces rhazinaline (16*S*), reported in *Rhazya stricta* (Chatterjee *et al*., 2006) and rhazimal (16*R*). Rhazimal synthase (RHS) was first identified and characterized in *Alstonia scholaris*. Sharing 62% amino acid identity with CrGO, AsRHS generates both rhazimal and preakuammicine aldehyde iminium, with the latter spontaneously deformylating to akuammicine (Fig. 1). These findings were recently corroborated by the characterization of several AsRHS variants in *A*. *scholaris* (Méteignier *et al*., 2025). The C16 stereochemistry of AsRHS products was confirmed by reducing the products to their corresponding alcohols and comparing them with a chemically synthesized rhazimol standard (Zhang *et al*., 2019; Wang *et al*., 2022).

To further investigate the natural occurrence of these compounds, we purified rhazimal directly from *A. tabernaemontana* leaves (Supplementary Fig. 9) and confirmed its structure and 16*R* stereochemistry using NMR (Tables 1 and 2, Supplementary Fig. 23–28). NOESY spectra revealed correlations between H17 and H15, H14, and indole H9, consistent with the 16*R* configuration. Instead of rhazimal, we found and purified strictamine from *V. minor* leaves, which derives biosynthetically from rhazimal deformylation, (Tables 1 and 2, Supplementary Fig. 29–34). Similar to rhazimal, the H16 correlations with H15, H14, and indole H9 confirmed the 16*R* stereochemistry of strictamine, confirming its intermediacy from rhazimal. These findings suggested that akuammiline type MIA biosynthesis occurs in both *A. tabernaemontana* and *V. minor*.

Using the *A. scholaris* RHS sequence as a reference, we searched transcriptomes from *A. tabernaemontana* and *V. minor* and identified two new RHS orthologs AtaRHS and VmRHS, which share 84% and 74% amino acid identity with AsRHS, respectively. Given the phylogenetic proximity of *A. tabernaemontana* and *R. stricta*, AtaRHS might produce both rhazimal epimers. To test this, we expressed AsRHS, AtaRHS, and VmRHS in baker’s yeast. Microsomes containing these enzymes oxidized 19*E*-geissoschizine to both rhazimal and akuammicine, as confirmed by LC-MS/MS comparison with authentic standards (Fig. 6). Notably, we observe no evidence of a rhazimal 16-epimer for any of the three enzymes. The akuammicine standard was isolated from a *C. roseus* mutant that accumulates significant quantities of this compound, with its structure previously verified by NMR (Qu *et al*., 2018a; Li *et al*., 2024).

**Figure 6.**
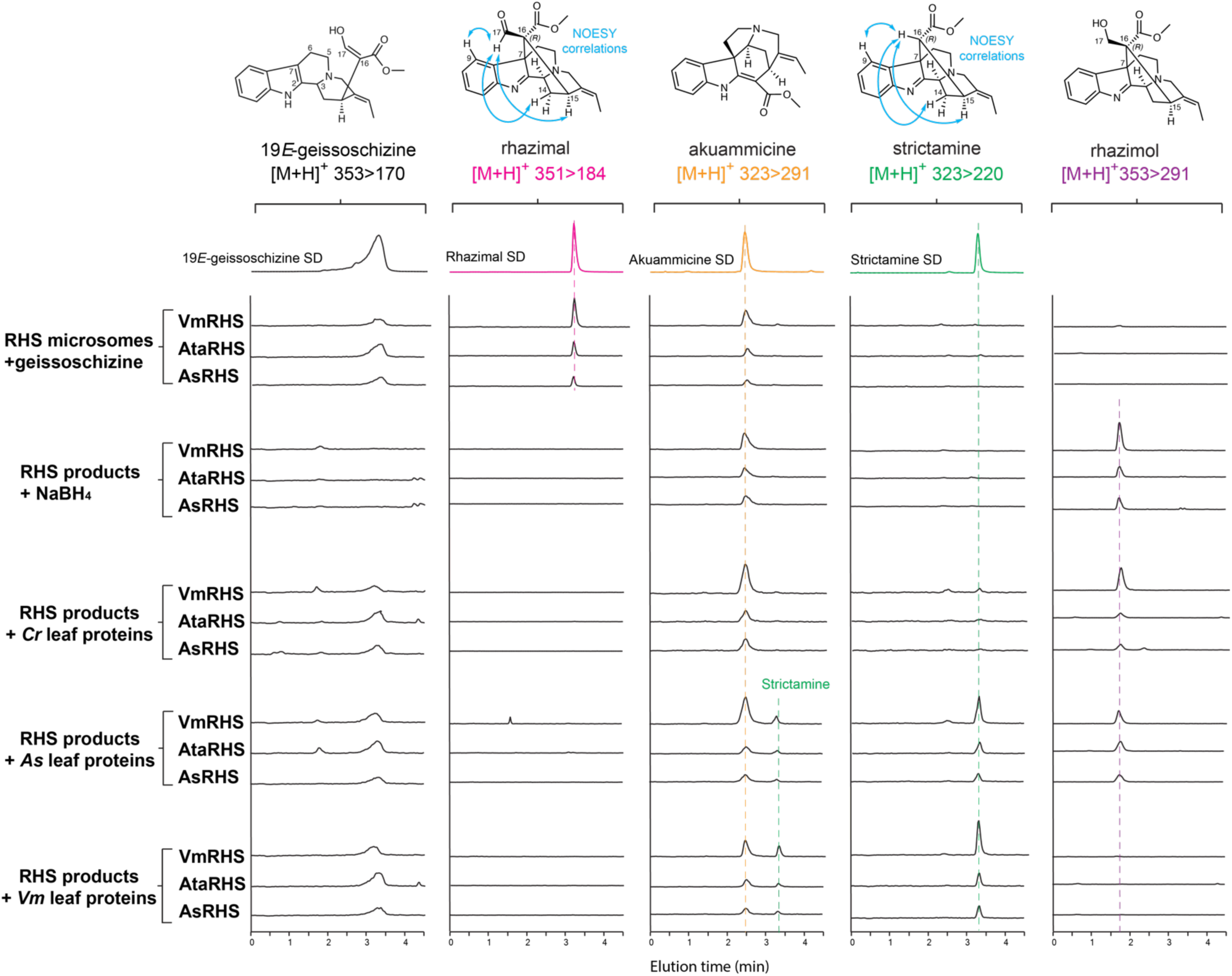
Rhazimal synthases (RHSs) from *A. scholaris (As)*, *A. tabernaemontana (Ata)*, and *V. minor (Vm)* catalyzed in vitro production of rhazimal (16*R*) but not rhazinaline (16S) from 19*E*-geissoschizine, which are subsequently converted enzymatically to rhazimol and strictamine by species-specific enzyme activities. RHSs also generated significant amounts of akuammicine. Total leaf proteins of *V. minor* deformylated rhazimal to strictamine preserving the C16 stereochemistry. In contrast, total leaf proteins of *C. roseus* reduced rhazimal to rhazimol while *A. scholaris* leaf proteins exhibited dual activity, producing both strictamine and rhazimol simultaneously. NaBH₄ reduction of RHS products yielded a single product rhazimol, further confirming rhazimal’s stereochemical identity. The LC-MS/MS chromatograms show multiple reaction monitoring (MRM) transitions using parameters [M+H]^+^ *m/z* 353>170 for geissoschizine, *m/z* 351>184 for rhazimal, *m/z* 323>291 for akuammicine, *m/z* 323>220 for strictamine, and *m/z* 353>291for rhazimol. Strictamine also exhibits m/z 323 > 291 ion transitions, which are observed alongside those of akuammicine in the same chromatograms. The MS/MS spectra for these alkaloids are shown in Supplementary figure 2.

Given that several Apocynaceae plants exhibit deformylation activity on akuammidine/polyneuridine aldehydes, it is plausible that some species may possess rhazimal deformylase to produce strictamine. Consistent with this hypothesis, we found that the total leaf proteins from *V. minor* and *A. scholaris* could deformylate rhazimal to strictamine (Fig. 6), providing a biochemical explanation for the biosynthesis of strictamine in *V. minor*. In contrast, *A. tabernaemontana* leaf proteins and *C. roseus* leaf proteins lacked deformylation activity, consistent with the observed accumulation of rhazimal but not strictamine in *A. tabernaemontana* (Supplementary fig. 9). Additionally, we discovered that leaf proteins from *A. scholaris* and *C. roseus* could reduce rhazimal to rhazimol, which co-eluted with the alcohol generated by NaBH₄ reduction (Fig. 6). These findings further confirmed the absence of rhazinaline biosynthesis activity of the three tested RHS enzymes, while validating the presence of reductase and deformylase activities in Apocynaceae plants, which together facilitate the biosynthesis of rhazimol and strictamine.

## Discussion

With the rapid advancement of plant transcriptome and genome sequencing at tissue and cellular levels, remarkable progress has been made in understanding monoterpene indole alkaloid (MIA) biosynthesis. The elucidation of the complete biosynthetic pathways for over 30 steps of anhydrovinblastine production *in C. roseus* and 20 steps of ajmaline biosynthesis in *R. serpentina* has provided a foundational framework for studying MIA biogenesis (Qu *et al*., 2015, 2018a,b; Dang *et al*., 2017, 2018; Caputi *et al*., 2018; Eng *et al*., 2022; Li *et al*., 2023; Sun *et al*., 2023; Guo *et al*., 2024). These breakthroughs have revealed the extraordinary enzymatic machinery responsible for the formation of the highly complex MIA skeletons.

MIA biosynthesis originates from the central precursor strictosidine. Reduction of strictosidine aglycone by cinnamyl alcohol dehydrogenase (CAD)-like reductases generates the corynanthe, yohimbine, and heteroyohimbine backbones, introducing multiple stereocenters in the process (Stavrinides *et al*., 2016; Qu *et al*., 2017; Kim *et al*., 2023; Stander *et al*., 2023). The next critical step involves three major cyclization events that catalyze the transformation of corynanthe-type MIA 19*E*-geissoschizine. The activities of homologous CYPs, geissoschizine oxidase (GO), sarpagan bridge enzyme (SBE), and rhazimal synthase (RHS), convert geissoschizine into strychnos, sarpagan, and akuammiline skeletons (Dang *et al*., 2018; Qu *et al*., 2018a; Wang *et al*., 2022). Subsequent modifications and structural decorations generate additional MIA types, such as iboga, aspidosperma, pseudoaspidosperma, and ajmalan, resulting in over 3,000 structurally diverse MIAs.

In this study, we investigated the stereochemical and mechanistic outcomes of geissoschizine cyclization mediated by SBE and RHS, using two previously characterized CYPs and five novel CYPs derived from six Apocynaceae species. Notably, the stereochemistry of the SBE products had remained elusive due to the instability of the intermediates The products of SBEs from four different species, as well as pure polyneuridine aldehyde, can be readily deformylated by NaOH, yielding a pericyclivine isomer with a distinct elution time (Fig. 4D). We tentatively identify this MIA as 16-epi-pericyclivine based on its MS/MS fragmentation and the expected base-catalyzed reaction mechanism (Fig. 4E, Supplementary fig. 2). While the formation of 16-epi-pericyclivine strongly suggests that polyneuridine aldehyde is the primary SBE product, the base-catalyzed deformylation step likely proceeds through an enolate intermediate, allowing epimerization to a more stable form (Fig. 4E). This conclusion is further complicated by the production of akuammidine from *C. roseus* and *T. elegans* SBEs following NaBH₄ reduction, albeit at lower levels than polyneuridine (Fig. 4C). It is possible that borate generated during reduction may catalyze 16-aldehyde epimerization via an enolate intermediate. Considering both sets of results, we suspect that SBE products are predominantly, if not exclusively during the initial sarpagan bridge formation, polyneuridine aldehyde with 16*R* stereochemistry. *C. roseus* and *T. elegans* SBEs may produce some akuammidine aldehyde ab initio. However, further studies using alternative methods or direct NMR characterization of larger-scale SBE products are needed to unequivocally determine the stereochemistry.

Regardless of whether SBE produces only polyneuridine aldehyde or a diastereomeric mixture, the apparent discrepancy between the polyneuridine aldehyde with downstream products harboring the opposite C16 stereochemistry, such as akuammidine, pericyclivine, perivine, and vobasine across Apocynaceae species, can be explained by the intriguing epimerization during enzymatic 17-aldehyde reduction, 16-deformylation, and 16-decarbomethoxylation steps. Under our testing conditions, AtaAAR reduced SBE products predominantly into akuammidine, with minor amounts of polyneuridine (Fig. 4C). This result is consistent with reductions using *A. tabernaemontana* total leaf proteins, which also produced akuammidine as the major product (Fig. 4C). We suspect that enzymatic 17-aldehyde reduction involves sarpagan bridge opening and 16-epimerization steps prior to, or concomitant with, the reduction step (Fig. 4E). This also explains the production of both polyneuridine and akuammidine epimers from pure polyneuridine aldehyde by AtaAAR and the preferential akuammidine production by *A. tabernaemontana* total leaf proteins. The apparent difference in the akuammidine/polyneuridine ratio across different reduction conditions suggests that additional factors and reaction parameters may influence the extent of epimerization, and the exact reaction mechanism warrants further investigation. Nonetheless, our findings shed light on the biosynthesis of akuammidine in *A. tabernaemontana* and other MIA-producing species.

The new AtaAAR shares 85% amino acid identity with *C. roseus* Redox2, a key enzyme responsible for reducing stemmadenine aldehyde to stemmadenine in the formation of iboga/aspidosperma MIAs (Qu et al., 2018a). It also shares 91% amino acid identity with *A. scholaris* rhazimal reductase 1, which reduces the aldehyde group on rhazimol (Wang et al., 2022). The similar aldehyde reduction activity observed among these reductases is consistent with their high sequence similarity. Additionally, AtaAAR’s aldehyde reduction activity extends to the vellosimine epimers, analogous to the polyneuridine/akuammidine aldehydes. It will be interesting to further investigate whether the other two aldehyde reductases can substitute for AtaAAR function, and vice versa, to explore their substrate specificity and catalytic mechanisms in more detail.

The enzymatic 16-deformylation of SBE products and PNA standard produced exclusively pericyclivine, in contrast to NaOH-catalyzed aldehyde deformylation, which yields 16-epi-pericyclivine (Fig. 4D). A sarpagan bridge opening mechanism, similar to that observed during AAR-mediated reduction, may underlie the enzymatic epimerization. Alternatively, epimerization may occur during the deformylation step via an enolate intermediate under strong enzyme stereochemical control (Fig. 4E). Identifying and characterizing the missing deformylase will be essential to elucidate the precise mechanistic details. Nonetheless, the production of pericyclivine from SBE products provides a clear explanation for the previously enigmatic origins of perivine and vobasine, which are derived from pericyclivine.

We also revisited the decarbomethoxylation of SBE products by RsPNAE. Our findings confirmed the biosynthesis of 16-epi-vellosimine and vellosimine from polyneuridine aldehyde and SBE products as a diastereomeric mixture by PNAE activity (Fig. 5). Prolonged incubation with PNAE favoured gradual epimerization of 16-epi-vellosimine into vellosimine (Fig. 5), likely via an enolate intermediate at enzyme active site. The acetylation of 16-epi-vellosimine by vinorine synthase channels it into the ajmaline biosynthetic pathway, while both epimers can be reduced by AtaAAR to form 16-epi-tombozine and tombozine. Vellosimine-reducing activity has also been previously documented in *R. serpentina* cell cultures (Pfitzner et al., 1984). Out of the two alcohol epimers, tombozine, bearing the akuammidine stereochemistry, can undergo additional hydroxylation at C10 to form sarpagine (Fig. 1), which is widely distributed in Apocynaceae (Pfitzner et al., 1984). The epimerization of 16-epi-vellosimine to vellosimine evidently contributes to tombozine becoming the predominant alcohol product.

The physiological relevance of CrSBE *in planta* was validated using virus-induced gene silencing (VIGS) in *C. roseus* leaves. While silencing CrSBE alone led to significant reduction of pericyclivine, this MIA naturally accumulate as a minor MIA in *C. roseus* (Figs. 2B, C, Supplementary Fig. 1D). To redirect metabolic flux toward sarpagan MIAs, we silenced CrGO, the major competitor of CrSBE for 19*E*-geissoschizine substrate. In this background, silencing CrSBE clearly reduced the accumulation of pericyclivine and perivine, confirming CrSBE’s in planta role in sarpagan biosynthesis and supporting the notion that GO and SBE compete for the same substrate (Fig. 2A–C).

In the case of RHS-mediated C7-C16 cyclization, we demonstrated that RHS enzymes *from A. scholaris, A. tabernaemontana*, and *V. minor* exclusively produce rhazimal (16R), not its epimer rhazinaline (16S) (Fig. 6). We verified the stereochemistry by direct rhazimal NMR analysis. This strict stereochemical control is consistent with CrGO (Qu *et al*., 2018a; Caputi *et al*., 2018). In the phylogenetic tree (Fig. 3), RHS, GO, and SBE enzymes clearly exhibit a monophyletic origin, indicating that all geissoschizine-cyclizing CYPs evolved from a common ancestor. RHSs and GOs are more closely related than SBEs, consistent with their shared stereospecificity in product formation and their role in producing the preakuammicine aldehyde iminium intermediate, which decomposes into akuammicine. This suggests that in RHS-bearing species, these enzymes likely also contributed to downstream iboga and aspidosperma MIA production, highlighting the evolutionary complexity of MIA biosynthesis. Although *A. tabernaemontana* is closely related to *R. stricta*, which reportedly produces rhazinaline (Chatterjee *et al*., 2006), its RHS did not show any evidence of rhazinaline production. It remains possible that *R. stricta* contains an epimerase or that its RHS produces both C16 diastereomers.

Additionally, we identified deformylase activity in the leaf proteins of *V. minor* and *A. scholaris*, which deformylated rhazimal to strictamine (Fig. 6). Although the rhazimal deformylase and the polyneuridine aldehyde deformylase share functional similarities, our results indicated that they are distinct enzymes. Specifically, in our experiments, *C. roseus* leaf proteins demonstrated the ability to deformylate polyneuridine aldehyde but showed no activity toward rhazimal (Fig. 4D, Fig. 6). The absence of rhazimal deformylase in *A. tabernaemontana* additionally explains the accumulation of rhazimal without strictamine. In addition to enzymatic rhazimal deformylation, Méteignier et al. (2025) demonstrated high-pH-induced rhazimal deformylation, providing a non-enzymatic method for accessing strictamine.

In conclusion, this study provides further stereochemical insights into SBE- and RHS-mediated cyclization reactions, deformylase, esterase, and reductase activities, and epimerization transformations that underpin the extraordinary structural diversity of MIAs. These findings deepen our understanding of how unique enzyme activities in Apocynaceae species shape the biosynthesis and accumulation of MIAs, contributing to their ecological and pharmacological significance.

## Material and Methods

### Chemicals

Catharanthine, vindoline, and vindolinine standards were purchased from Sigma Aldrich (Millipore Sigma, Burlington, MA, USA). Perivine was purchased from AvaChem Scientific (San Antonio, TX, USA). Vindorosine and epi-vindolinine standards were previously purified from *C. roseus* plant (Eng *et al*., 2022). 19*E-* and 19*Z-*geissoschizine were produced as previously described by reacting strictosidine aglycone produced by CrSTR and CrSGD with CrGS in vitro (Qu *et al*., 2017). The ethyl acetate extracted products were dried under vacuum and purified by preparative thin layer chromatography (TLC, Silica gel 60 F254, Millipore Sigma, Burlington, MA, USA) and solvents toluene: ethyl acetate: methanol 15:4:1 (v:v:v). Both pericyclivine and akuammicine were purified from VIGS plants in previous studies and structurally confirmed by NMR (Qu *et al*., 2018a). In this study, pericyclivine structure was confirmed again in CDCl_3_ with NOESY. For alkaloids purified from plant materials, 100-150 g of *A. tabernaemontana* or *V. minor* leaves were submerged in ethyl acetate for 30 min. The resulting extract was reduced in volume in a rotavap to ∼20 ml, and extracted by 1M HCl. The HCl extract was basified to pH 8 by NaOH and extracted by ethyl acetate to afford total alkaloids. *V. minor* alkaloids were separated by TLC with solvents ethyl acetate: methanol 9:1 (v:v). The band with Rf=0.09 was harvested and further separated on TLC with pure methanol to afford strictamine with Rf=0.47. *A. tabernaemontana* alkaloids were separated by TLC with solvents ethyl acetate: methanol 9:1 (v:v) to afford akuammidine (Rf=0.47) and rhazimal (Rf=0.17).

### Cloning

Full-length *CrSBE, VmSBE, VmRHS, AsRHS, AtaRHS, AtaAAR, RsPNAE,* and the VIGS fragment of *CrSBE* and *CrGO* were amplified from respective plant leaf cDNA using primers 1-18 (Supplementary table 2). Codon-optimized TeSBE was synthesized (Bio Basic, Markham, ON, Canada). VmRHS was cloned in pYES-DEST52 vector using Gateway® BP/LR clonases (ThermoFisher, Waltham, MA, USA) according to manufacture’s protocol. All other CYPs were cloned in pESC-Leu vector containing CrCPR, within BamHI/SalI sites. RsPNAE and AtaAAR was cloned in pET30b+ vector within BamHI/SalI sites. For silencing CrSBE and CrGO with a single chimeric construct, the VIGS fragment of CrSBE was cloned in EcoRI site of pTRV2 vector and the VIGS fragment of CrGO was cloned within XhoI/XbaI sites. For single GO-silencing, a previous pTRV2-GO construct was used. ^6^ All CYPs were mobilized to *Saccharomyces cerevisiae* strain BY4741 (MATα his3Δ1 leu2Δ0 met15Δ0 ura3Δ0 YPL154c::kanMX4). AtaAAR and RsPNAE were mobilized to BL21DE3 *E. coli* for protein expression. The pTRV2 vectors were mobilized to *Agrobacterium tumefaciens* strain GV3101. The sequences for new enzymes in this study have been deposited to NCBI Genbank (PQ564467-564471). The *Amsonia tabernaemontana* leaf RNA-seq data in this study has been deposited in NCBI under the bioproject PRJNA1256117. All remaining RNA-seq datasets from *C. roseus*, *V. minor*, *T. elegans*, and *A. scholaris* were obtained from publicly available datasets from NCBI.

### Virus-induced gene silencing

The VIGS experiments were carried out as described previously with a phytoene desaturase (PDS) silencing indicator (Qu *et al*., 2017). In brief, cells from overnight cultures of *A. tumefaciens* carrying pTRV1 vector and various pTRV2 vectors grown at 28 °C shaking incubator were collected by centrifugation and resuspended in infection buffer (10 mM MES pH 5.6, 10 mM MgCl_2_, 0.2 mM acetosyringone) to OD_600_ 1.5. The suspensions were incubated at 28 °C for 2.5 hr, then equal pTRV1 and pTRV2 suspensions were mixed just before the infection. A sharpened toothpick dipped in the mixed suspension was used to pierce just underneath the apical meristem of 4-week-old *C. roseus* (cv. Little Delicata for SBE single-silencing, and cv. Pacifica XP White for SBE/GO double silencing) with 2-3 pairs of true leaves, then 0.1 mL suspension mixture was used to flood the wound. The infected plants were grown at a glasshouse with 18/6 hr photoperiod at 25 °C for approximately 4 weeks, when the VIGS-PDS leaves developed white sectors. The developing leaves were cut in half along the main vertical vein. One half of the leaves were used for RNA extraction and qPCR, whereas the other half leaves were ground and extracted with methanol for alkaloid analysis using LC-MS/MS.

### cDNA synthesis and qRT-PCR

Plant leaves (10-100 mg) were ground in liquid nitrogen using a small pestle and a microtube. The total RNA were extracted with TRI Reagent^TM^ (ThermoFisher, Waltham, MA, USA) according to the manufacture’s protocol. The cDNA was synthesized using the LunaScript® RT SuperMix (New England Biolabs, Ipswich, MA, USA) according to the manufacture’s protocol. qRT-PCR experiments were performed on an Agilent AriaMx Real-Time PCR instrument using the SensiFAST SYBR No-ROX qPCR 2X master mix (FroggaBio, Concord, Canada) according to the manufacture’s protocol. The qRT-PCR (10 μL, 5 ng total RNA) cycles included 40 cycles of 95 °C for 10 s and 58 °C for 30 s. The Ct values and standard ΔΔCt method was used to quantify gene expression levels, which are normalized by using the expression of *C. roseus* 60S ribosomal RNA housekeeping gene (Qu *et al*., 2017). The changes of gene expression levels were evaluated by two-tailed, unpaired Student-t test from 7 independent biological samples with 3 technical replicates using Microsoft Excel.

### Yeast microsome isolation

Overnight culture (1 mL) of yeast was used to inoculate 100 mL SC-Leu medium with 2% glucose. The yeast was grown at 30°C for 24 h. The harvested cells were lysed in TES buffer (10 mM Tris-HCl pH 7.5, 1 mM EDTA, 0.6 M sorbitol) with glassbeads at 30 Hz, 3 min duration, and four times at 4°C using a Beadbug® homogenizer (Benchmark Scientific, Sayreville, NJ, USA). The lysate was centrifuged at 8,000 g for 10 min at 4°C, and the supernatant was further centrifuged at 100,000 g for 1 hr at 4 °C to pellet microsomes. The microsomes were suspended in TEG buffer (10 mM Tris-HCl pH 7.5, 1 mM EDTA, and 10% (v/v) glycerol) and stored in - 80°C.

### *E. coli* protein purification

LB media (200 mL) were first inoculated with 2 ml overnight *E. coli* BL21-DE3 cultures harboring various genes, then grown to OD_600_ 0.6-0.7 at 37 °C. The cultures were induced with 0.1 mM IPTG at 15 °C overnight. The harvested cells were sonicated in lysis buffer (20 mM Tris-HCl pH 7.5, 100 mM NaCl, 10 mM imidazole, 10% (v/v) glycerol). After centrifugation at 10,000 g for 10 min, the supernatants were incubated with 1 ml Ni-NTA resin at 4°C for 20 min. The resins were washed with 20 ml wash buffer (20 mM Tris-HCl pH 7.5, 200 mM NaCl, 30 mM imidazole, 10% (v/v) glycerol). The recombinant proteins were eluted with elution buffer (20 mM Tris-HCl pH 7.5, 100 mM NaCl, 250 mM imidazole, 10% (v/v) glycerol), desalted with a PD-10 desalting column (Citiva, Marlborough, MA, USA) into 20 mM Tris-HCl pH 7.5, 100 mM NaCl, 10% (v/v) glycerol, and stored at −80°C.

### Plant total leaf proteins preparation

Frozen leaves (2-3 g) from each plant species were ground into a fine powder in liquid nitrogen using a mortar and pestle. After mixing with 5% (w/w) polyvinylpolypyrrolidone (PVPP), the powder was suspended in ice cold sample buffer (20 mM Tris-HCl pH 7.5, 100 mM NaCl, 1 mM dithiothreitol, 10% (v/v) glycerol) and centrifuged at 20,000 g for 30 min at 4 °C. The aqueous phase containing total soluble proteins were desalted twice using a PD-10 desalting column equilibrated with the same buffer to obtain MIA-free total leaf proteins, which were stored −80°C until use.

### In vitro assay

Standard yeast microsome assays (100 μL) included 20 mM Tris-HCl pH 7.5, 100 ng geissoschizine, 1 mM NADPH, 50 μg yeast microsomal proteins, 1 μg *E. coli* recombinant proteins, and/or 20 μg plant total proteins. The reaction took place at 30 °C for 1 hr unless otherwise noted. The reaction was mixed with equal volume of methanol, centrifuged, and filtered for LC-MS/MS analysis.

### Chemical aldehyde reduction and deformylation

SBE products in 100 µl of 20 mM Tris-HCl buffer (pH 7.5) from the in vitro assays or MIAs (25 ng) dissolved in the same buffer were used as substrates. For reduction, NaBH₄ was first dissolved in water and immediately added to each sample at a final concentration of 1 µM. For the NaOH treatment, 2.5 M NaOH (2 µl) was added to the samples for a final pH 12. Samples from both treatments were incubated at room temperature for five minutes and then 100 µl of methanol was added. For NaOH treatment, the samples were additionally adjusted to neutral pH with Tris-HCl buffer before LC-MS/MS analysis.

### NMR and LC-MS/MS

NMR spectra were recorded on an Agilent 400 MR, a Bruker Avance III HD 400 MHz, or a Bruker Avance AV I 600 digital NMR spectrometer in acetone-*d6* or CDCl_3_. Alkaloids and in vitro assay samples were analyzed by Ultivo Triple Quadrupole LC-MS/MS system from Agilent (Santa Clara, CA, USA), equipped with an Avantor® ACE® UltraCore C18 2.5 Super C18 column (50×3 mm, particle size 2.5 *μ*m) as well as a photodiode array detector and a mass spectrometer. The following solvent systems were used: Solvent A, Methanol: Acetonitrile: Ammonium acetate 1 M: water at 29:71:2:398; solvent B, Methanol: Acetonitrile: Ammonium acetate 1 M: water at 130:320:0.25:49.7. The following linear elution gradient was used: 0-5 min 80% A, 20% B; 5-5.8 min 1% A, 99% B; 5.8-8 min 80% A, 20% B; the flow during the analysis was constant and 0.6 ml/min. The photodiode array detector range was 200 to 500 nm. The mass spectrometer was operated with the gas temperature at 300°C and gas flow of 10 L/min. Capillary voltage was 4 kV from m/z 100 to m/z 1000 with scan time 500 ms and the fragmentor performed at 135 V with positive polarity. The MRM mode was operated as same as scan mode with the adjusted precursor and product ion with collision energy (CE) 30 V.

To fully separate polyneuridine and akuammidine epimers, an Agilent Poroshell 120, EC-C18 column (4.6×100 mm, particle size 2.7 *μ*m) was used in combination with the solvent system: Solvent A: water; Solvent B: methanol. The following linear elution gradient was used: 0-0.5 min 100% A; 0.5 min 80% A, 20% B; 5 min 50% A, 50% B; 5-6 min 50% A, 50% B; 8 min 100% B; 8.3 min 100% A; 8.3-12 min: 100% A; the flow during the analysis was constant and 1 ml/min. The mass spectrometer was operated as described above.

## Acknowledgements

This research is supported by a Natural Sciences and Engineering Research Council of Canada (NSERC) Discovery Grant and a New Brunswick Innovation Foundation (NBIF) Research Assistantship Initiative Grant to Y.Q.

