## Supplementary information for "Stereochemical Insights into Sarpagan and Akuammiline Alkaloid Biosynthesis"

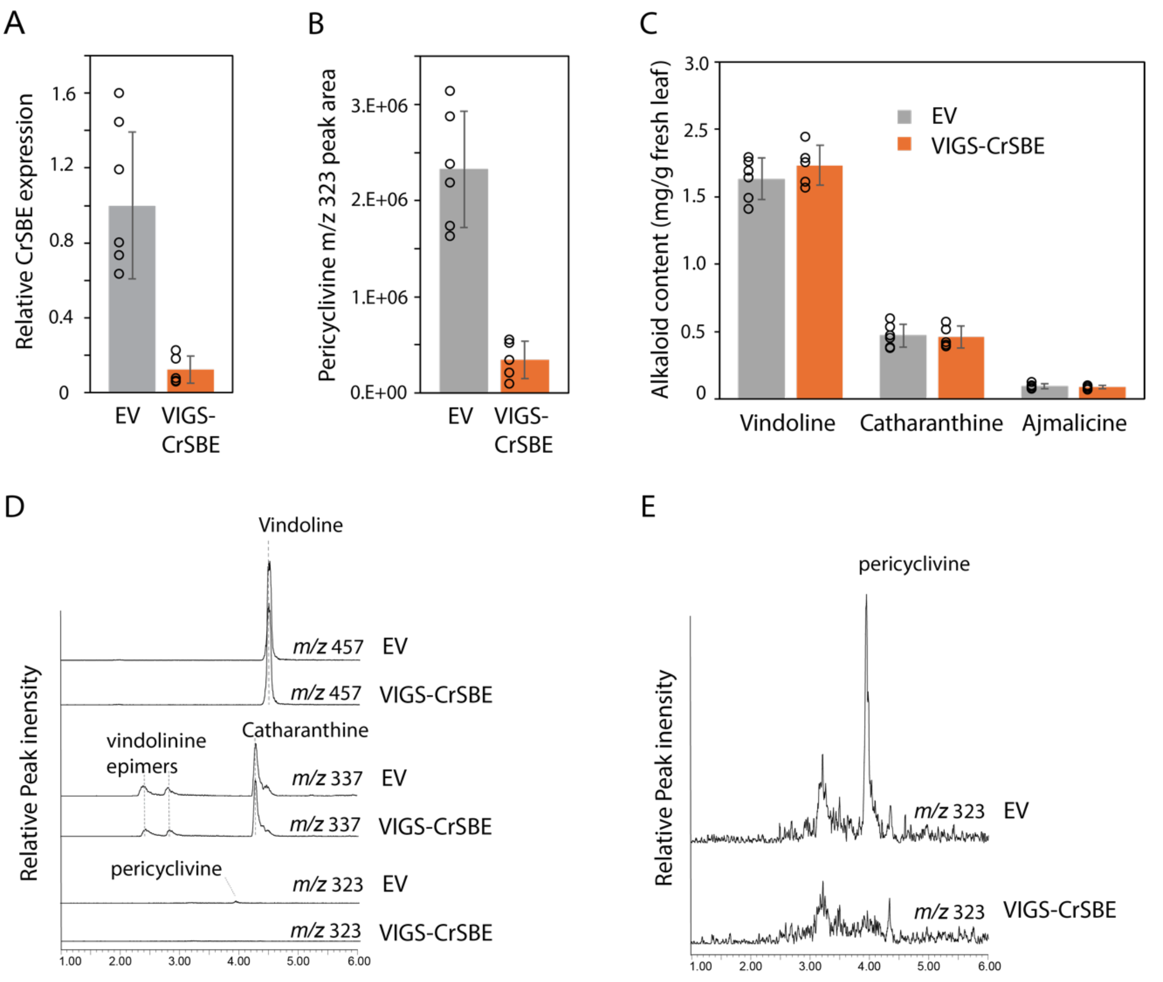

**Supplementary figure 1. Silencing CrSBE in *C. roseus* leaf by Virus induced gene silencing (VIGS) led to significant pericyclivine reduction while not affecting major leaf alkaloids.** (**A**) qRT-PCR analysis showing significant reduction of CrSBE transcript levels in VIGS plants compared to the Empty Vector (EV) controls. (**B**) Pericyclivine levels dropped to 12.3 % in VIGS plants. (**C**) Major alkaloids vindoline, catharanthine, and ajmalicine were not impacted by CrSBE-silencing. (**D**) Representative LC-MS chromatograms from EV and VIGS plants showing [M+H]+ *m/z* 457 for vindoline, *m/z* 337 for catharanthine and vindolinine 19-epimers, and *m/z* 323 for pericyclivine, which accumulated at very low levels in EV plants. (**E**) Scaled-up LC chromatograms from (D) showing the difference of pericyclivine levels in EV and VIGS plant. Data was generated from six EV plants and five VIGS-CrSBE plants. Individual data points are shown as circles, and error bars represent standard deviation. The structures of alkaloids are found in Figure 1.

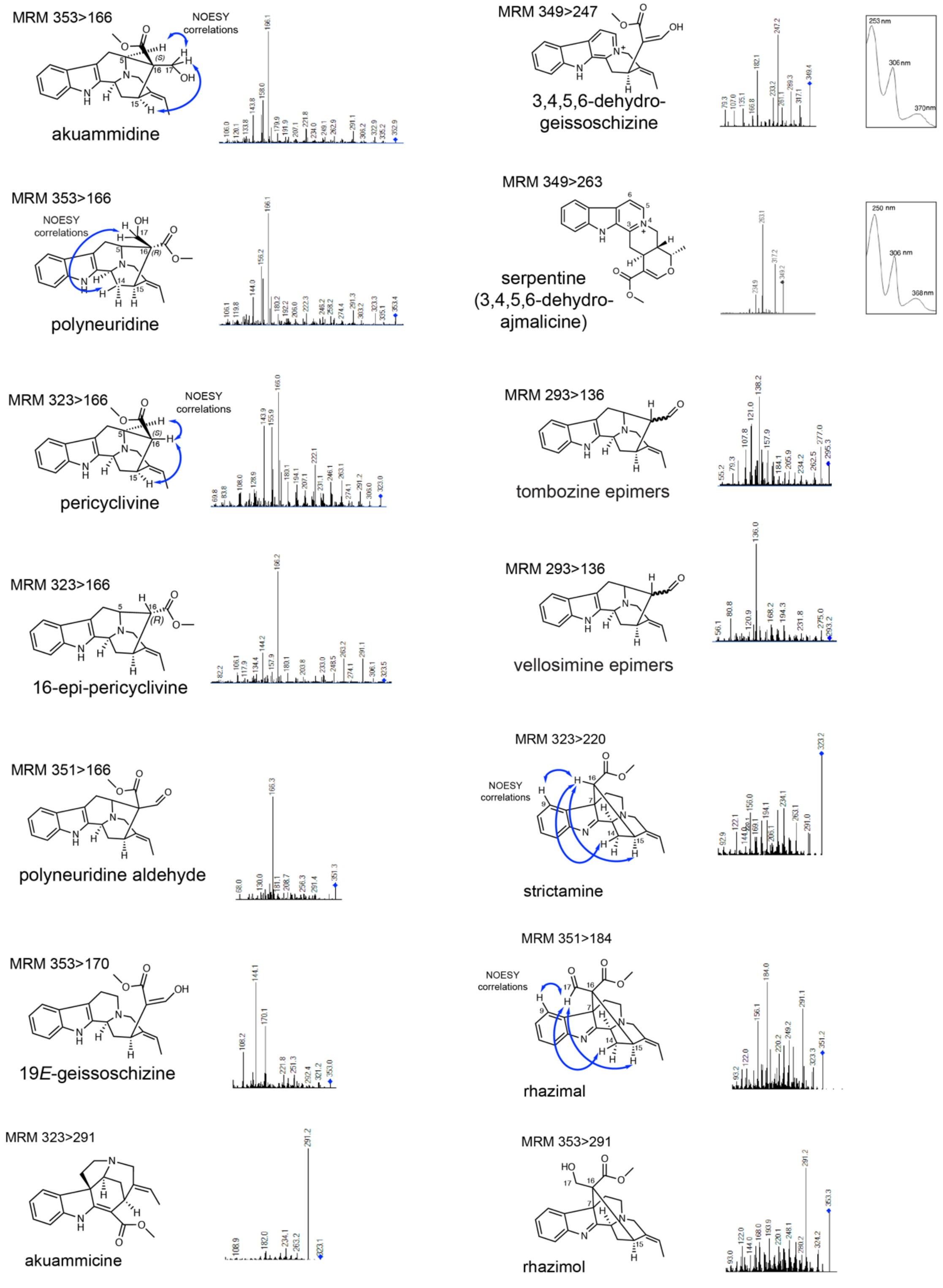

**Supplementary figure 2. MS/MS spectra for MIAs in this study.**
The MS/MS operated at 135 V for fragmentor and 30 V for collision energy. The by-product 3,4,5,6-dehydrogeissoschizine were concluded by comparing its UV absorption profile with that of serpentine, characteristic for fully aromatized beta-carboline structure. 16-epi-pericyclivine was produced by deformylating polyneuridine aldehyde with NAOH.

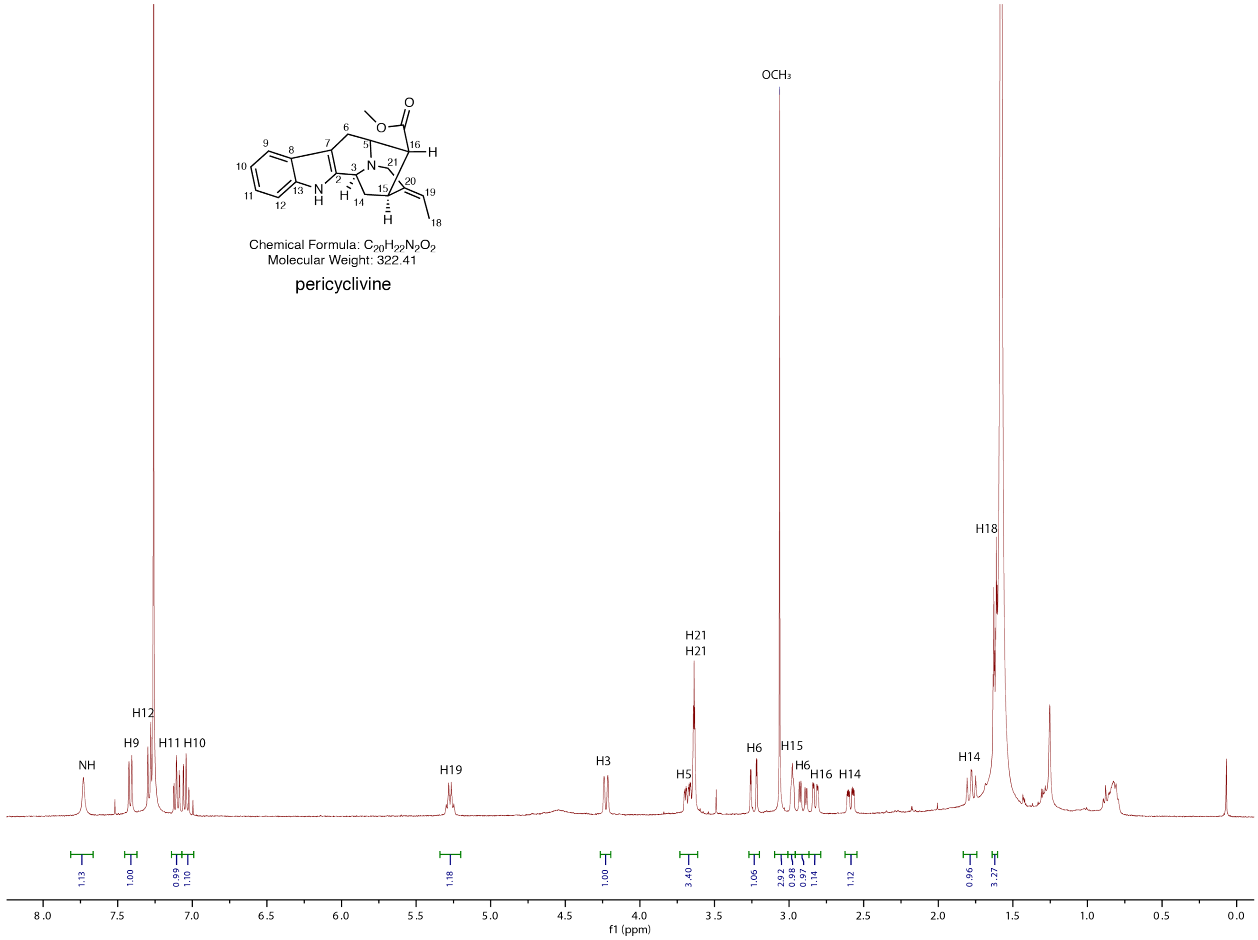

**Supplementary figure 3. [1]H NMR of pericyclivine in $CDCl_3$.**

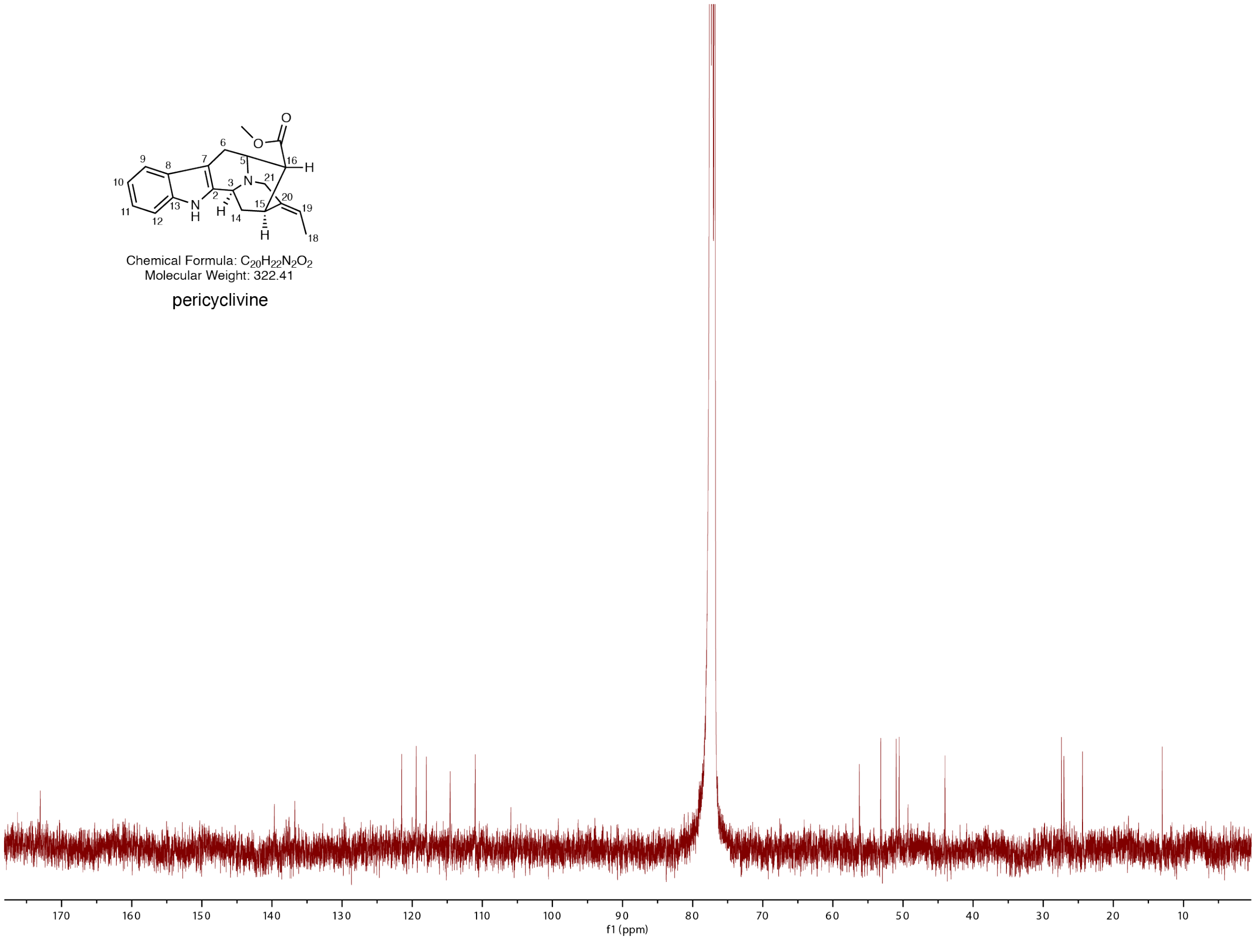

**Supplementary figure 4. $^{13}$C NMR of pericyclivine in $CDCl_3$.**

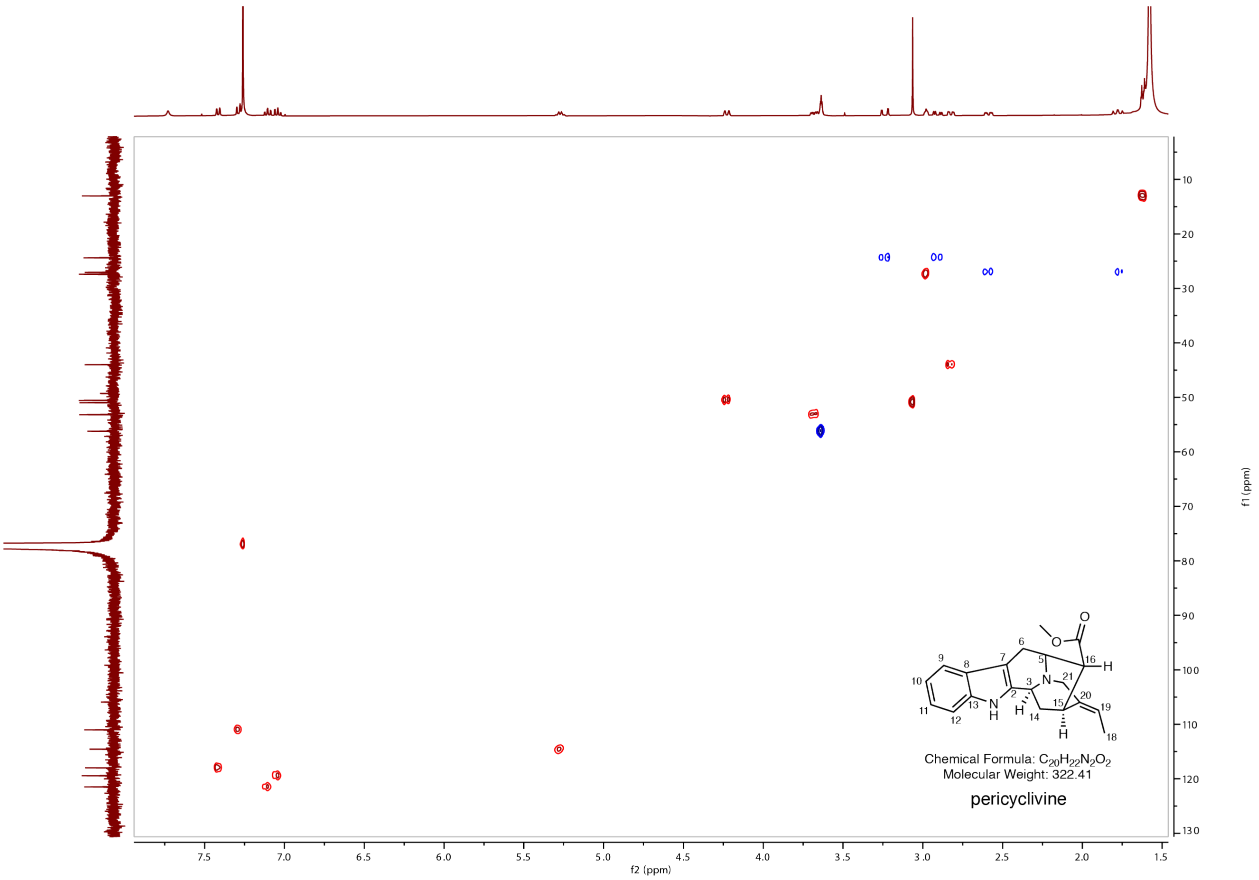

**Supplementary figure 5. HSQC NMR spectra of pericyclivine in $CDCl_3$.**

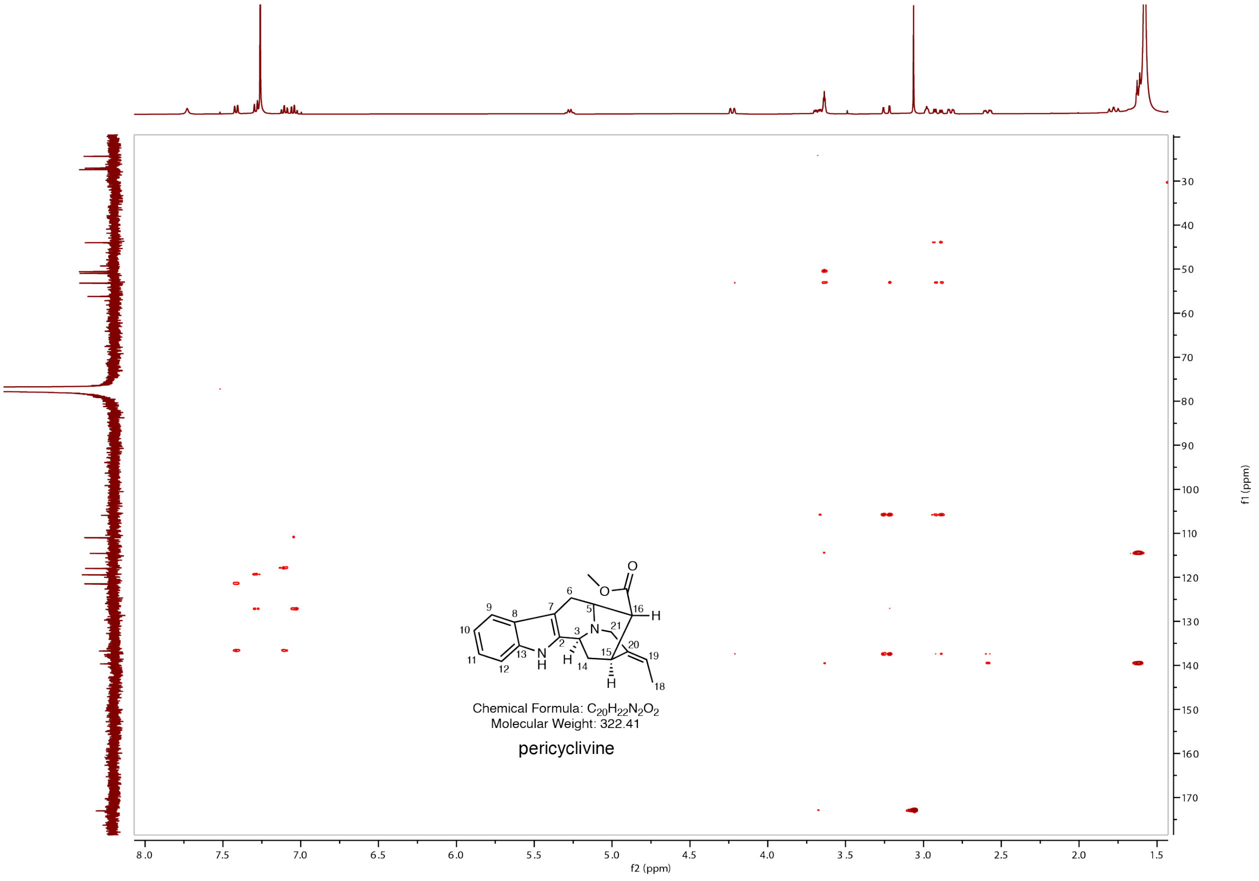

**Supplementary figure 6. HMBC NMR spectra of pericyclivine in $CDCl_3$.**

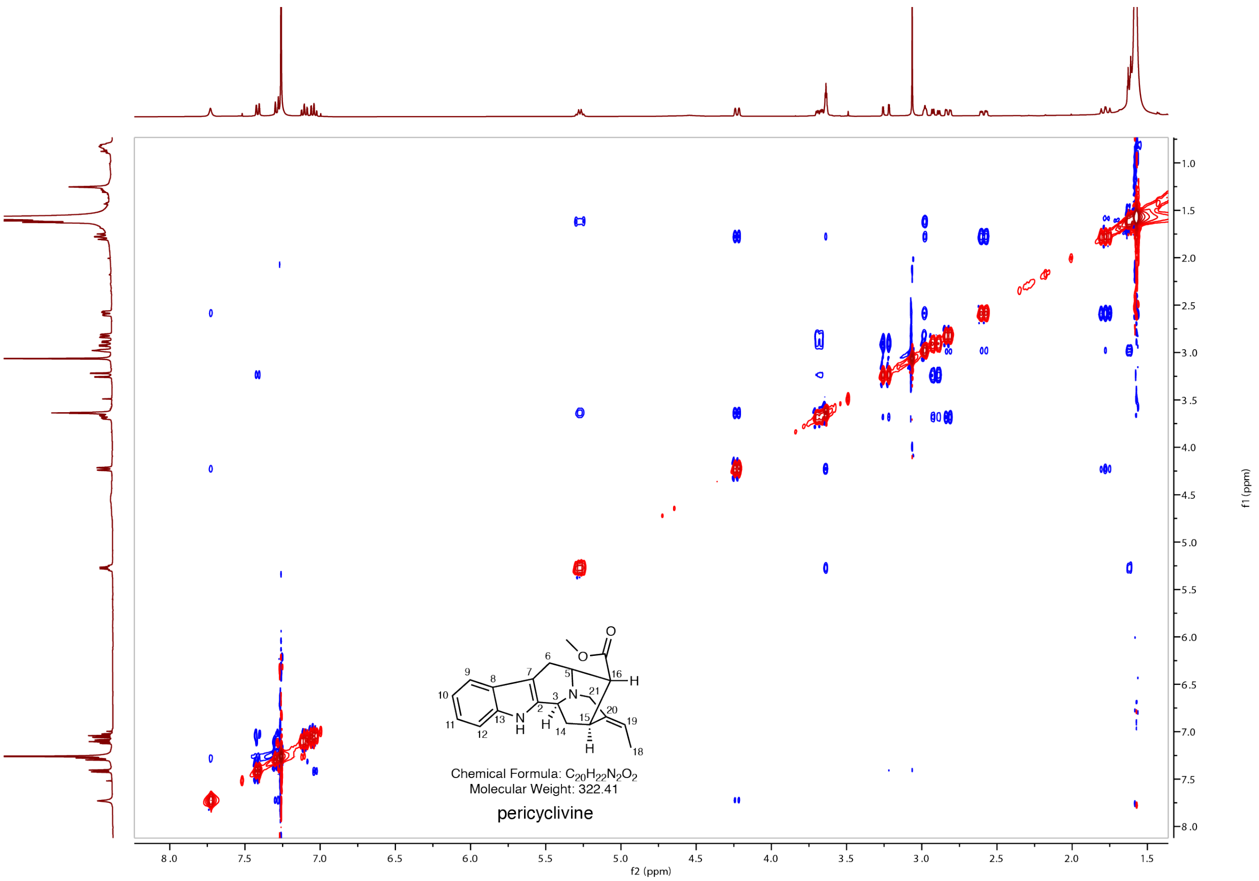

**Supplementary figure 7. NOESY NMR spectra of pericyclivine in $CDCl_3$.**

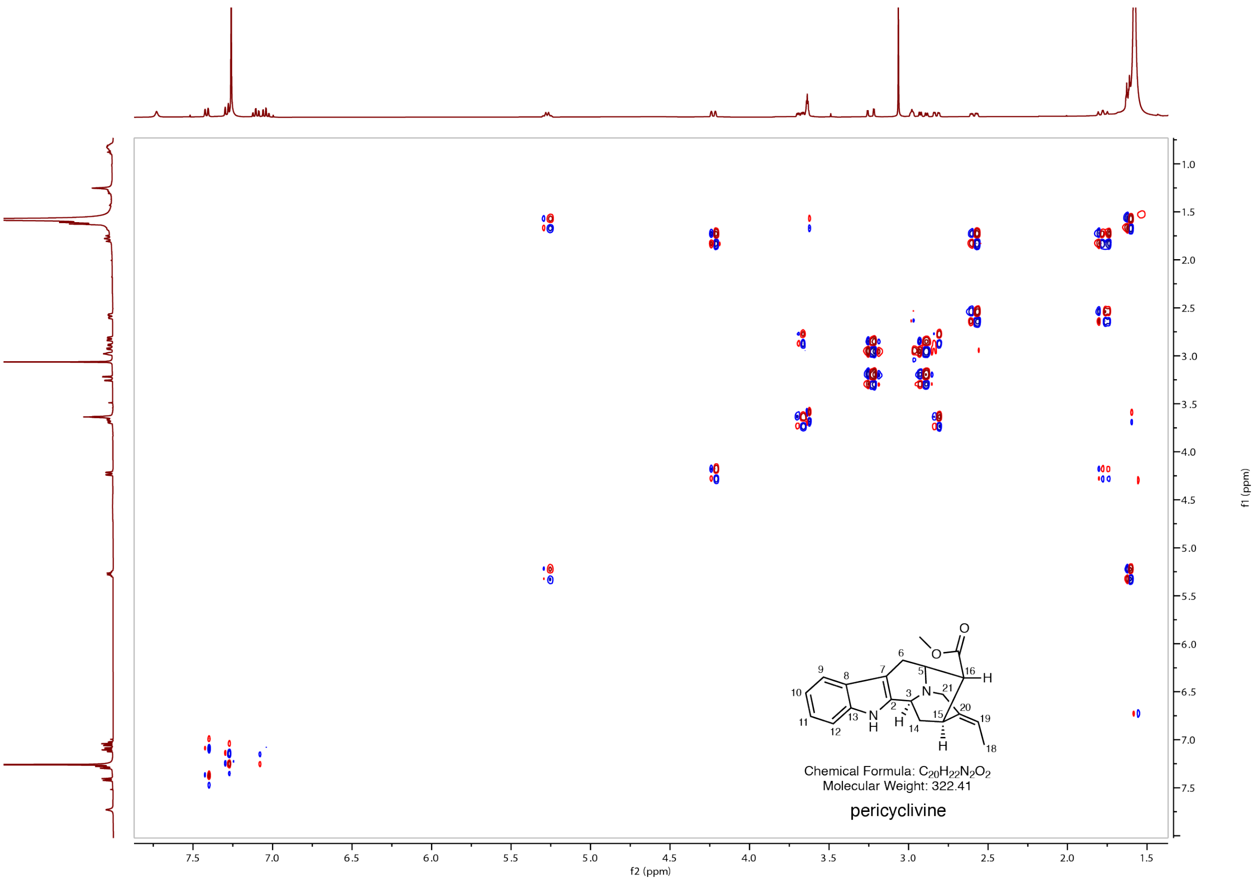

**Supplementary figure 8. COSY NMR spectra of pericyclivine in $CDCl_3$.**

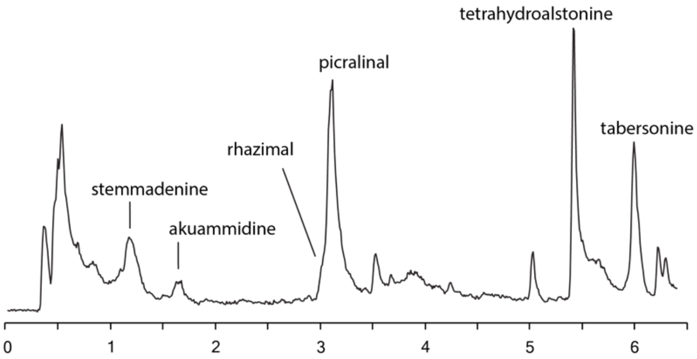

**Supplementary figure 9. Total ion chromatogram of *Amsonia tabernaemontana* leaf extract**

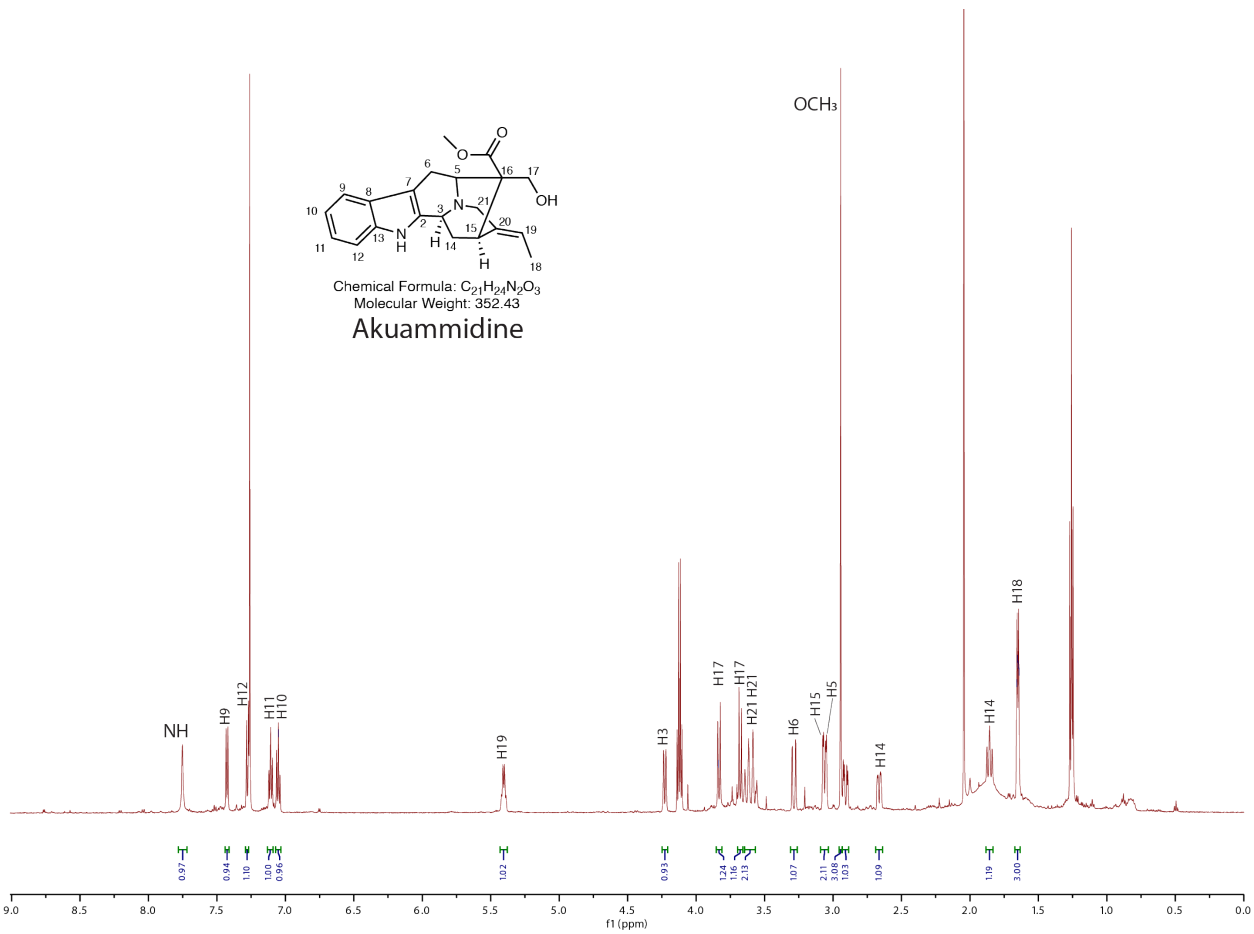

**Supplementary figure 10. [1]H NMR spectra of akuammidine in $CDCl_3$.**

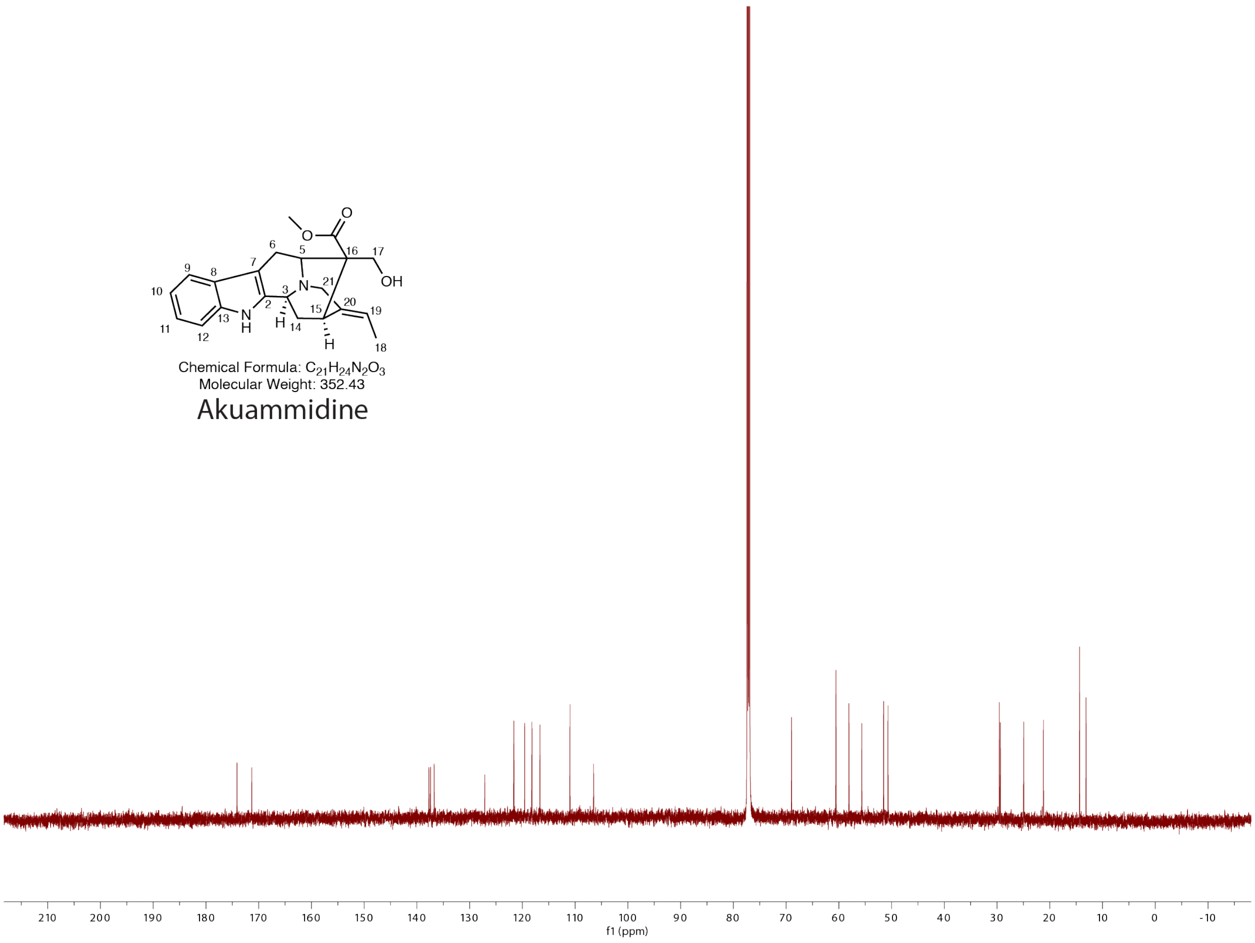

**Supplementary figure 11. $^{13}$C NMR spectra of akuammidine in $CDCl_3$.**

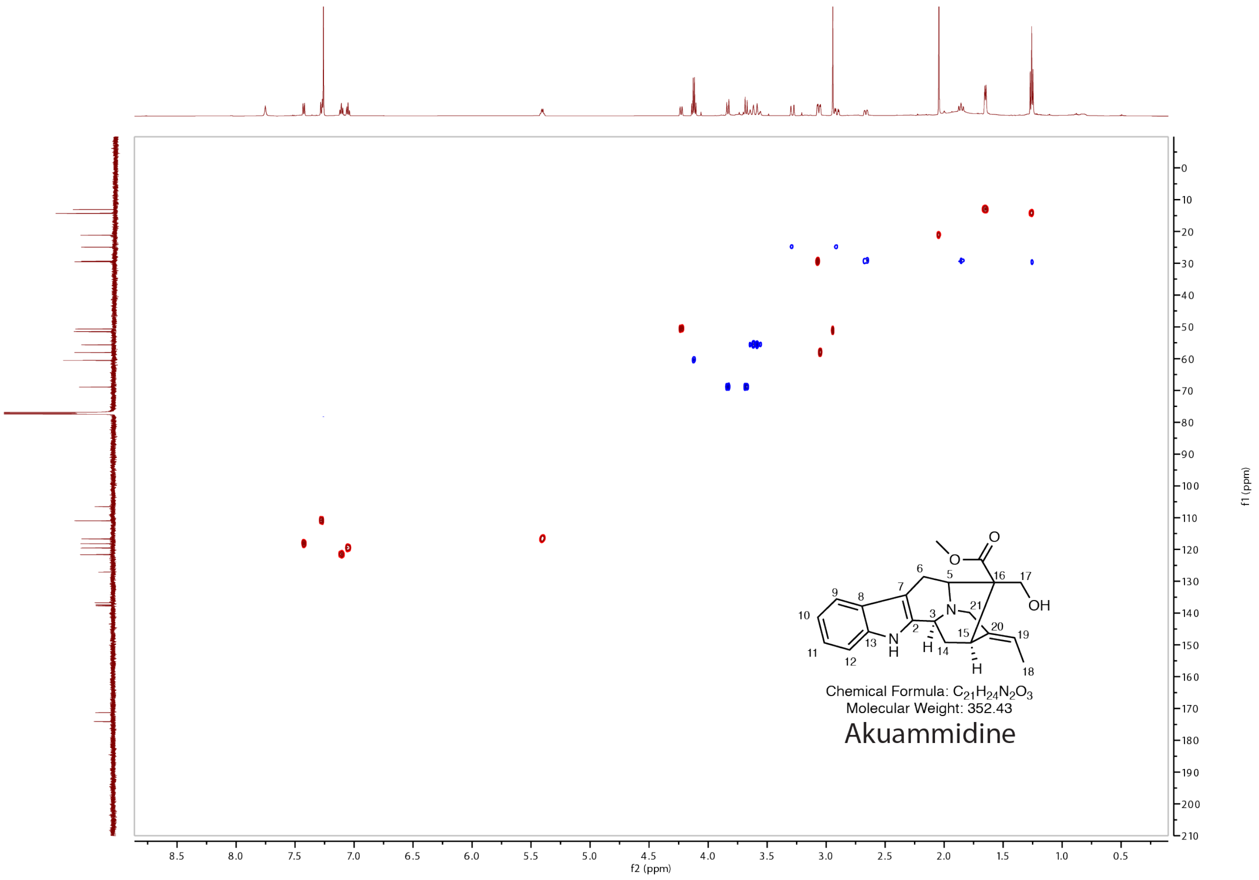

**Supplementary figure 12. HSQC NMR spectra of akuammidine in $CDCl_3$.**

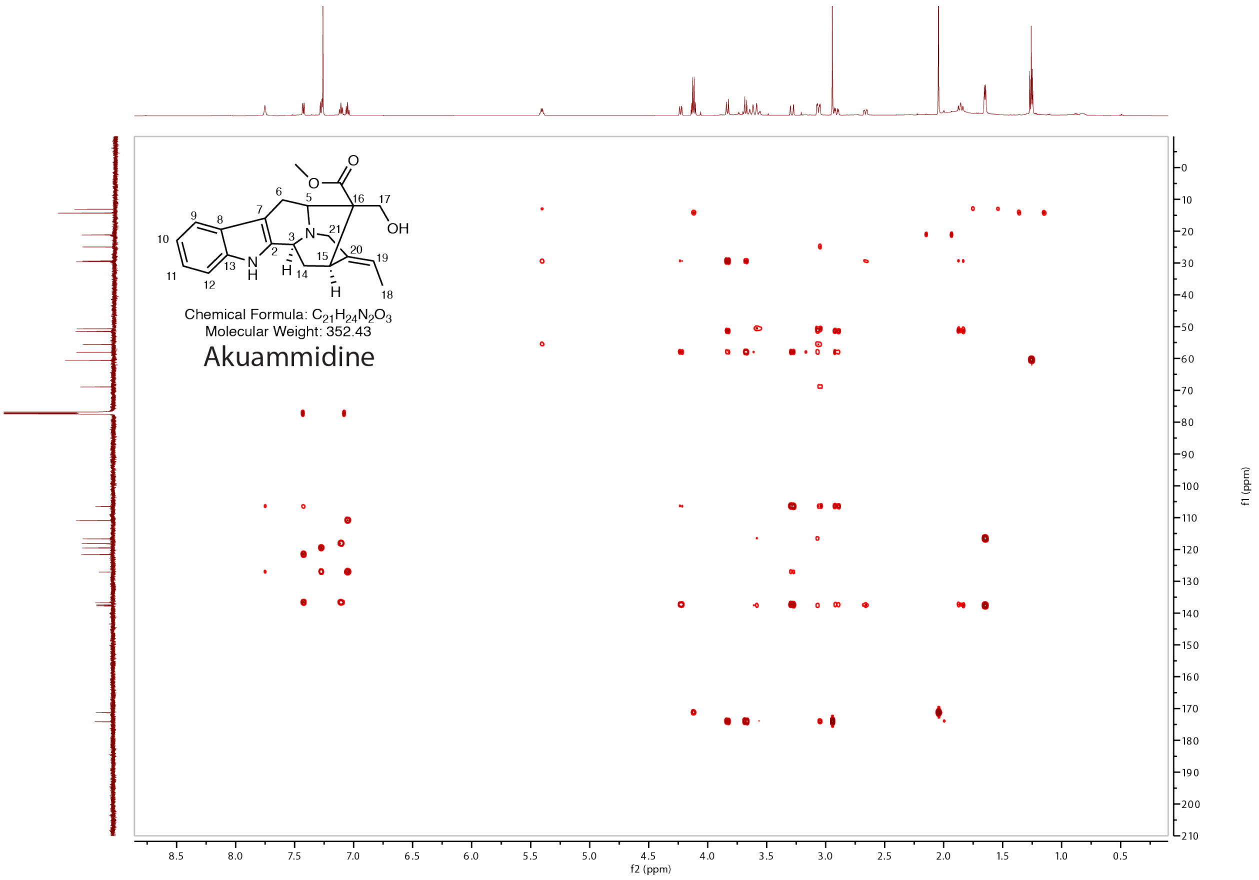

**Supplementary figure 13. HMBC NMR spectra of akuammidine in $CDCl_3$.**

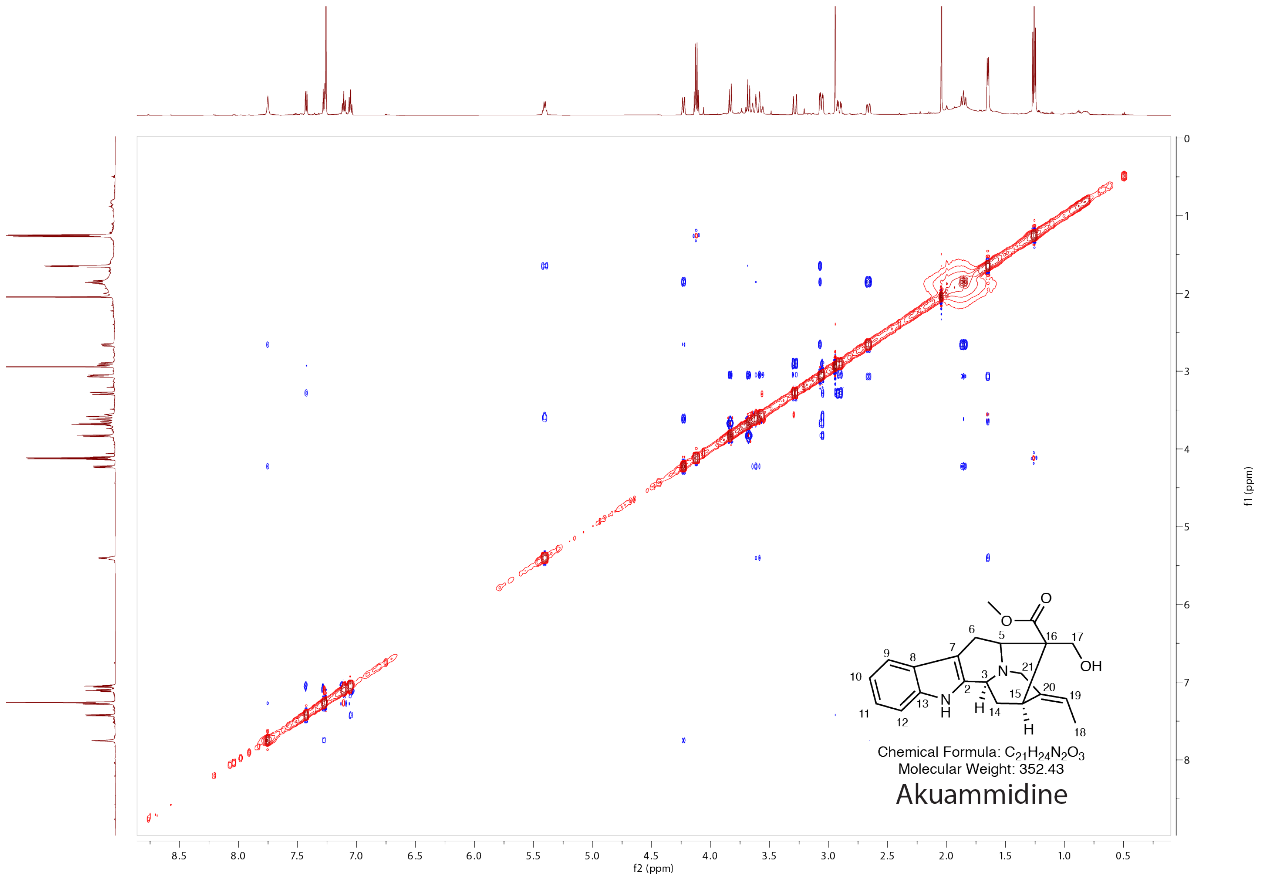

**Supplementary figure 14. NOESY NMR spectra of akuammidine in $CDCl_3$.**

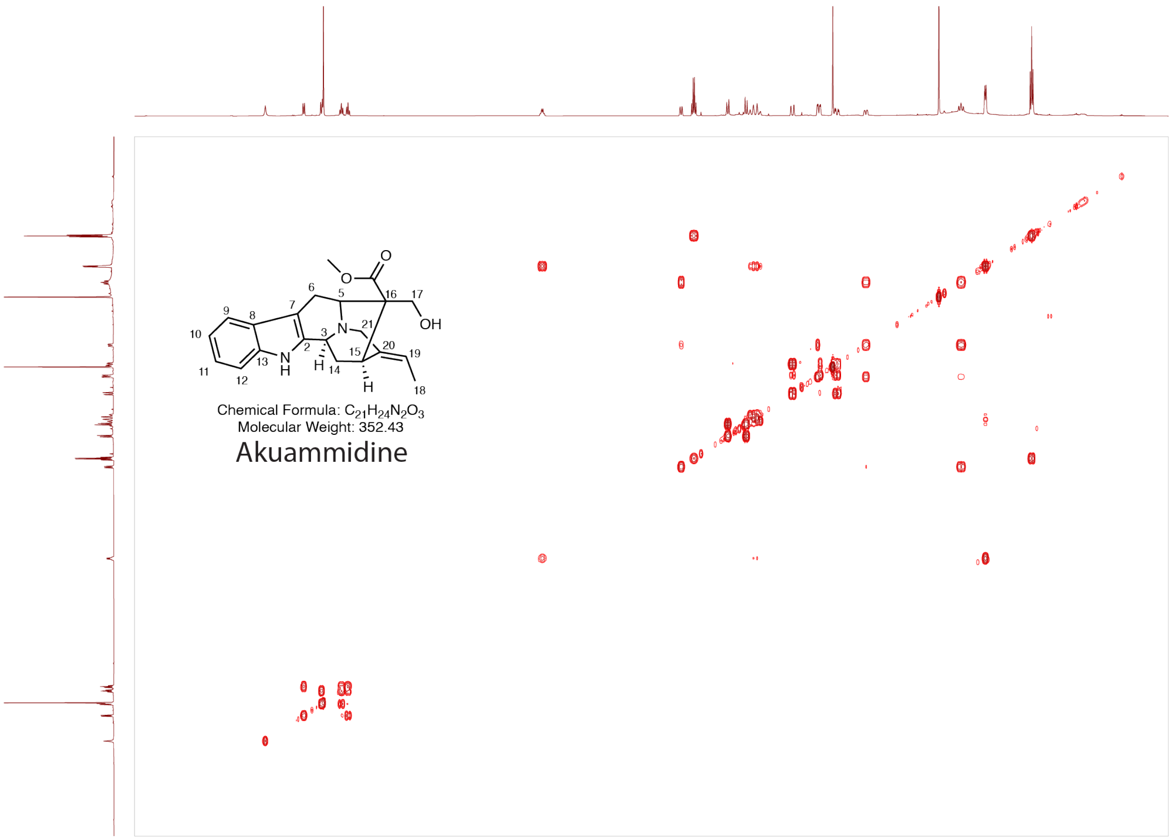

**Supplementary figure 15. COSY NMR spectra of akuammidine in $CDCl_3$.**

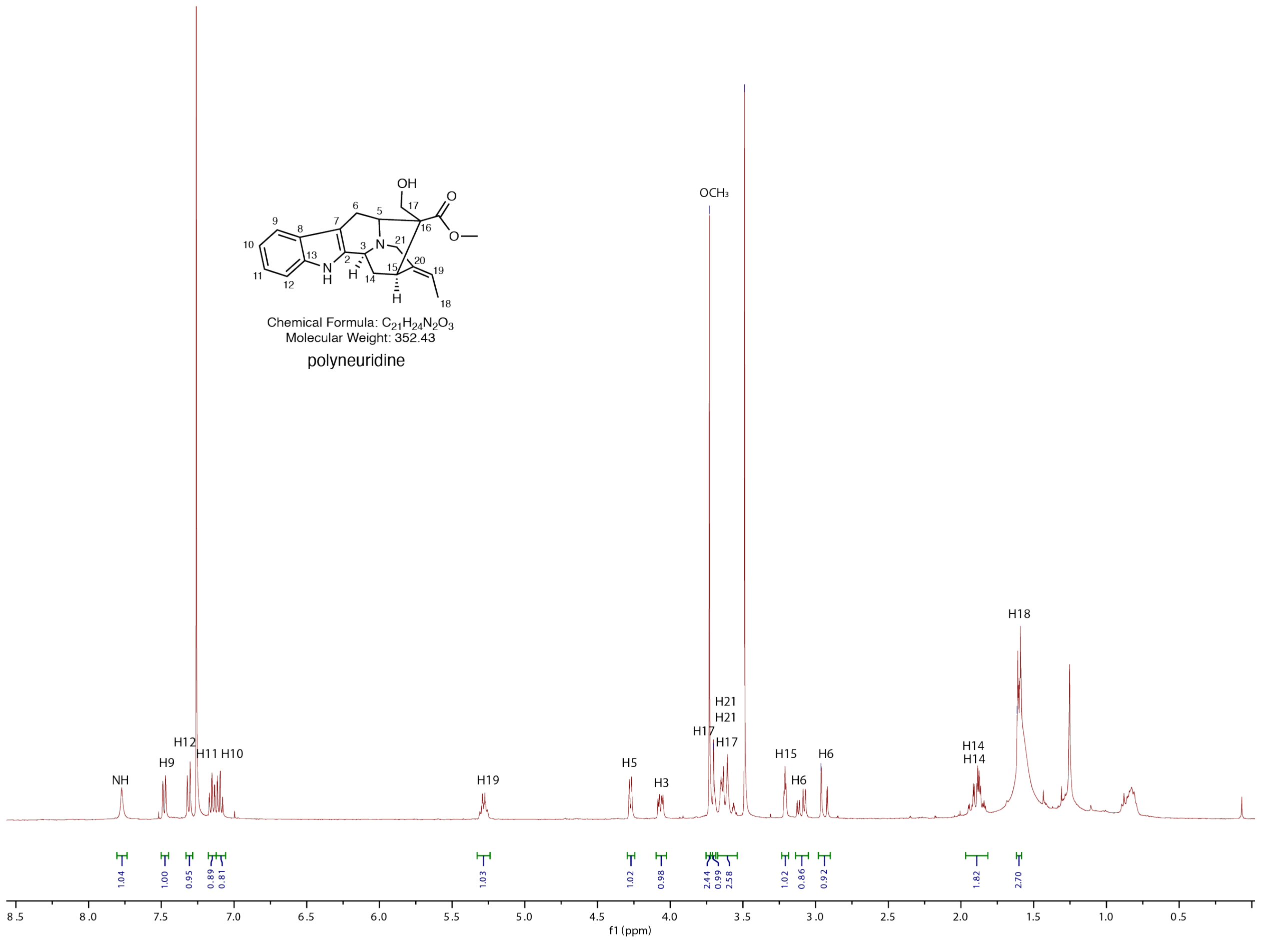

**Supplementary figure 16. $^1$H NMR spectra of polyneuridine in $CDCl_3$.**

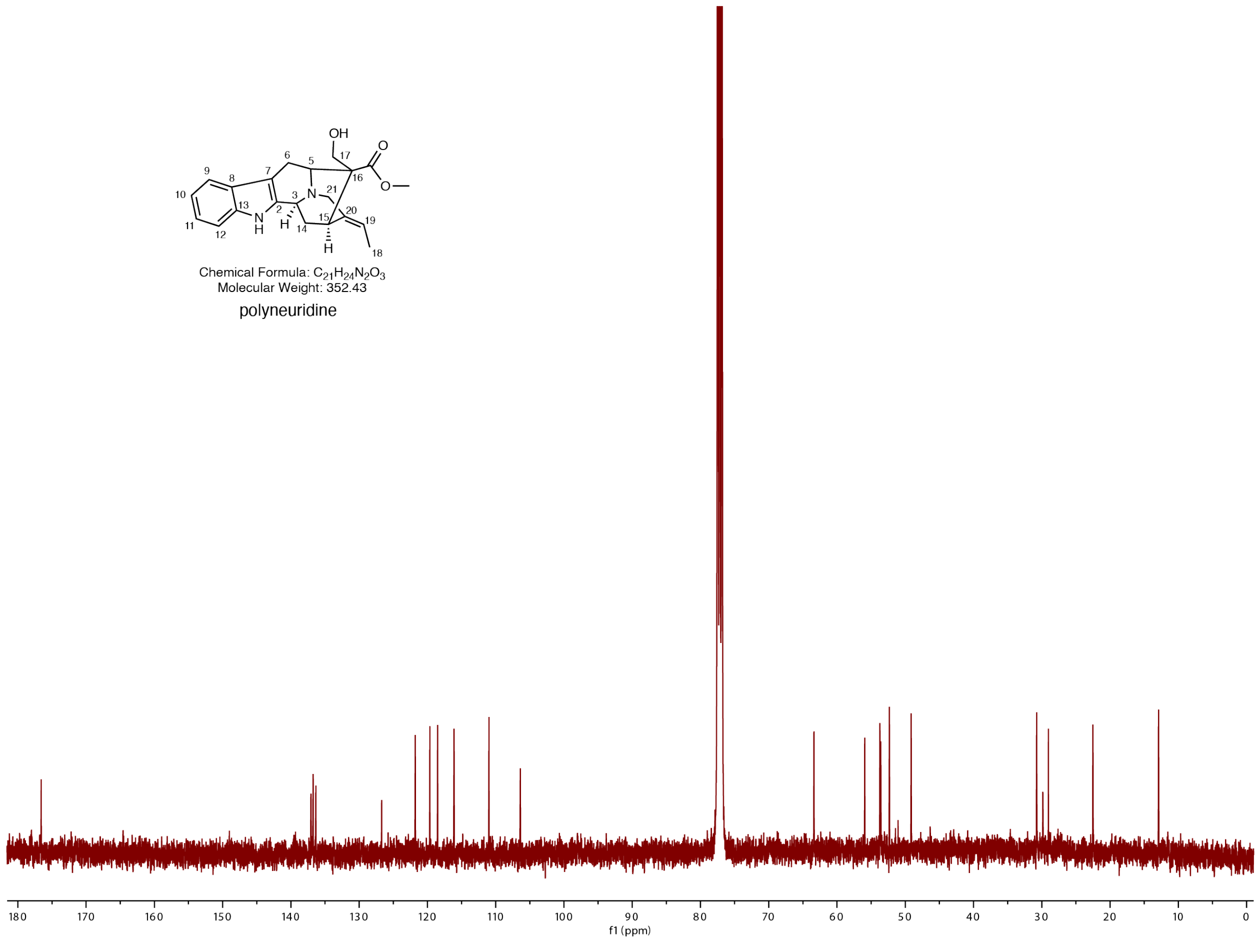

**Supplementary figure 17. $^{13}$C NMR spectra of polyneuridine in $CDCl_3$.**

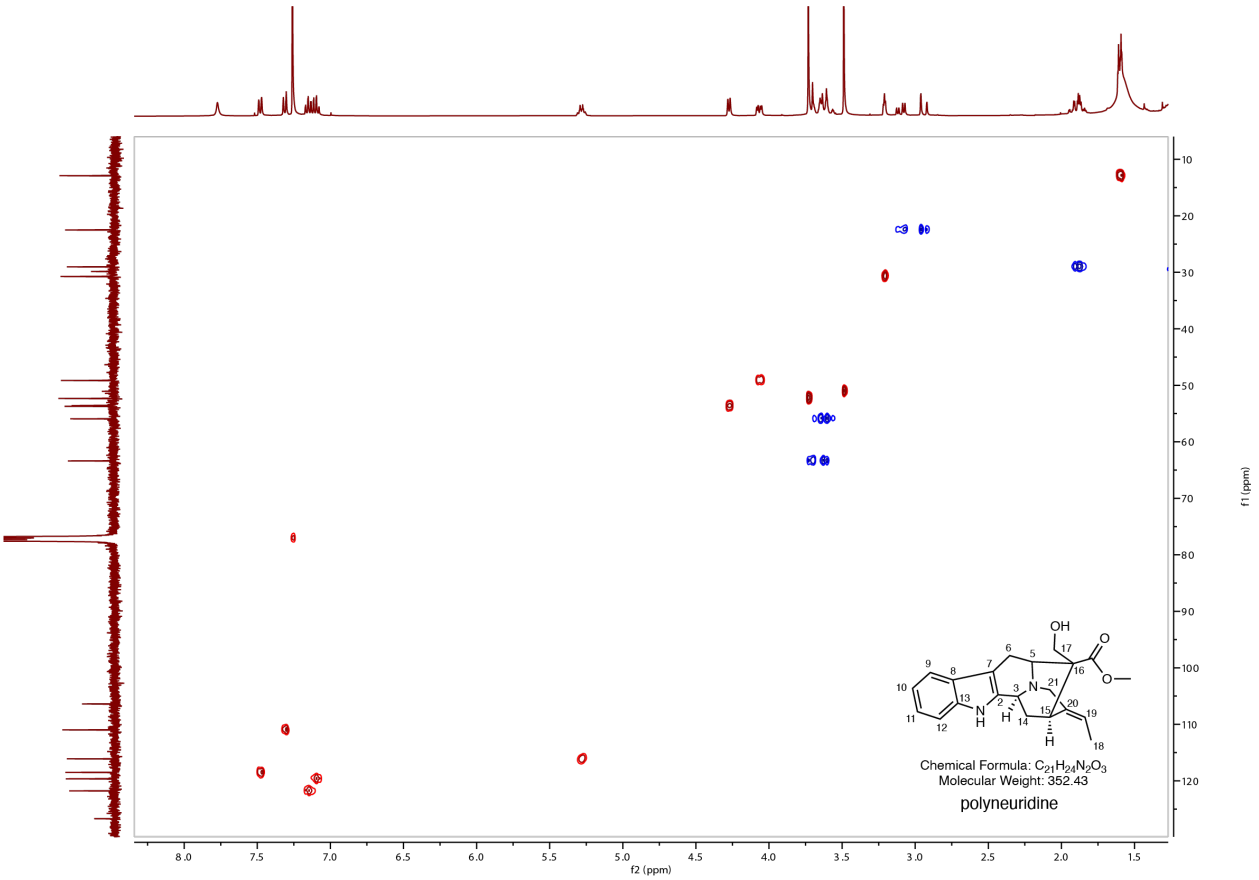

**Supplementary figure 18. HSQC NMR spectra of polyneuridine in $CDCl_3$.**

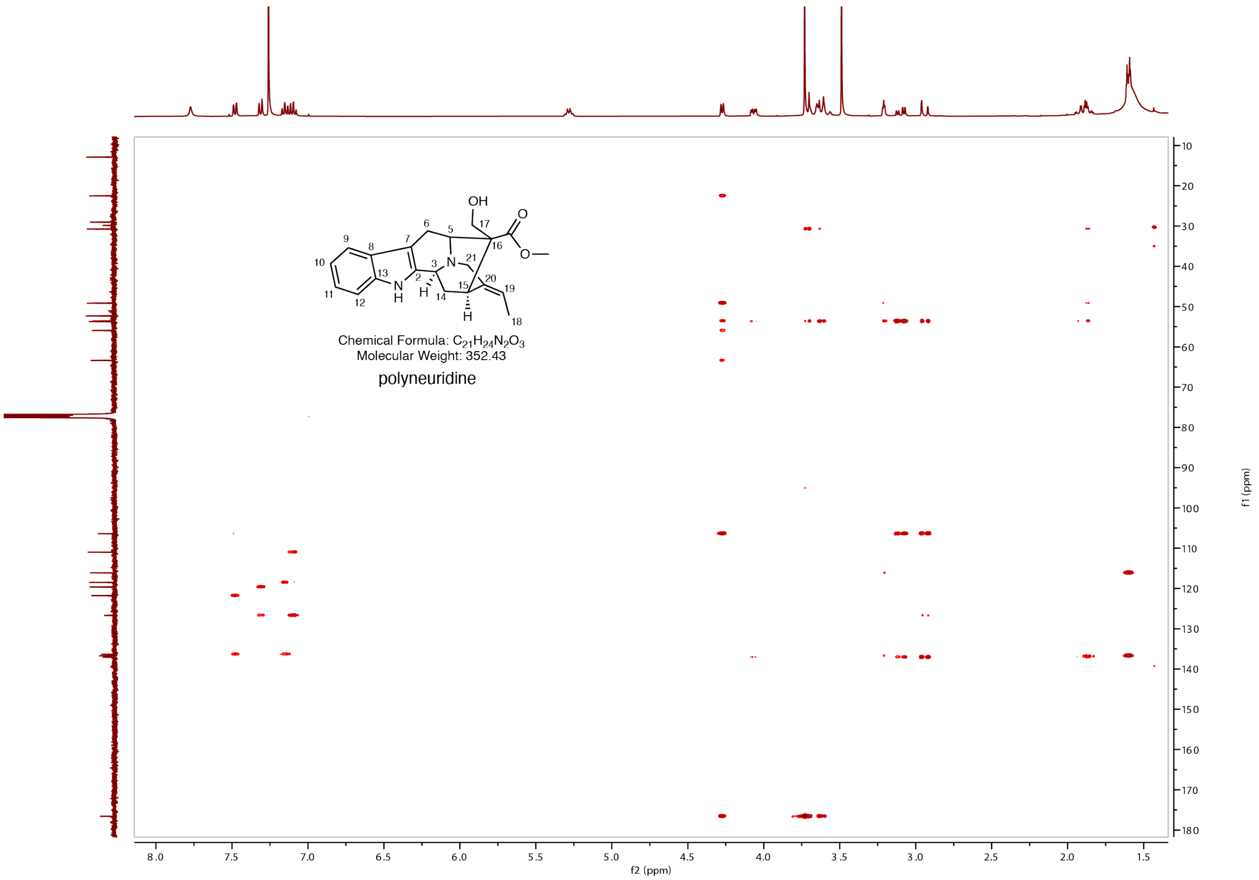

**Supplementary figure 19. HMBC NMR spectra of polyneuridine in $CDCl_3$.**

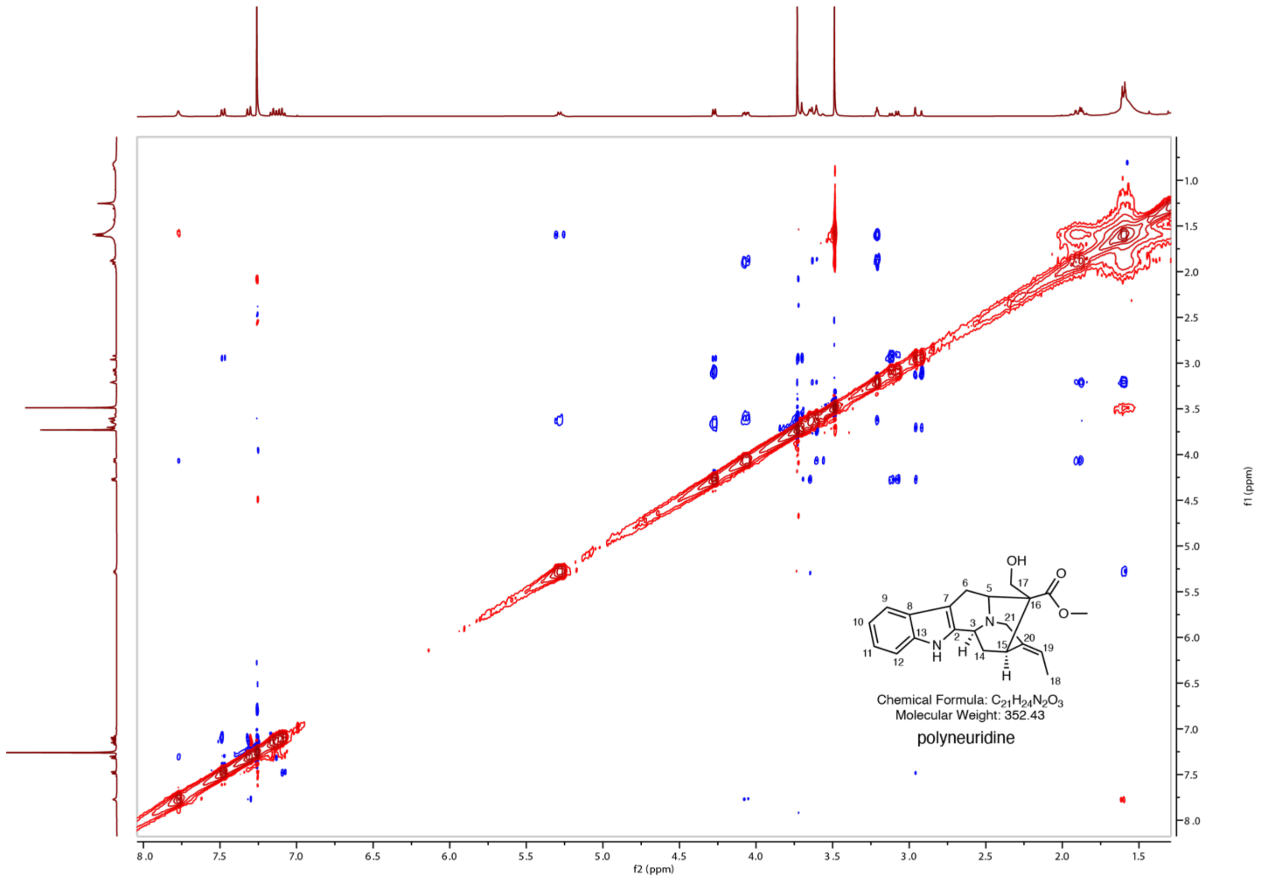

**Supplementary figure 20. NOESY NMR spectra of polyneuridine in $CDCl_3$.**

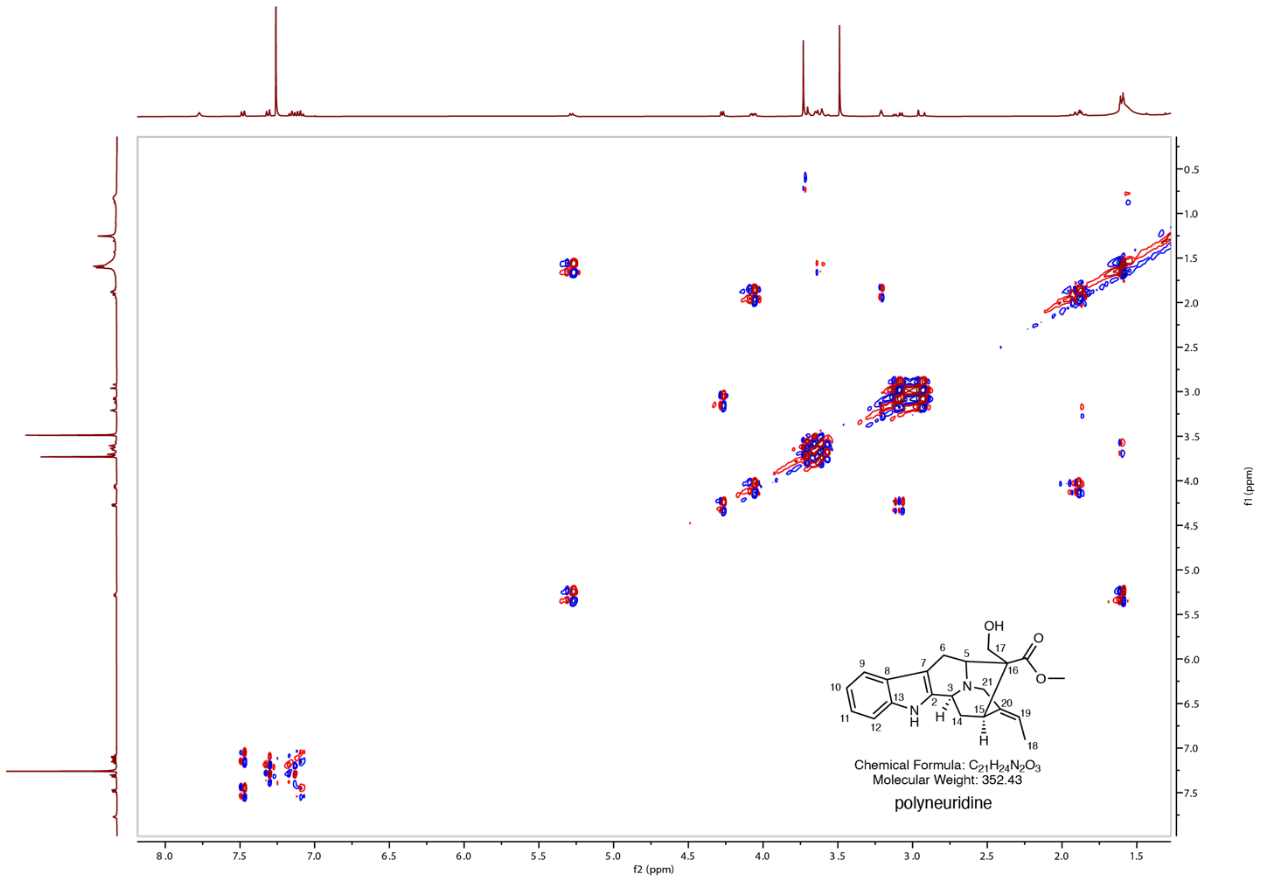

**Supplementary figure 21. COSY NMR spectra of polyneuridine in $CDCl_3$.**

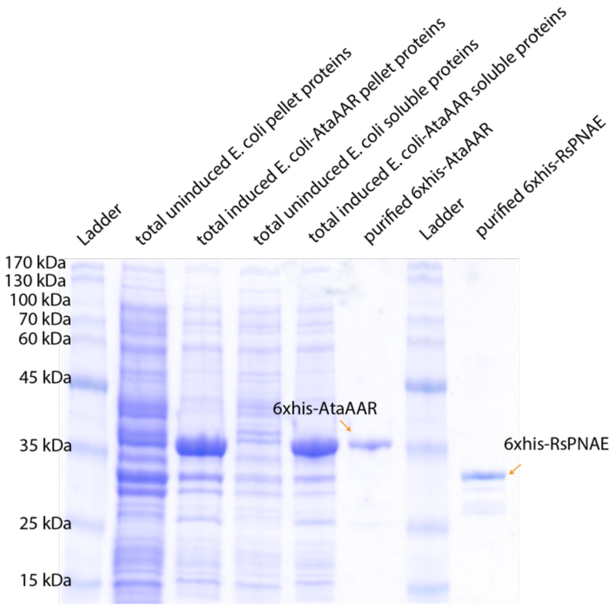

**Supplementary figure 22. SDS-PAGE showing the purifications of AtaAAR and RsPNAE.** The purified proteins migrated according to their theoretical molecular mass and the added 6x-histag: 6xhis-AtaAAR 40.7 kDa and 6xhis-PNAE 33.9 kDa.

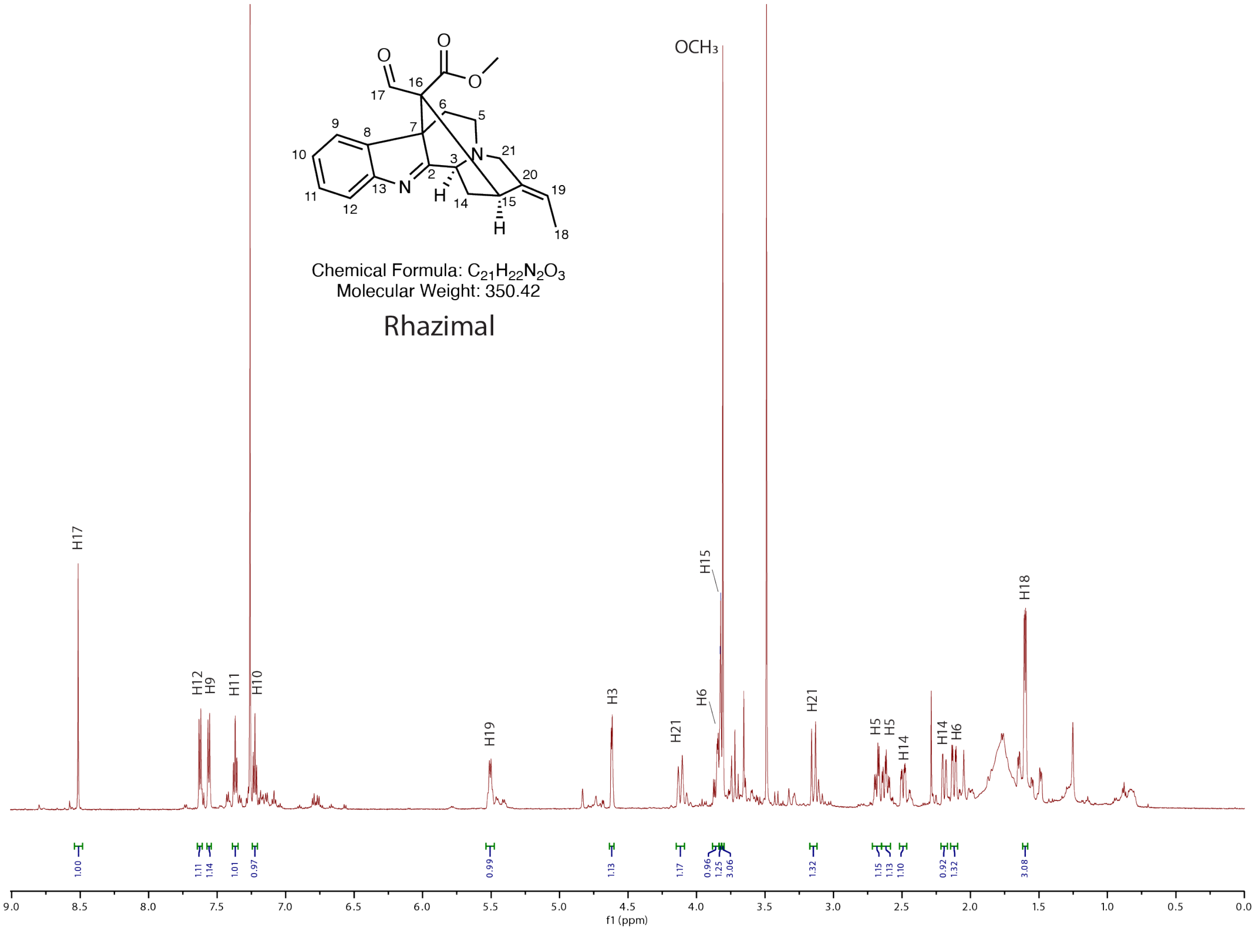

**Supplementary figure 23. [1]H NMR spectra of rhazimal in $CDCl_3$.**

[1]H NMR (600 MHz, CDCl3) δ 8.51 (s, 1H, H-17), 7.63 (d, J = 7.8 Hz, 1H, H-12), 7.56 (d, J = 7.5 Hz, 1H, H-9), 7.37 (dt, J = 7.6, 1.3 Hz, 1H, H-11), 7.22 (dt, J = 7.5, 1.1 Hz, 1H, H-10), 5.51 (q, J = 7.5 Hz, 1H, H-19), 4.62 (d, J = 4.9 Hz, 1H, H-3), 4.12 (d, J = 17.7 Hz, 1H, H-21), 3.85 (dd, *J* = 14.8, 5.7 Hz, 1H, H-6), 3.83 (s, 1H, H-15), 3.81 (s, 3H, $OCH_3$), 3.15 (d, J = 17.2 Hz, 1H, H-21), 2.68 (dd, J = 14.3, 5.8 Hz, 1H, H-5), 2.62 (td, J = 14.2, 4.5 Hz, 1H, H-5), 2.49 (ddd, J = 14.7, 5.1, 2.4 Hz, 1H, H-14), 2.19 (d, J = 14.5 Hz, 1H, H-14), 2.12 (dd, J = 15.0, 4.3 Hz, 1H, H-6), 1.60 (dd, J = 7.2, 2.6 Hz, 3H, H-18).

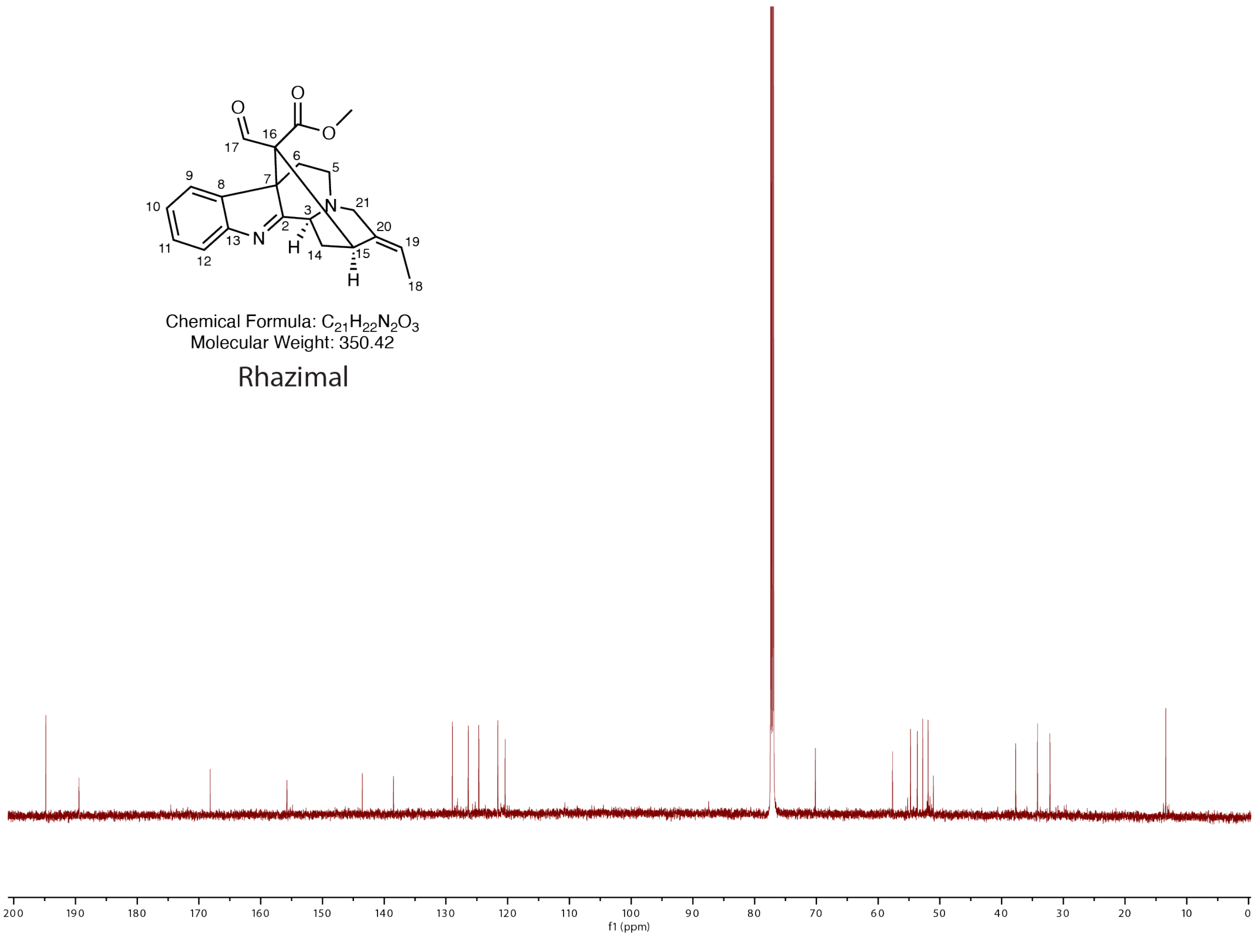

**Supplementary figure 24. $^{13}$C NMR spectra of rhazimal in $CDCl_3$.**

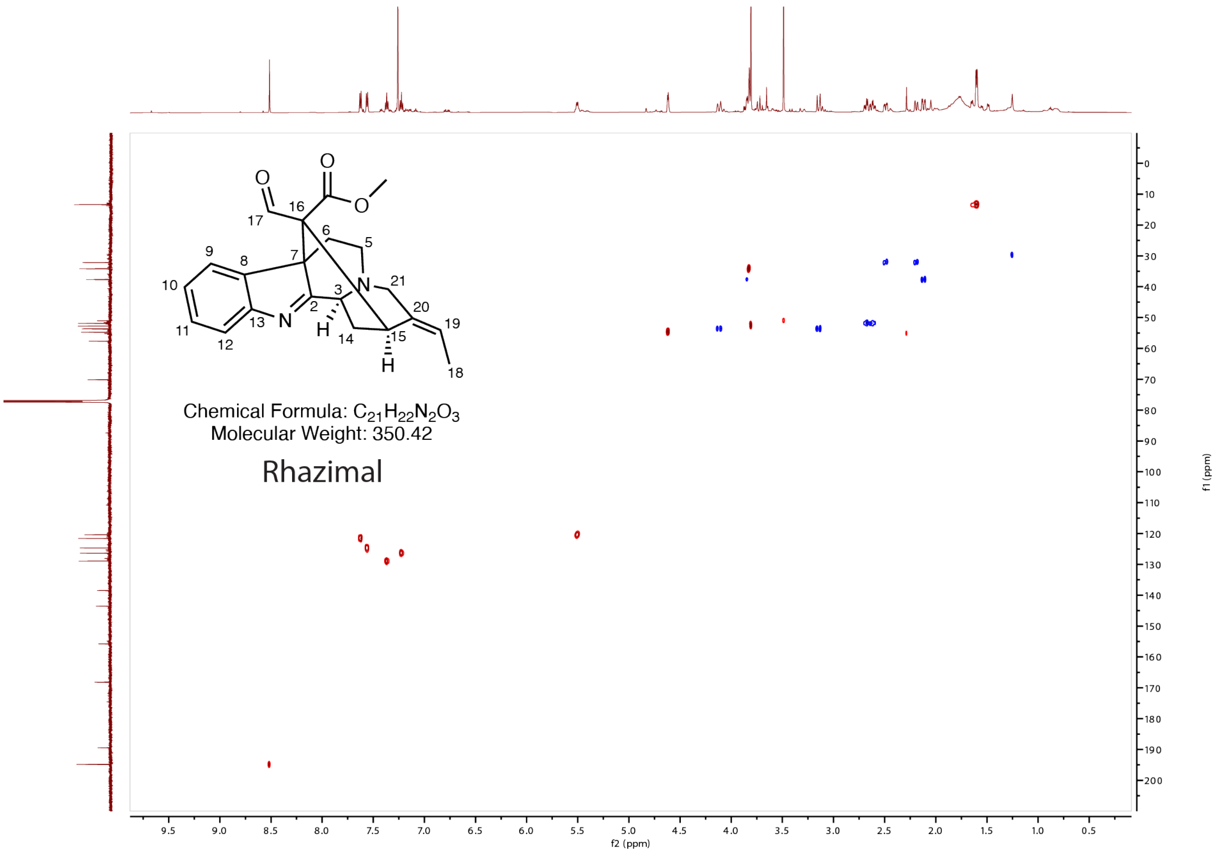

**Supplementary figure 25. HSQC NMR spectra of rhazimal in $CDCl_3$.**

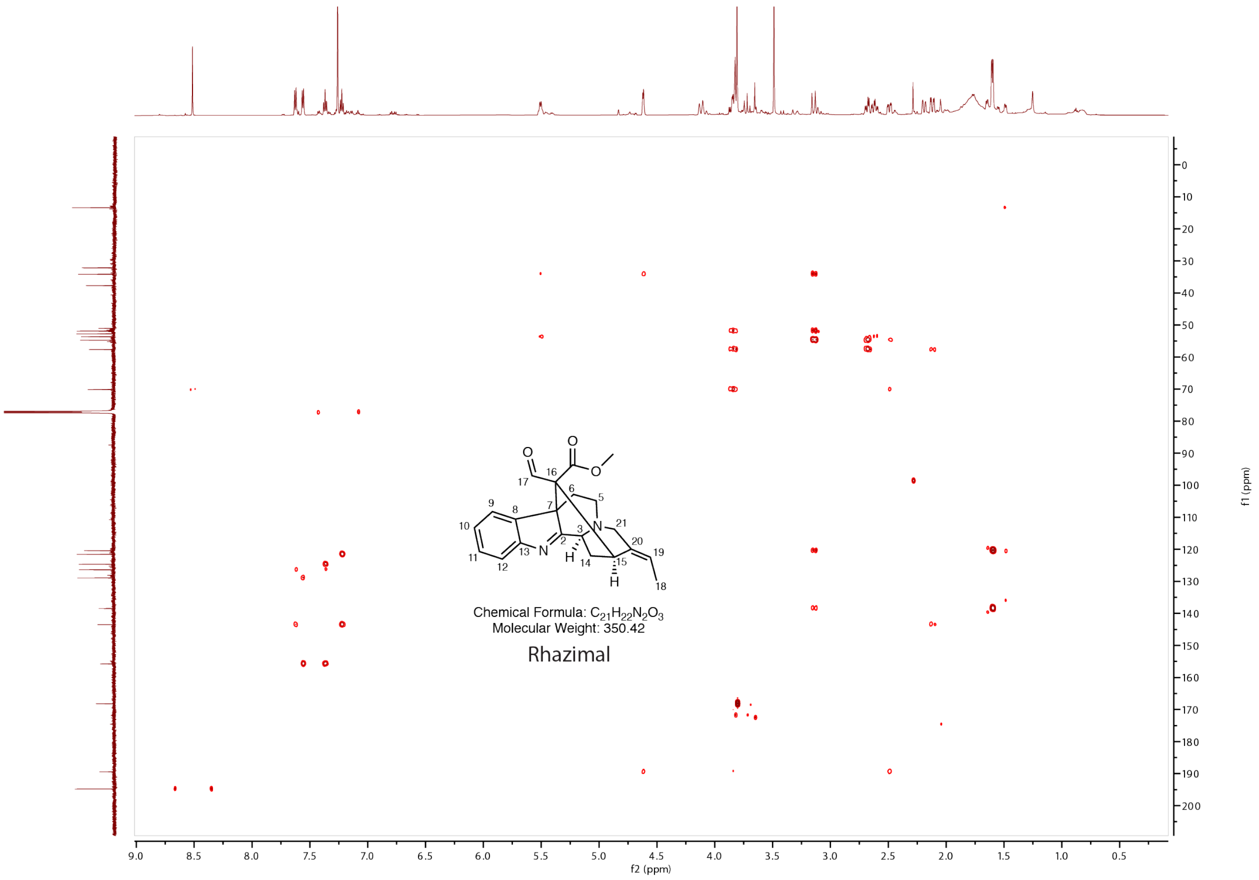

**Supplementary figure 26. HMBC NMR spectra of rhazimal in $CDCl_3$.**

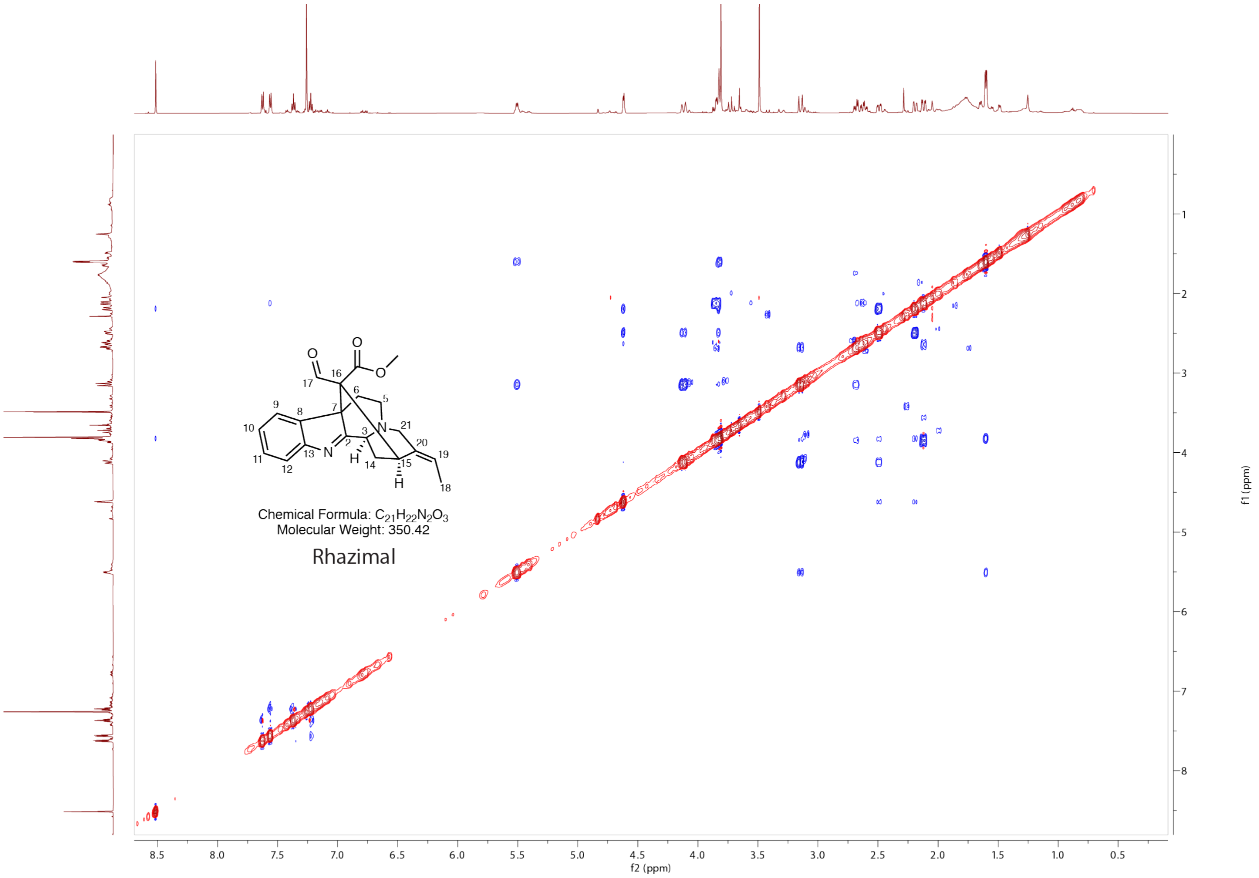

**Supplementary figure 27. NOESY NMR spectra of rhazimal in $CDCl_3$.**

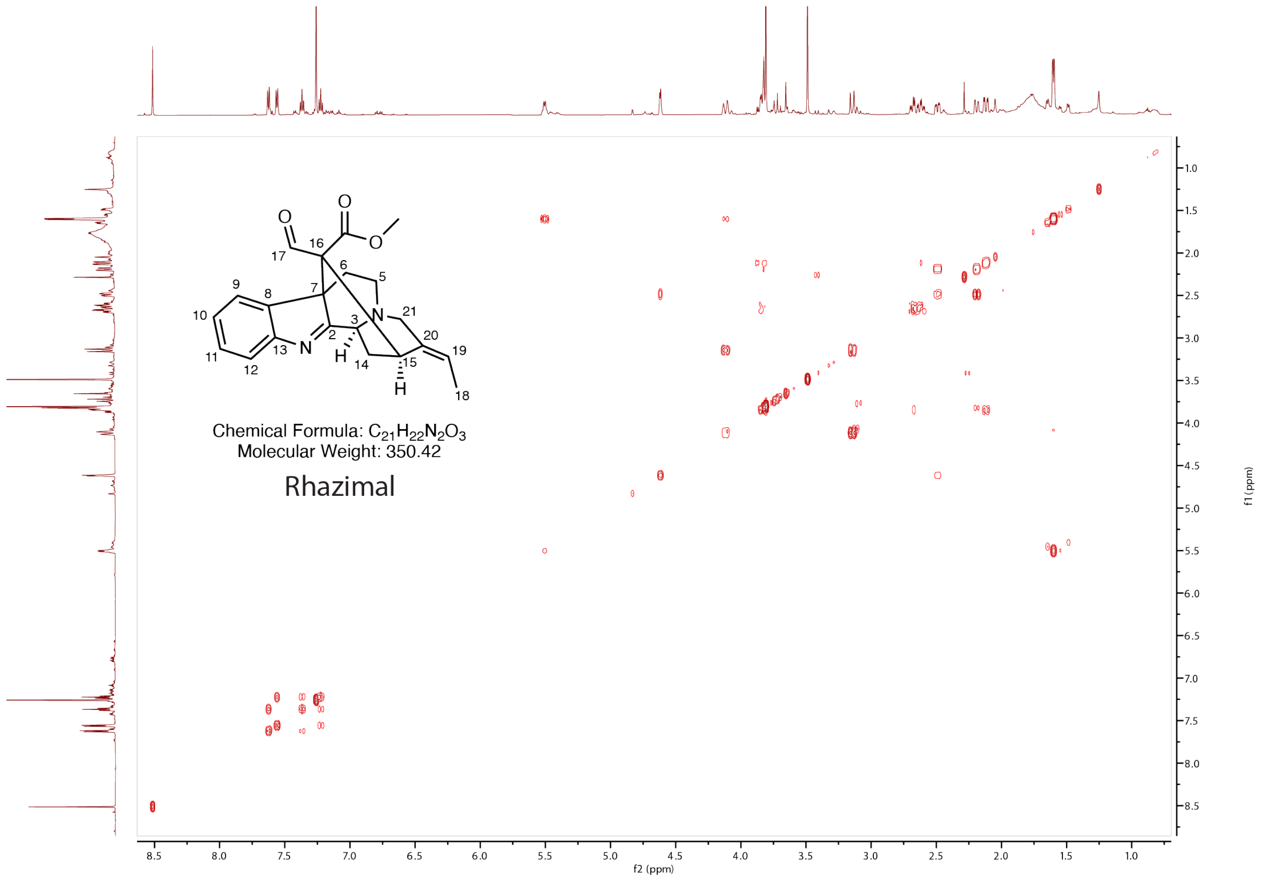

**Supplementary figure 28. COSY NMR spectra of rhazimal in $CDCl_3$.**

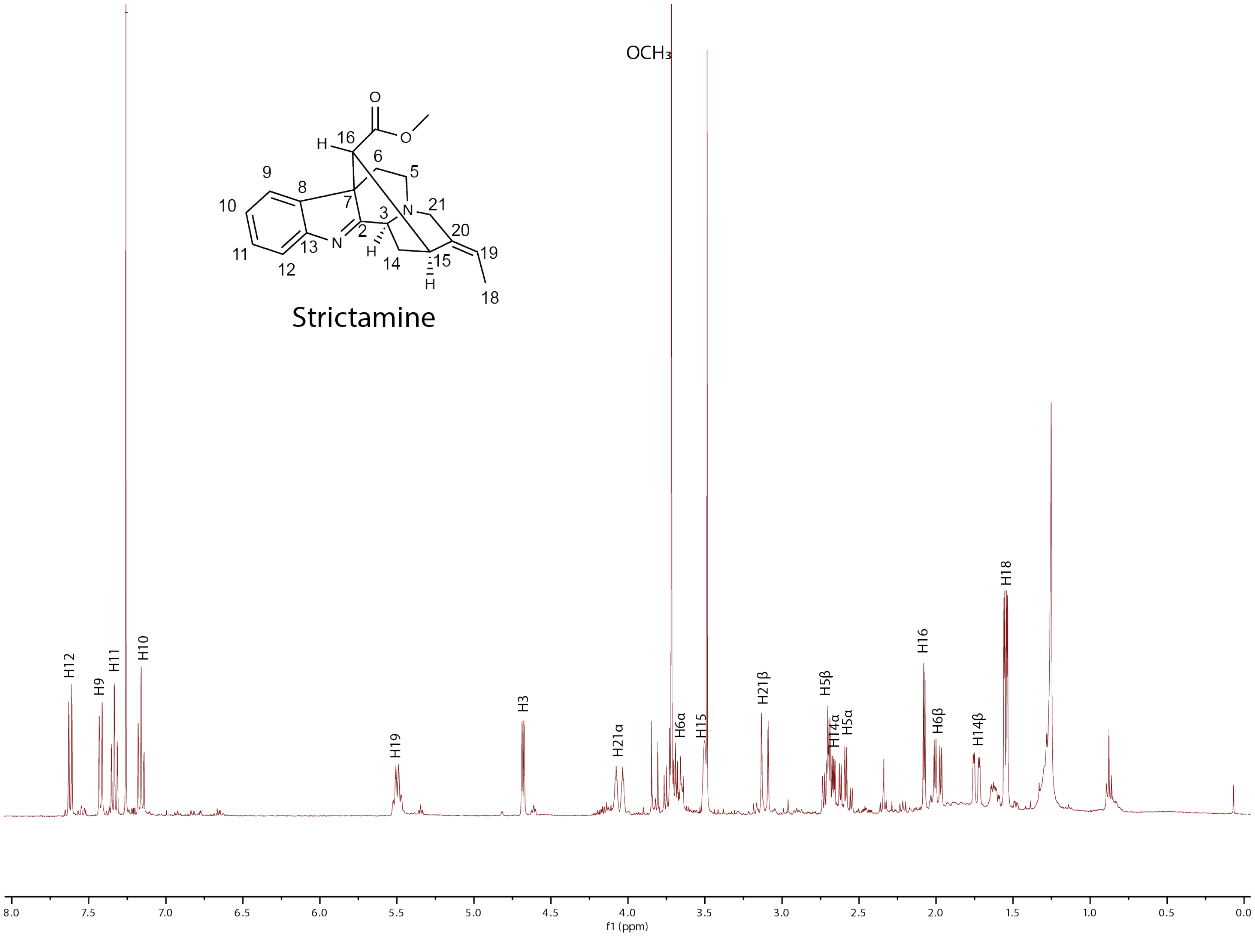

**Supplementary figure 29. $^{1}$H NMR spectra of strictamine in $CDCl_3$.**

$^{1}$H NMR (400 MHz, $CDCl_3$) δ 7.62 (d, *J* = 7.7, 1H, H-12), 7.42 (d, *J* = 7.5, 1H, H-9), 7.33 (dt, *J* = 7.6, 1.3 Hz, 1H, H-11), 7.16 (dt, *J* = 7.5, 1.1 Hz, 1H, H-10), 5.50 (q, *J* = 7.2 Hz, 1H, H-19), 4.68 (d, *J* = 5.3 Hz, 1H, H-3), 4.05 (dt, *J* = 17.4, 2.2 Hz, 1H, H-21), 3.72 (s, 3H, $OCH_3$), 3.72 (m, 1H, H-6), 3.50 (s, 1H, H-15), 3.11 (d, *J* = 16.9 Hz, 1H, H-21), 2.71 (dd, *J* = 10.8, 5.6 Hz, 1H, H-5), 2.68 (ddd, *J* = 11.6, 5.2, 2.8 Hz, 1H, H-4), 2.59 (ddd, *J* = 17.8, 14.3, 5.0 Hz, 1H, H-5), 2.08 (d, *J* = 3.9 Hz, 1H, H-16), 1.99 (dd, *J* = 14.7, 4.9 Hz, 1H, H-6), 1.74 (dd, *J* = 13.8, 3.4 Hz, 1H, H-14), 1.55 (dd, *J* = 7.1, 2.6 Hz, 3H, H-18).

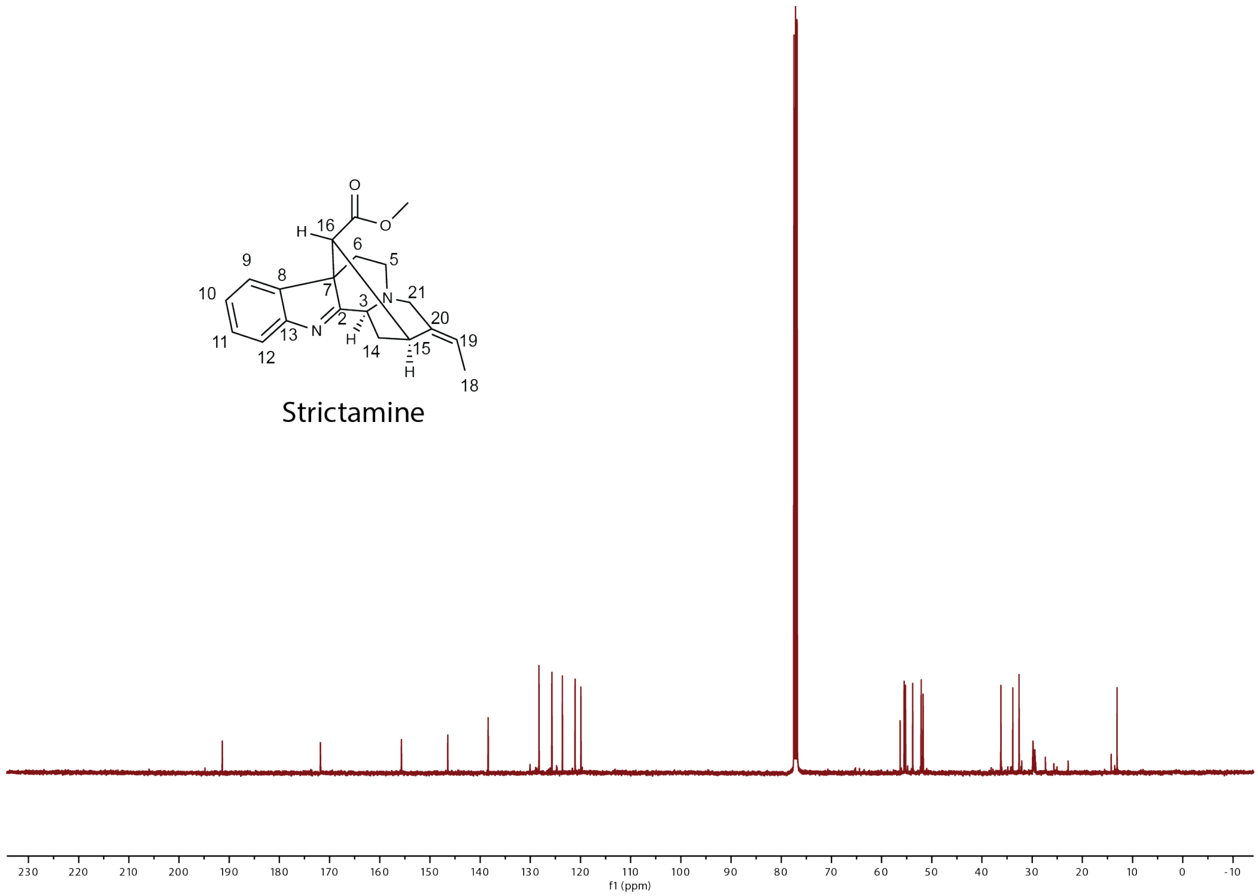

**Supplementary figure 30. $^{13}C$ NMR spectra of strictamine in $CDCl_3$.**

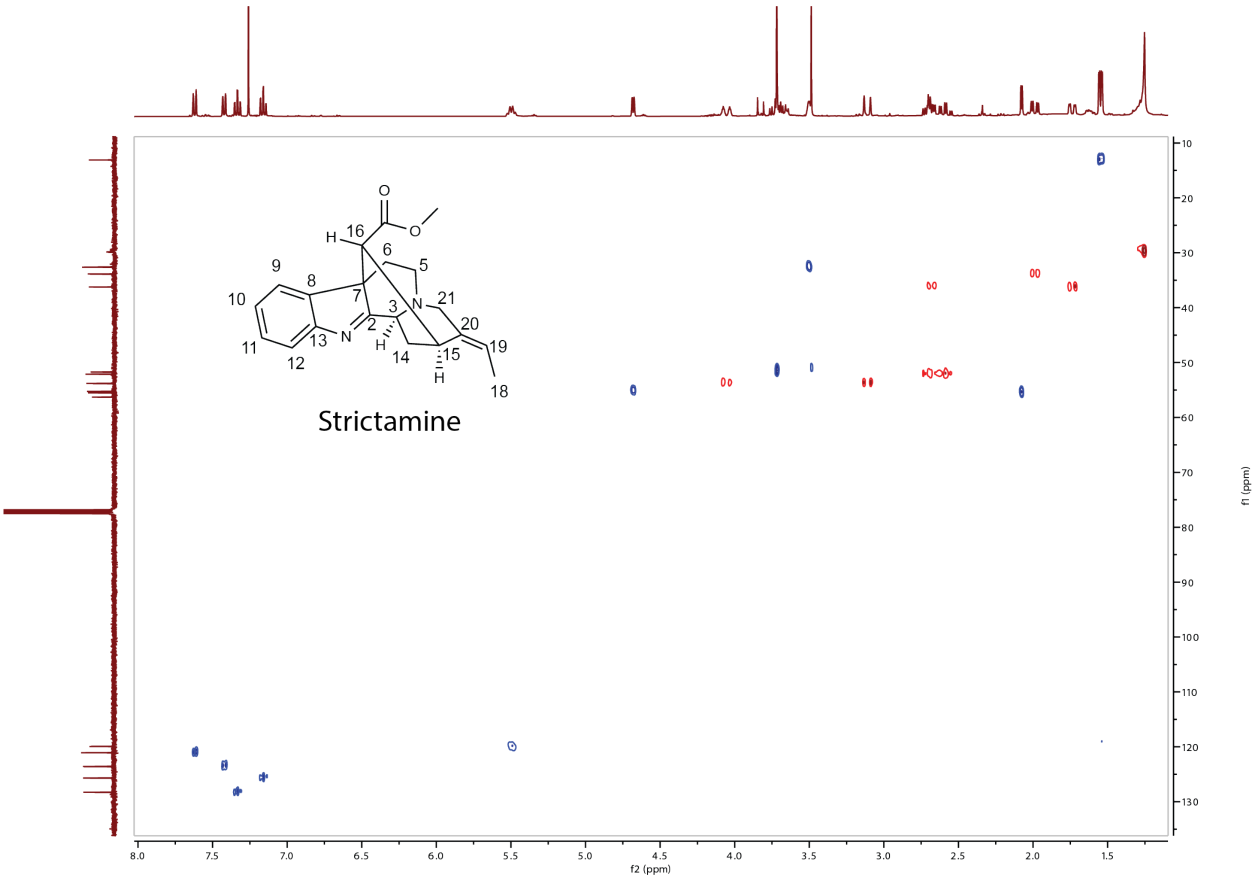

**Supplementary figure 31. HSQC NMR spectra of strictamine in $CDCl_3$.**

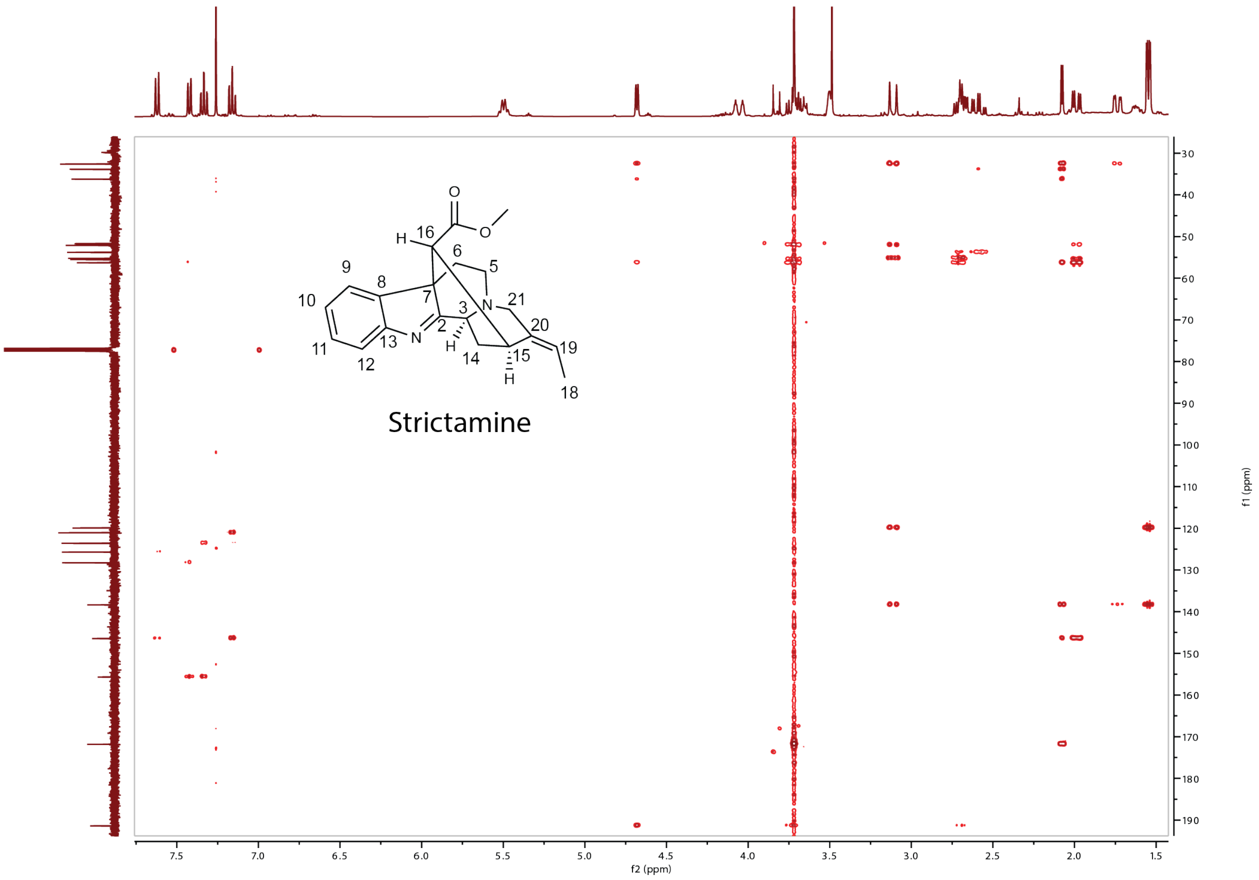

**Supplementary figure 32. HMBC NMR spectra of strictamine in $CDCl_3$.**

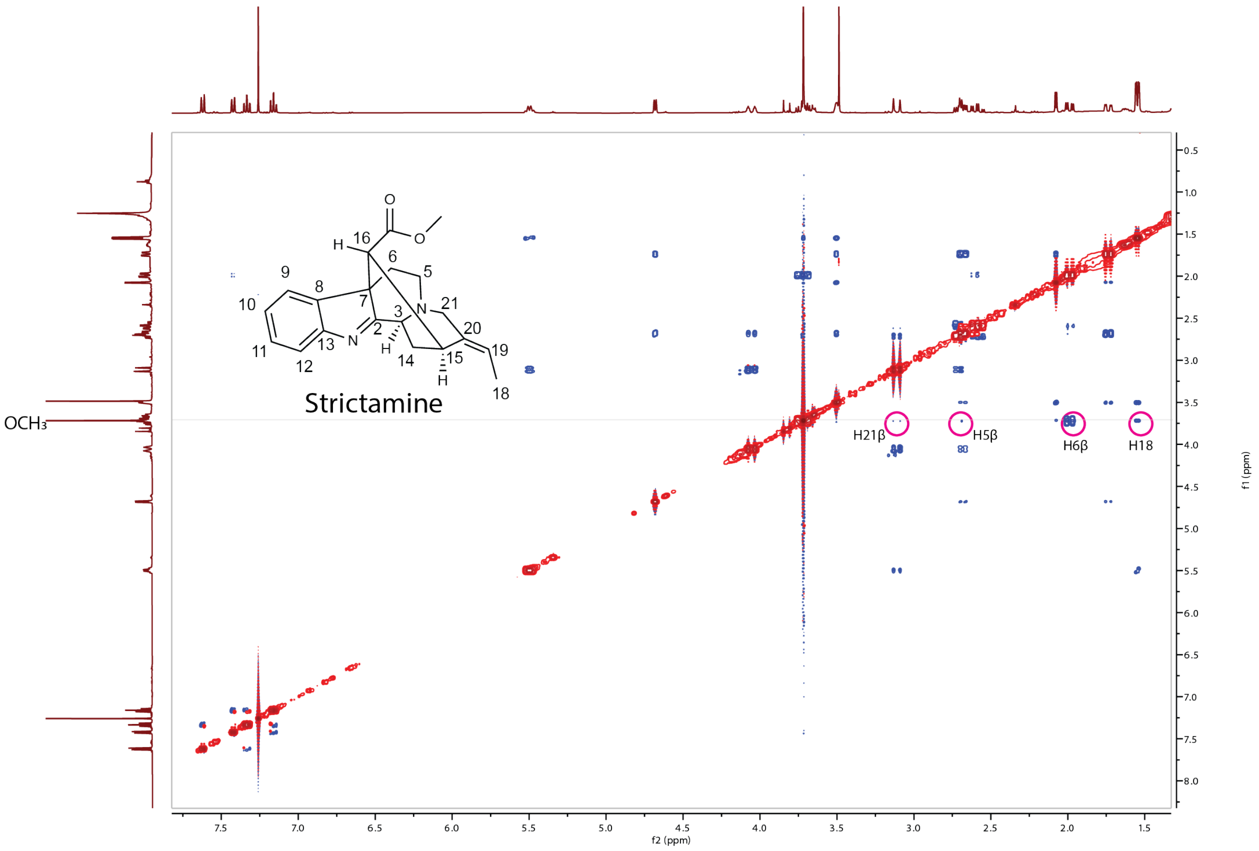

**Supplementary figure 33. NOESY NMR spectra of strictamine in $CDCl_3$.**

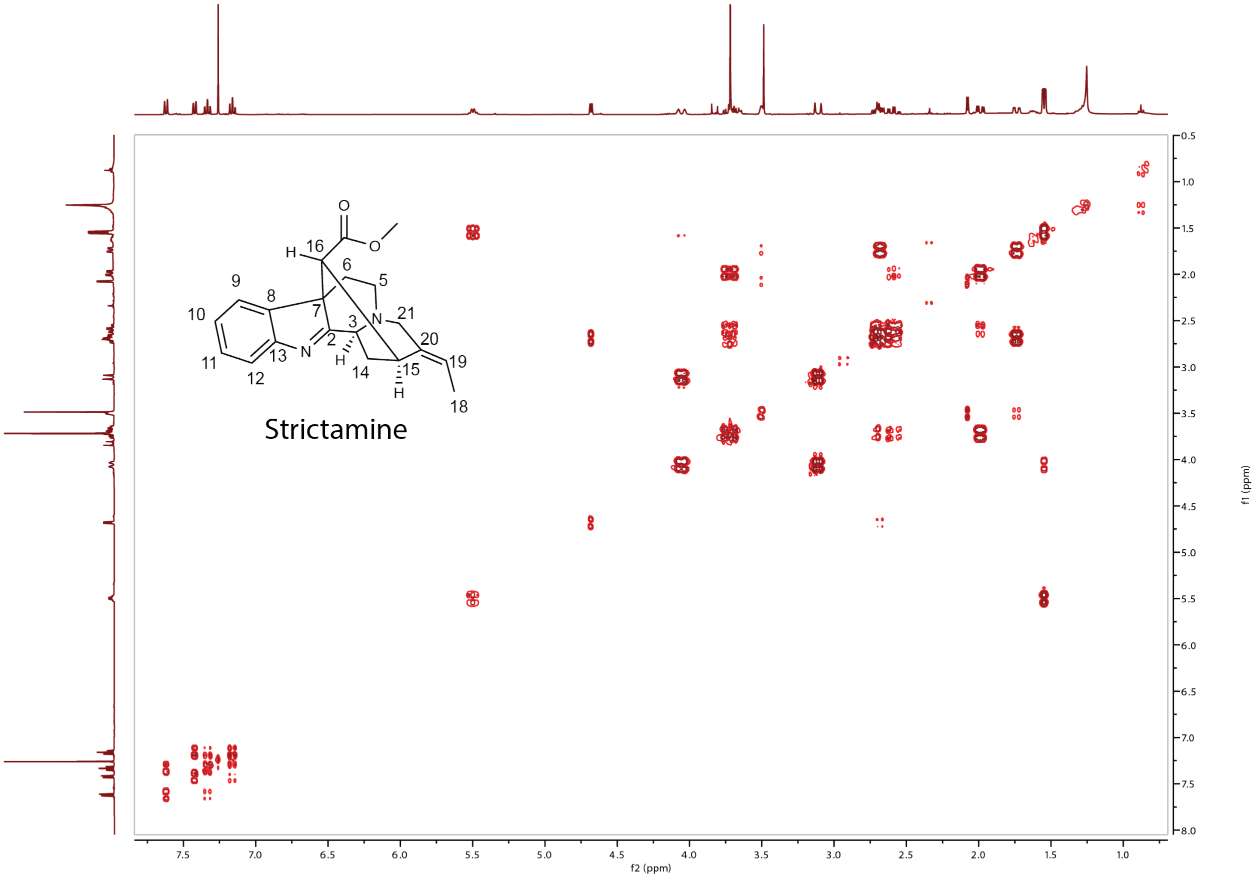

**Supplementary figure 34. COSY spectra of strictamine in $CDCl_3$.**

### Supplementary table 1. cDNA sequences of new enzymes in this study.

| | |
|---|---|
| CrSBE | ATGGATGAGATGATGAACTTCTCCCTCACCTCTCCCATTTCCTTTCTCCTCTCCTTACTTCTTATCCTATTTCTTATCCTATTTATGATGAAAGGAAACCAAACCCACAATGGCA<br>AAAAATTACCACCGGGTCCCAAAAAACTTCCTATTATAGGAAACCTCCATCAAATGATAGGTTCACATCCTCATCGTGTTCTAAAAAAATTGGCCGATAAATATGGACCTATTAT<br>GCACTTGCAAATGGGCCAACGATCAGCCGTTGTTATATCATCAGCGGAAAAAGCAAAAGAAATATTGAACATTTATGGTGTTCAAGTTGCTGATAGACCTCAACTCATCGTATCT<br>AAAATTATGTTGTACAATAGTTTGGGTGTCACTTTTGCTCCTTATGGTGACTACTTAAAGCAATTGCGTCAAATTTATGCCGTAGAACTTCTTAGCCCTAAAACAGTTAGGTCCT<br>TTTGGACAATTATGGAAGATGAAGTTTCCACAATGGTTAACTCAATTAAATCTGAAATTGGTCAACCAATAATTTTGCATGATAAAATGATGACTTATTTGTATACTATGCTTTG<br>TAGAGTTACATTTGGTGGCGTATGTAATGGACGCGAGACACTAATAATGGCAGCTAAAGAAACGTCAGCGCTTTCTGCTGCTATTAGGATTGAGGATTTGTTTCCTTCAGTGAAA<br>ATACTTCCTTTAATTAGTGGATTAAAGTCAAGATTAAACAATTTGTTGAAAACACTTGATACCATTCTTGAGGATATTATCAGTGTTCGTGAGAAGAAATTATTAACTCAGCCAT<br>TGTTGGATGATGAAGATATGTTGGGAGTTCTCCTCAAGTACAAAAATGGAAAGGGAAAAGATACTAAATTTAGAGTCACCAACAACGACATCAAGGCAGTCATTTTTGAAATTAT<br>CTTAGCTGGCACCCTAAGTTCATCAGCTATAGTAGAATGGTGTATGTCTGAAATGATAAAAAACTCTGAAAAGTTTAAAAAGGCACAAGATGAAGTTAGGAAAGTTTTAAAGGGT<br>AAAAAAACAATTAGTGGAAGTGATGTTGGTAAAATGGAATATGTTAAAATGGTGGTTAAGGAATCTTTGAGATTACACCCTCCTGCTCCCATATTGTTTCCAAGAGAATGTAGGG<br>AAGAATTTGAGATAGATGGAATGACTATACCTAAAAAAACTTGGTTGATTGTAAACTACTGGCCAATAGGAAGAGATCCCAGACTTTGGGACGACGCCGACAAGTTTGAGCCGGA<br>GAGGTTCAGTAATAGCAGCATTGATTTCAATGGAAGCCATTTTGAGTTGATACCATTTGGTGCTGGAAGAAGGATTTGTCCTGGAATATTACTTGGAACAACAAATGTTGAGCTT<br>TTACTTGCTACATTTCTCTATTATTTTGATTGGAAACTTCCTCAAGGTATGAAACCCGAAGAATTAGATATGAATGAGGTATTTGGTGCTGGTTGCATAAGGGAAAGTCCATTGT<br>GTCTCATTCCTAGCATCTCATCAGCAGTTGAAGGAAATTAA |
| TeSBE | ATGGAGATGATGAACTTCTCTTTCACCTCCCCAGTTTTCCTTTTCCTCTCCTTCTTTCTTCTCATGTTACTCAAACAACTGAGAGGAAACAAAAGTCACAAGGGCAAAAAATTGC<br>CGCCAGGTCCTAAAAAACTTCCCATCATTGGAAATTTGCACCAGATGGTAGGATCACTGCCTCATCGTGTTCTCAAAAAATTGTCAGATCAATATGGACCGGTCATGCACTTGCA<br>GATCGGTGAAATTTCAGCAGTAATAATATCGTCTGCCGAAAAAGCAAAAGAAGTCTTGAACACTCATGGCGTCGCGGTTGCAGATCGGCCCCAAACTATTGTTGCTGAGATCATG<br>TTGTATAATAGTTTGGGTGTCACTTTCGCTGCATATGGCGAATACTTGAAGCAATTGCGTCAAATTTATGCATTGGAACTTCTTGGCCAGAAAAGTGTTCGATCCTTCTCGACTA<br>TCATGGAAGATGAGCTTTCGACTATGGTTGCATCAATCAAGTCTGAAGCAGGGCAACCGGTGATTTTGAATGATAAAATGATGACTTATTTGTATGCCACACTTTGCAGAGCAAC<br>TGTTGGTGGTATATGCAATGGACGTGAGACATTGATAATGGCAGCAAAAGAAACTTCAGCGCTTTCTGCATCCATTAGGATTGAAGATCTCTTCCCTTCGGTGAAAATACTTCCC<br>ATTGTCACTGGACTAAAGTCCAGATTAACCAAACTGTTGAAACAACTTGATAGCGTTCTTGAGGATATAATCAGTGAGCGTGAGAAGAAATTATTCTCATCCAACGACAACCAGC<br>CATTGTTGGATGAAGAAGATATGCTGGGCGTTCTCCTGATGTACAAGAGTGGAAAGGGAAAAGGCTCTAAATTCCGAATCACCAATAATGACGTCAAGGCTATAATTTTTGAATT<br>GTTCTTGGCTGGAACTCTAAGTTCATCAGCCATAGTTGAATGGTGCATGTCTGAAATGATTAAAAACCCTATAGTTCTGAAGAAGGCACAGGATGAAGTGCGGCATGTTCTCAAG<br>GGTAAGCAAACAGTCAGCGGAAGTGATCTTGCGAAATTGCAGTACGTGAAAATGGTGGTTAAGGAATCTATGAGATTACACCCTCCTGCTCCCTTATTGTTTCCAAGAGAATGCA<br>GGGAAGCATTTGAGATGGACGGGATGACCATACCAAAAAAATCTCGGGTGATCATAAATTATTGGGCAGTGGCAAGAGATCCTGAACTTTGGCATGATGCCGACAAGTTTGAGCC<br>GGAGAGATTCAGCAATAGTACCGTTGATTTCTATGGAAGCCATTTTGAGTTGATACCATTTGGTGCTGGAAGGAGGATTTGTCCAGGCATATTATTTGGAGTAACTAATGTTGAG<br>CTTTTGCTTGCTGCATTTCTCTATCATTTTGATTGGCAAATTCCTGGAGGCATGAAACCTGAAGAATTAGATATGACAGAGTTATTTGGTGCTGGTTGCATAAGGGAAAATCCAT<br>TATGCCTAATTCCCAAGATCTTCATTTGA |
| VmSBE | ATGGAGATGATGAACTTTTCTCTCACATCCCCAGCTTTCCTTCTCTTCTCTTTCTTTCTTCTGATGCTACTTTGGCAATTGAAACCTAAGAAGGGCAACAAAAGATTGCCACCTG<br>GTCCAAACAAACTTCCCATAATTGGAAATCTCCACCAGATGATGGGCTCACTTCCTCATCGTGTCCTCAAGAATTTAGCTGACAAATATGGACCAGTCATGCACCTGCAAATCGG<br>TGAACTTTCAGCTGTGATCGTTTCATCAGCTGATAAAGCTAAAGAAGTCTTGAACACTCATGGCGTTCAGGTCGCAGACAGGCCCCAGACAACCGTGGCTAAGTTAATGTTGTAC<br>AGTAGTTTGGGAGCGACTTTCGCACCATACGGCGATTACTTGAAGCAATTGCGTCAAATTTATGCAGTTGAACTTCTTAGTCAAAAAAGTGTAAAATCATTTTGGAATATAATGG<br>AAGACGAGCTTTCGACTTTTGTTACATCAATAAAGTCCGAAGTCGGACAACCTATGATTTTGCATGAAAAAATGATGACTTATTTGTATTCCACACTGTGCAGAACAACGGTTGG<br>TAGTGTATGCAGTGGAAGAGAGACATTAATTATGGCAGCTAGAGAAACTTCTGCTCTTTCAGCTGCCATTAGGGTCGAAGATCTTTTTCCTTCAGTGAAATTCCTTCCTGTTATT<br>AGTGGTTTAAAATCTAGATTAAGCAACCTTTCGAAAAAACTTGATGCTGTTCTTGAGGATATAATCAGCCAACGTGAAAAGACAATGTTATCCTCCGACTATGAGCCATCCTTGG<br>ATGAAGAAGATATGCTTGGTGTTCTCCTTTTGTACAAGAATGGAAAGGGAAAAGACACTAAATTCCGCATCACCAATAATGACATCAAGGCTATCATTTTCGAACTGATATTGGC<br>TGGAACTCTAAGTTCAGCAGCAATAGTAGAATGGTGCATGTCTTTACTGATGAAAAACCCAAGTGCTTTAAACAAGGCGAAGGATGAAGTGAGACAAGTTTTCAAGGACAAGAAA<br>AACATAACCGGAAGCGAAGTCTCAAAATTACAATACCTTAAAAATGTAATTAAGGAAGCTGTAAGATTACACCCACCTGCTCCATTATTATTCCCAAGAGAATCAAGAGAAGAAT<br>TTGAAATGGATGGAATGATAATACCAAAAAAATCTTGGATTATTATAAATTATTGGGCAGTTGGAAGAGATCCAAAGGTTTGGGAAGAAGCAGATAAGTTTAATCCAGAGAGGTT<br>TAACAATAGTGCAATTGATTTCTATGGAAGCCATTGTGAGTTGATACCATTTGGTGCTGGAAGAAGGGTTTGTCCAGGCATACTATATGGAGTAACTAATGTTGAGCTTTTGCTA<br>TCTGCATTTCTATATCATTTTGATTGGGAACTTCCTGGTGGTATGAAACCTGATGAATTGGATATGGATGAGTTATTTGGTGCTGGTTGCATTAGGGCTAATCCATTGCGTCTCA<br>TCCCAACCATCTCCGATGCTGAACAAGAGTAA |
| VmRHS | ATGGAAACTAAACACATCTTCTCTTCCTCCTCAGTTTTCTACTTGTTCATCAGCTTCATTGCTTTCCTAATAGTAGTGAAGCGACTCTTAAAACCTAAAAGCCATAAGAAATTGC<br>CTCCAGGTCCATGGACCTTACCTTTCATTGGCAATTTACACCAACTTACTGGATCATTGCCACATCGAATCCTCAAAAATATGTCCGATAAACACGGACCATTAATGCGAATAAT<br>GATGGGCGAGAGAACAACTATAATCGTATCATCCGCAAGAATGGCAAAGCTAGTGTTACACACAAATGGCCTCTCTGTGGCCAACAGACCAATTAACAGCGTAGCTGAAATCATT<br>TGTTACAACAATTTAGGGACTACTTTTGCGAAATACGGGGAATATTTAAAACAATTGCGTCAACTTTATATGTTAGAACTTTTGAGTACAAAAAGAGTGAAATCATTTTGGGGTA<br>TTTTTGAAAATGAGTTAGAAAAATTTGTTGGGTCAATTGAGTTAGAGGTTGAAAAACCACTGACTATTTATTGGAAATCAACTGAGTATTTATATGCAGCAATTTGTAGAGTTAT<br>GTTTGGAAGCGTTTGTAGTGAAAGGGATAAACTAATTAAAGTTTGTAAAAAAGTATCAAAATTATCTGCCGCTCCCATTAGGTTTGAAGATCTATTCCCTTCAATTAAAGGTCTT<br>TCTTTTCTTACTGGAACAGACTCAGTTTTAAAAGGTCTTGTCAAAGATCTTGATGATGTTCTTGATAAATTAATTTCTGAGCGAGAAAACAGCTGTTCCACACTGGACGAAGGTG<br>ATATGCTTGGCATTCTCTTGAAGATTAAGAAGGGAGATTTGTACAACTCTAAACTCCAAATCACAAACGACGATATCAAAGCTATCATTTTCGAACTGATGTTGGCTGCGACGCT<br>GAGTGTAGCAGATGTTGTAGAATGGGCGTTTGTTGAAATAATAAGATACCCAGAAGTCTTAAGGCAAGTAAAGGAAGAACTTAGACAAGTATTAAAGGGGAAGAAGACGATAACC<br>GGAGACGATTTAGAGAAGATGGAATATCTACACATGTGTGTTAAGGAATCAACAAGATTACACCCAGCTGCACCCCTTCTTTTCCCAAGAGAAGCAAGAGAAGAGTTCCATGTTG<br>ACGAAGAATATACGATCCCTAAAGGTTCATGGATTATGACTAATTATTGGGCAATCGGAAGGGACCCCGAAATTTGGCCTAAACCTGACGAGTTTGATCCTGAAAGATTTAGAAA<br>CAGTAACATGGATTTCTATGGAAATAGCTTTGAGTTGATTCCATTTGGGGCTGGAAGAAGGGGATGTCCTGGGATCTTATTCGGAATAACCGAAGCCGAGTTCTTGCTTGCTGGT<br>TTGTTCTATCATTTTGACTGGGAACTTCCTGGTGGGATTACACCCAAAGAGATTGATATGACTGAAGTTTTTGGTGCTGGTTGTATTCTCAAAAACCCATTAAGTGTCATTCCAG<br>TTCTTGCTGATAATTGA |
| AtaRHS | ATGATGCAGTTCTCTTTGTCCTCCCCAGTTGTATACATTCTCATTTTAATTGCTTTCCTGATAGTGAAGCAACTAGTAAAACCCAAAAGTCACAAGAAATTGCCTCCAGGTCCAT<br>GGACGTTACCCCTCATTGGCAATCTGCATCAGATTTTAGGTGCATTGCCACATCGAATCCTCAAGAAACTGTCCGATCACCATGGACCTTTGATGCGAATAATGATGGGTGAGCG<br>GACAACAATAATTGTATCATCCGCTAGGATGGCAAAACTAGTGCTGCACACACAGGGTCTAGCAGTGGCCAACAGGCCAATTAACAGTGTGGCGAAAATCATCTGCTACAACAAT<br>CTGGGAACTACTTTTGCCAAATATGGCGAGTACTTGAAGCAATTGCGTCAACTTTATATGTTGGAACTCCTGAGTCCCAAAAGAGTTCAATCATTTTGGGTTATTTTTGAAGATG<br>AGCTCGAGACTTTTGTTAAGTCGATCAAGTCCGACGTGGGACAACCAATGGTTATATACGAGAAATCGACTGCTTACTTGTATGACACAATTTGCAGAGTGATGCTTGGAAATGT<br>CTGCAATGAGAGAGAGAAACTGATCAAGGTATGCAAAAAAGTATCAAAACTTTCTGCAGCCCCCATTAGGGTTGAAGATCTCTTCCCTTCAATGAAATTCATTCCGTTGCTGACT<br>GGAACAAACTCCGTTTTAAAAGGTCTGGTAAAAGACCTCGACGATGTCCTTGATATCGTAATTGCTGAGCGTGAAAATAGCCAAGACGCAGTGAAGGAGGAAGATATGCTCAGCA<br>TTCTCTTGAAGCATAAGAGTGGTGACGGCAATAACTCAAAACTCCAAATCACGAACGACGACATCAAAGCTATCTTTTTTGAATTGCTGTTAGCTGCAACTCTGAGCGTAGCGGA<br>CGTTGTTGAATGGGCTATGGTAGAAATACTGAAGCATCCTAACGTTTTGAAGCAGGTACAAGATGAACTAAGGCAAGTGTTGAAGGGAAAGAAAAGAATCACTGGAAGCGACATC<br>AACAATTTGAAATATCTACACATGTGTGTTAAGGAATCTACGAGATTACACCCTGCTGCTCCCCTCTTATTCCCAAGAGAAGCCAGAGAAGAATTTGAGATTGATGGATATACAA<br>TTCCCGAAGGAGCATGGGTGTTGACAAATTACTGGGCAATCGGAAGAGACCCTGAACTTTGGCCAAAGCCCGACGAGTTTGATCCAGAGAGATTCAGAAATAGTAGCATCGATTT<br>CTATGGAAACCACTTTGAGTTAATTCCTTTTGGTACCGGAAGAAGAGGATGTCCAGGCATATTATTTGGAATAACAGAAGCTGAGTTCTTGCTAGCTGCAATGTTCTATCACTTT<br>GACTGGAAACTTGCAGGTGGGATCACACCTGCGGAGATAGACATGACTGAGGTATTCGGTGCTGGTTGCATTCTCAAAAATCCTTTGAATCTGATACCATATATTGCAGACAATT<br>GA |
| AtaAAR | ATGGAAAAGCAAGTGAAGATCCCGGAGATAGAGTTGAACTCAGGCCACAAGATGCCGCTGGTGGGCTTCGGGACATGTGTGCCGGATCCCATTCCACCATTAGAGGAACTTGCTG<br>CAATCTTTTTGGAAGCTATTAAGGTGGGTTATCGCCACTTTGACACAGCATCAAGCTATGGCACAGAGGAAGCCCTCGGTAAAGCTGTTGCTCAAGCAATAGAGAATGGCTTGGT<br>CAAGGGTAGAGAAGAGTTCTTTATCACTTCTAAATTGTGGTGTGAAGATGCTGATCATGATCTTATCTTGCCTGCCCTCAAGAAAACACTTGGGAATCTGGGGCTTGATTATTTG<br>GATCTTTATATGATTCACATGCCAGTGAGAGTGAGGAAGGGTGCTCCAATGTTTAATTATTCAAAAGATGACTTACTCCCTTTTGACATACAAGGGACATGGAAAGCCATGGAAG<br>AGTGCAGCAAATTGGGCTTGTCTAAGTCTATTGGTGTAAGCAACTACACTTGTGAAAAACTCTCAAAACTGCTCGAAAATGCCACCATCCCCCCAGCAGTCAATCAGGTGGAAAT<br>GAATGTTTCTTGGCAACAGAGGAAATTACTGCCATTTTGCAAAGAGAAAAACATCCATGTTACAGCATGGTCTCCTCTTTTGTCTTATGGTTCCGCTTGGGGCAGTAATGCAGTG<br>ATGGAGAACCCAGTCCTTGTAAATATTGCAGCTTCCAAAAGCAAAACGGTGGCACAGGTTGCACTAAGATGGATATACGAACAAGGAGCAAGTTTCATTATGAGGACATTCAACA<br>AGGAAAGGATGTTCCAAAATGTTCAGATTTTCGATTGGGAACTCTCTAAAGAAGAATTGGATCAAATTCAACAAATCCCTCAACGTAGAGGCACTCTAGGAGAAGATTTTATCAA<br>TCCAGAGGGACCAATCAAGTCGGTAGAGGAACTTTGGGATGGAGACTTGTAA |
| AtaAAR<br>-like | ATGGACAGAATAGCGGACATCCCAGAAACAATGTTGAATTCGGGCCACAAGATGCCACTGGTGGGCTTGGGATGTGCAGCGCATCCCACTCCACCATTAGAAGAACTAACAGCAA<br>TTTTATTGGAAGCCATCAGAATTGGTTATCGCCACTTTGACACCGCAGCATCCTATGGCACTGAGGAAGCCCTTGGTAGAGCTGTCTCTCAAGCAATACAGACTGGTTTGGTTAA<br>GGGAAGGGAGGAATTGTTCATCACTTCTAAATTGTGGTGCATAGATGCAGATCATGACCTGGTCATGCCTGCCCTCAAGAAAACACTTGGGAAGCTCGGACTGGTTTATTTGGAT<br>CTTTATCTGGTTCACTGGCCCGTAAGAATGAAGCAAGACGTTACCAATTTTTTCAAGTTGTCAAAGGATGACCTCCTCCCTTTCGATATGCACGGGACATGGAAGGCCATGGAAG<br>AGTGTAAGAAATTGGGTTTGGCCAAGTCTATTGGTGTAAGCAACTTTACTTGTGAAAAGCTCTCAAAACTCCTAGAAAATGCCACAATTTGCCCATCAGTTAATCAGGTGGAGAT<br>GAATGTGGCGTGGCAACAGAGAAAATTGCTGCCGTTTTGCCTGCAGAAAGGAATTCATATCAGTGCATGGTCTCCATTGGGAGCATATGGTAACTTCTGGGGCTCTAATGCAGTA<br>ATGGAGAATCCAATCCTCAAAGAAATTGCATCTTCCAGAAGCAAGACTGTGGCACAGGTTGCGCTAAGATGGATATATGAACAAGGAGTAAGTGCAATAGTGAAGAGCTTCAATA<br>AGGAGAGGATGATTCAAAATCTACAACTTTTCGATTGGGAATTCACTAAGGAAGAATTGGATCGAATTCAAGAACTCCCTCAATGCAGGGGTTTTTCAGGAGATATGTTTATCCA<br>TCCGGAGGGGCTAATCAAATCAGCGGAGGAACTCTGGGATGGAGATATTTGA |

### Supplementary table 2. Primers used in this study

| # | primer | sequence 5'-3' |
|---|---|---|
| 1 | CrSBE-BamHI-F | CTAAGGATCCATGGATGAGATGATGAACTTCTCCC |
| 2 | CrSBE-SalI-R | CGGTGTCGACATTTCCTTCAACTGCTGATGAT |
| 3 | VmSBE-BamHI-F | ATAGGATCCATGGAGATGATGAACTTTTCTCTCACATCC |
| 4 | VmSBE-SalI-R | ATAGTCGACCTCTTGTTCAGCATCGGAGATGGT |
| 5 | Attb-VmRHS-F | GGGGACAAGTTTGTACAAAAAAGCAGGCTTCATGGAAACTAAACACATCTTCTCTT |
| 6 | Attb-VmRHS-R | GGGGACCACTTTGTACAAGAAAGCTGGGTCATTATCAGCAAGAACTGGAATGACA |
| 7 | AsRHS-BamHI-F | ATAGGATCCATGGAAATGATGCAGCTCTCTT |
| 8 | AsRHS-SalI-R | GACGTCGACATTTTCTGCGAGACGTGGTATGA |
| 9 | AtaRHS-BamHI-F | gtcaaggagaaaaaaccccggatccATGATGCAGTTCTCTTTGTCCT |
| 10 | AtaRHS-SalI-R | aaatcaacttctgttccatgtcgacAATTGTCTGCAATATATGGTATCAGA |
| 11 | AtaAAR-BamHI-F | acaaggccatggcgatatcggatccAATGGAAAAGCAAGTGAAGATCC |
| 12 | AtaAAR-SalI-R | cgagtgcggccgcaagcttgtcgacTTACAAGTCTCCATCCCAAA |
| 13 | RsPNAE-BamHI-F | ATAGGATCCCATGGATTCTGCTGCAAACGC |
| 14 | RsPNAE-SalI-R | GAAGTCGACTTATGAATCTGATATATCAAGCAGGCA |
| 15 | VIGS-CrSBE-F | TCAGGAATTCAAGAGATCCCAGACTTTGGGACGA |
| 16 | VIGS-CrSBE-R | TCAGGAATTCTCCCTTATGCAACCAGCACCAA |
| 17 | VIGS-CrGO-F | ATCTCTAGACCATCTTCAGTCTCCTCTCTTG |
| 18 | VIGS-CrGO-R | ATTCTCGAGTCTCCATACTGAGCAAAGGT |
